# Field–Flow–Front mapping of breast cancer evolutionary dynamics in situ

**DOI:** 10.64898/2026.06.23.733638

**Authors:** Yuhong Zhang, Juan Ji, Mingquan Gao, Yiyao Zhang, Zifei Wu, Chuan Wu, Junchao Wang, Hongyuan Jia, Yiyan Yang, Luo Liang, Shuang Li, Yajie Tu, Lan Lei, Yaxing Pei, Hong Yang, Shenglin Luo, Yang Liu, Rong Li, Junjie Li, Weidong Wang

## Abstract

Tumour progression reflects not only which clones arise but where, when and in what context they expand—dimensions genotype-centred reconstructions leave unresolved. Here we reconstruct breast cancer evolution in situ across 34 Visium HD, 16 Xenium, 32 MIBI-TOF and 10 CODEX samples, serial-section 3D reconstruction, single-cell and bulk transcriptomes, and genome-wide CRISPR dependency profiles. We partition tumours into 296 cancer microzones—stroma-bounded units within which expansion is reconstructed—and define in each a Cancer Progression Metric(CPM) coupling a transcriptomic clock to expansion geometry. Projected onto tissue, CPM yields a Field–Flow–Front model rendering progression as a continuous physical process and resolving subclonal architecture into spatially coherent domains rather than predefined branches. Unexpectedly, the most advanced fronts were not the most proliferative but low-dependency, slow-cycling populations with directional expansion, driven by an extracellular matrix programme whose evolutionary force exceeded inflammation by nearly an order of magnitude yet whose genes were the least cell-autonomous. Under chemotherapy it persisted while its clonal carriers reshuffled, marking a transferable, stroma-coupled front process—not a fixed clone—as the unit of advance. Distilled into an evolutionary advantage load, this front process predicted recurrence and survival across independent cohorts and improved on conventional staging, establishing a tissue-embedded paradigm for mapping tumour evolutionary dynamics, from local fronts to patient outcome.

## Introduction

Tumour growth, invasion, and therapeutic resistance are fundamentally ecological and evolutionary processes.^1^ Since the molecular clock^2^ and clonal evolution theory,^3^ high-throughput sequencing has greatly accelerated the reconstruction of tumour history,^4, 5^ particularly in subclonal inference.^6–14^ Yet a central obstacle remains: most approaches follow a genotype-first paradigm,^15^ inferring evolutionary lineages largely apart from the spatial tissue context in which selection, competition, and expansion actually occur.^16, 17^

Although effective for lineage tracing through mutation frequencies, copy number variation,^18–21^ allelic imbalance,^10^ and joint genome–transcriptome profiling,^22^ abstracting away native tissue architecture gives rise to the “clonal illusion”,^17^ in which inferred subclones fail to map coherently onto histological structures or functional states. Purely genetic models thus struggle to capture phenotypic plasticity and the non-tree-like spatial constraints of the tumour microenvironment,^17, 23^ often overestimating diversity, generating spatially fragmented subclones,^10^ or treating space only as a downstream annotation.^21, 24^

We therefore pursued a two-stage strategy that establishes an organizational substrate before operationalizing spatiotemporal coupling. Because existing spatial pseudotime methods couple sparse gene expression directly against dense spatial coordinates at the level of individual cells or bins, ordering remains unstable and directionally indeterminate unless inference is first bounded within biologically coherent units;^25–27^ we accordingly delineated cancer microzones (CMZs)—stroma-encapsulated, histologically bounded regions within which cells can be reasonably modelled as a clonal expansion from a common ancestor and whose boundaries define a physically coherent expansion geometry. Conceptually, CMZs define the concrete structural space in which tumour evolutionary dynamics can be inferred, converting histological architecture from a passive spatial backdrop into the mesoscopic domain where selection, clonal expansion and invasion are organized. This pre-specification grounds inference in the physical habitats where selection and invasion occur and constrains local trajectories to clonally coherent populations, stabilizing molecular ordering that is otherwise unreliable under sparse signals. Cells remain the fundamental units of evolution, whereas CMZ-bounded populations define the mesoscopic units within which it unfolds. We then introduce the Cancer Progression Metric (CPM), an evolutionary encoder mapping transcriptomic variation and CMZ-defined spatial topology onto a single continuous coordinate: pseudotime encodes the evolutionary content of molecular adaptation, while a direction-normalized spatial distance anchors trajectory orientation against each CMZ’s expansion geometry. Through this dual-clock design, CPM invokes the ecological principle of space-for-time substitution, overcoming the conventional separation between phylogenetic reconstruction and spatial context and resolving tumour evolution as a computable, spatially organized process in situ (Fig. 1).

**Fig. 1:**
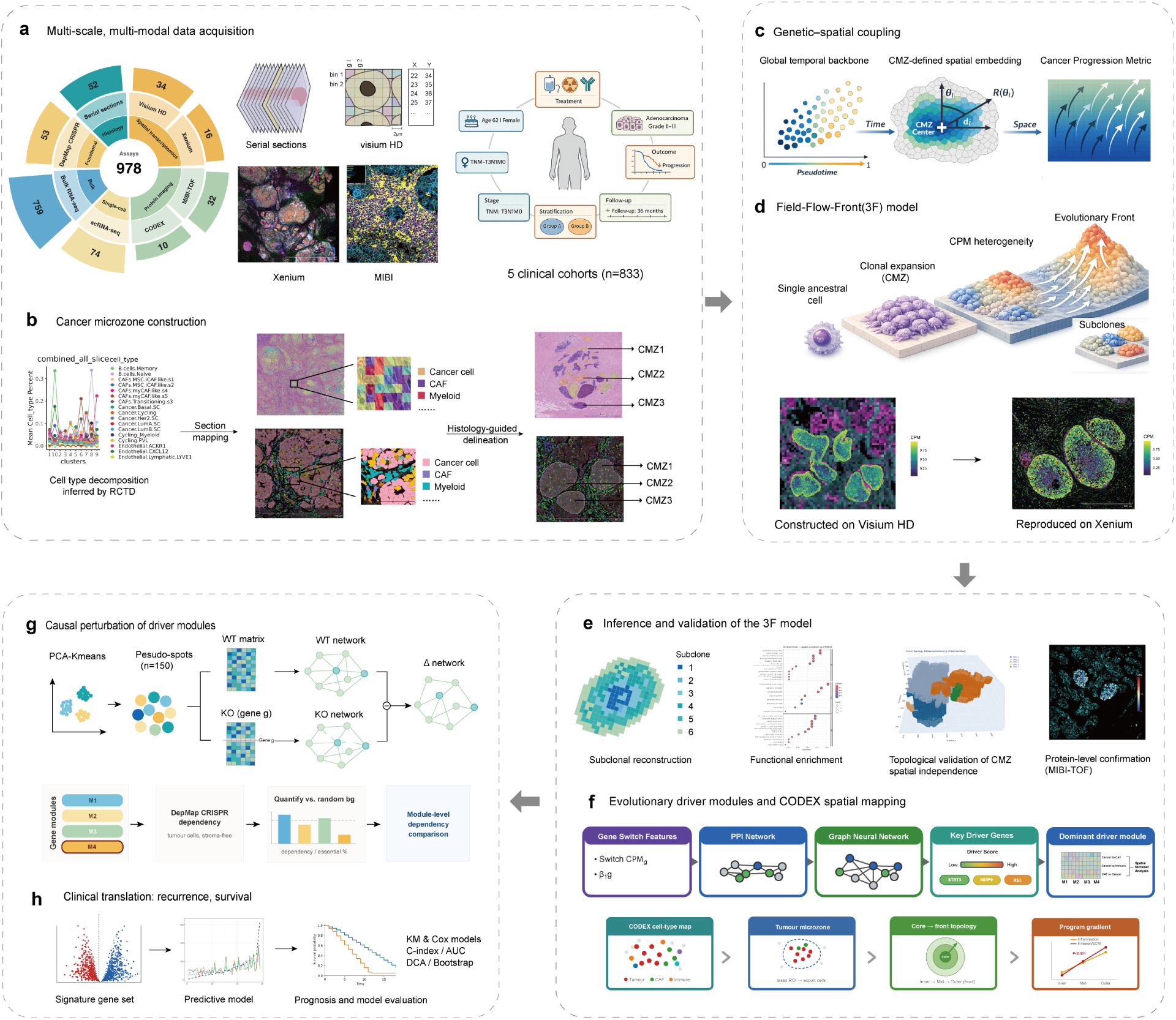
A tissue-embedded cartographic framework for decoding breast cancer evolution through Field–Flow–Front dynamics. **a,** Building the data foundation: multi-scale tissue profiling. The central value (978) is the total platform assays across all modalities (excluding the 52 serial sections). Representative images of serial sectioning, Visium HD, Xenium and MIBI are shown. The clinical cohorts comprise 833 patients used for downstream cohort-level analyses. **b,** Construction of cancer microzones (CMZs). Tumour cells inferred by RCTD-based cell-type decomposition were mapped back to tissue sections, and histology-guided delineation was used to define spatially adjacent, boundary-delimited tumour aggregates as CMZs. **c,** Genetic–spatial coupling and construction of the Cancer Progression Metric (CPM). Global pseudotime provides the temporal backbone, while CMZ-defined spatial embedding captures direction-normalized spatial progression, together yielding CPM as a unified spatiotemporal metric of tumour progression. **d,** Field–Flow–Front (3F) model. CPM is projected into tissue space to define an evolutionary field, directional flow and progression fronts, capturing tumour evolution as a continuous spatial process. The model was constructed on Visium HD data (left) and validated on an independent Xenium dataset (right), with consistent field–flow–front patterns confirming its generalizability. **e,** Inference and validation of the 3F model. Subclonal architecture was reconstructed and characterized by functional enrichment, while the spatial independence of CMZs was assessed by 3D topological reconstruction. Model predictions were further corroborated at the protein level using MIBI-TOF multiplex imaging. **f,** Evolutionary driver modules and spatial validation. GeneSwitch features, PPI networks and graph-based modelling prioritized key driver genes and dominant evolutionary modules, which were then spatially validated at the protein level using independent CODEX multiplex imaging. **g,** Causal perturbation of driver modules. Top, in silico per-gene knockout of pseudo-spots yields a differential (Δ) co-expression network for each module. Bottom, DepMap CRISPR profiles validate intrinsic dependency across modules against a random-gene background. **h,** Clinical translation and generalization. Spatially derived advantage signatures were projected into predictive models and evaluated for prognostic utility, enabling recurrence and survival prediction across independent patient cohorts.

Using this framework, we show that intratumoural heterogeneity is organized into physically segregated CMZs that serve as mesoscopic units for reconstructing evolution within native tissue, elevating spatial organization from a downstream annotation to the primary scaffold for evolutionary inference. Within this architecture, the CPM and the Field-Flow-Front(3F) model resolve tumour evolution as a continuous spatiotemporal process rather than a static lineage summary, shifting its representation from a phylogenetic tree to a computable map extending from proliferative cores to invasive fronts. Across this map, progression converges on a low-dependency, high-invasion front, and we identify a conserved, stromal-coupled ECM-driven programme preserved under chemotherapeutic selection while its clonal carriers are reshuffled—supporting the biological validity of the 3F model. Finally, projecting local front dynamics to the patient level yields an evolutionary advantage load (E-load) associated with recurrence and survival in independent breast cancer cohorts. Together, these findings establish a tissue-embedded paradigm for mapping tumour evolutionary dynamics, from local fronts to patient outcome.

## Result

### Cancer microzones as mesoscopic units for reconstructing tumour evolutionary dynamics in situ

To reconstruct tumour evolutionary dynamics directly within native tissue architecture rather than treating spatial organization as a downstream annotation, we first established the structural scale at which these dynamics are physically organized in situ. We generated multi-scale Visium HD spatial transcriptomic data with paired serial H&E sections and combined them with multi-modal external datasets spanning single-cell, bulk, proteomic, histological, functional-genomic (CRISPR dependency) and clinical data (Fig. 1a and Extended Data Fig. 1a–c). The discovery set comprised 34 Visium HD breast cancer samples from 24 patients; cross-platform validation used single-cell-resolution spatial transcriptomics (16 Xenium samples) and protein-level imaging (32 MIBI-TOF and 10 CODEX samples), together with a single-cell reference of 74 scRNA-seq samples, two bulk RNA-seq cohorts (759 samples with survival data) and genome-wide CRISPR dependency profiles from 53 breast cancer cell lines (DepMap) (Supplementary Table 1). Pathological observation that breast cancer predominantly grows as focal microregions^21^ indicated that its evolutionary dynamics are organized not at the level of isolated cells or the tumour as a whole, but within spatially bounded local compartments, defining the appropriate scale at which to reconstruct these dynamics.

Accordingly, we defined individual tumour cells as the minimal units of spatial analysis and delineated spatially adjacent, boundary-delimited tumour-cell aggregates as cancer microzones (CMZs)—the physical units within which local tumour evolution unfolds, and within which cells can be reasonably modelled as a clonal expansion from a common ancestor. Using sub-cellular spatial transcriptomics (10x Genomics Visium HD), cell-type annotation at single-cell resolution and histopathological priors, we delineated CMZs by histology-guided, stroma-bounded criteria (Fig. 1b, Extended Data Fig. 1b and Extended Data Fig. 2b), applying the same approach across Xenium, CODEX and MIBI-TOF platforms to yield 296 CMZs in total. As a physiological baseline, we defined benign microzones (BMZs)—non-malignant epithelial aggregates with topologies analogous to CMZs—within the peritumoral normal breast tissue.

By discretizing the continuous tumour mass into compartmentalized units, our framework establishes CMZs as the mesoscopic units for reconstructing tumour evolutionary dynamics in situ, preserving the physical contexts in which local progression, directional expansion and invasive-front formation occur. Treating CMZs as evolutionary–ecological units reveals that tumour evolution proceeds through parallel, spatially organized trajectories across locally constrained microregions. Tissue architecture is therefore not appended to evolutionary inference but constitutes the coordinate system from which it begins.

### Cancer Progression Metric: a unified spatiotemporal coordinate embedded in tissue space

To resolve tumour evolution within CMZs as a continuous, tissue-embedded process, we first established a biologically grounded ordering of local progression and then constructed the Cancer Progression Metric (CPM) as a unified spatiotemporal coordinate. Rather than relying on arbitrary algorithmic roots, we determined the ancestral root state of each CMZ trajectory using a spatial–size rule: ancestral states occupy the CMZ interior and form larger populations (“inner-early, outer-late” and “large-early, small-late”), whereas derived states expand toward the periphery as smaller clusters (Fig. 2a and Extended Data Fig. 3a). This ordering was independently supported by functional and transcriptomic features: EMT scores increased from root to terminal states while stemness declined,^28^ and differential expression shifted from basal-epithelial and proliferation modules early to ECM remodelling (*COL1A1*, *SPARC*, *FN1*) and context-dependent immune/inflammatory signalling (*B2M*, *C3*, *CXCL9*) late,^29^ validating the inferred direction (Fig. 2a and Extended Data Fig. 3b–c).

**Fig. 2:**
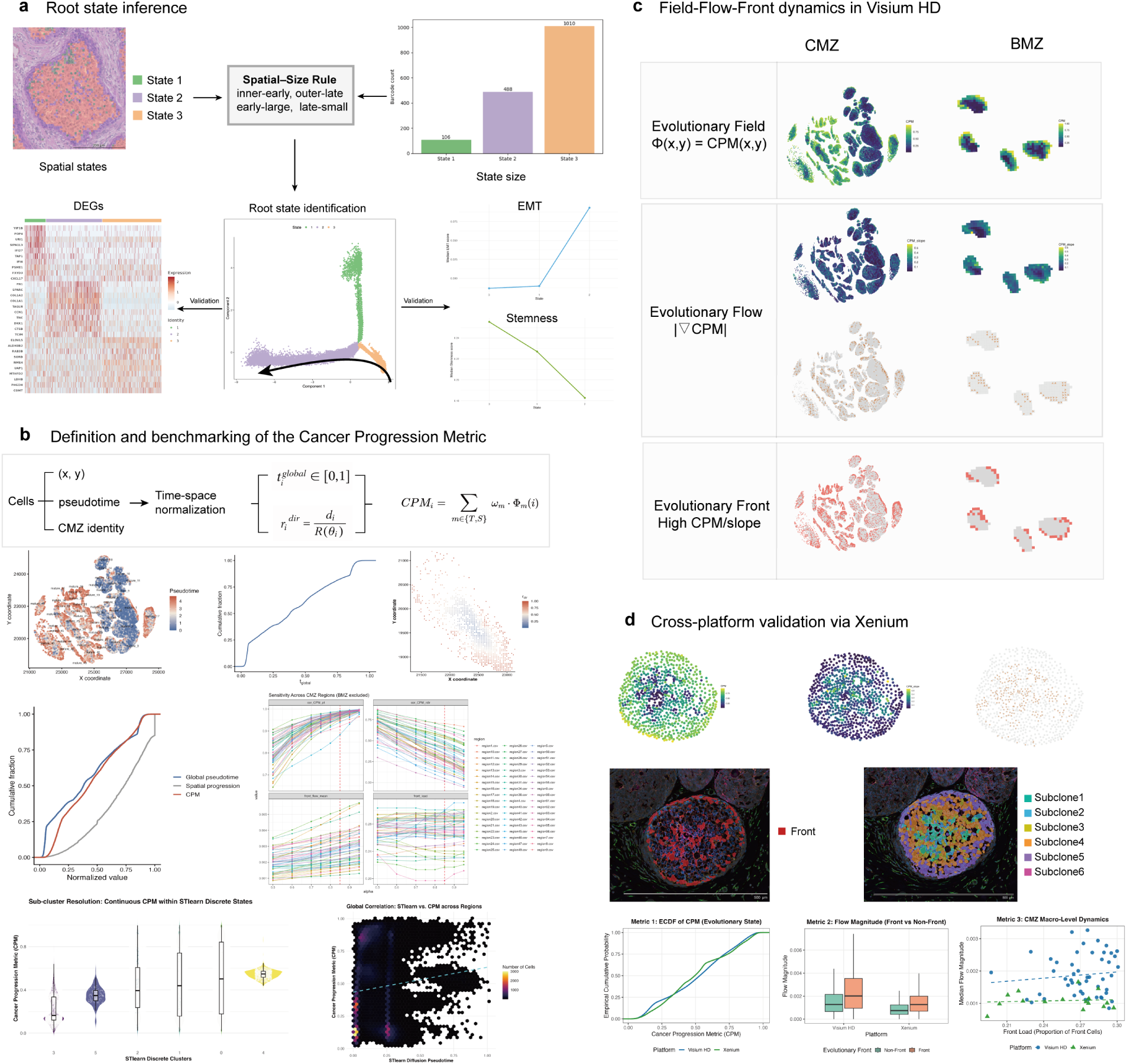
Genetic–spatial coupling derives the Field–Flow–Front model for resolving tumour evolutionary dynamics. **a,** Root state inference. Candidate root states were determined using a Spatial–Size Rule that integrates spatial localization (inner-early, outer-late) and state size (early-large, late-small). Root state identification was further validated by directional changes in EMT and stemness scores, together with differentially expressed genes (DEGs) across states. **b,** Cancer Progression Metric (CPM) integrates global transcriptional pseudotime (*t_i_*^global^) with local spatial progression (*r_i_*^dir^) within CMZs using a weighting parameter *α*. ECDF analysis indicates that CPM represents a unified evolutionary axis distinct from either component alone, and parameter sensitivity analysis (*α* = 0.5–0.95) demonstrates robustness of downstream metrics. Quantitative benchmarking reveals that CPM overcomes the terminal saturation of STlearn’s diffusion pseudotime and unmasks broad, continuous progression gradients within its discrete spatial clusters. **c,** The 3F model resolves the field, flow and front features of tumour evolution. Shown are the spatial maps of the evolutionary field (CPM), evolutionary flow (|∇CPM|) and evolutionary front (high CPM/high slope) in both CMZ and BMZ derived from the high-resolution Visium HD baseline data. **d,** Independent validation of 3F dynamics in single-cell resolution Xenium datasets. Shown is the independent replication of the 3F spatial maps in Xenium datasets. Cross-platform quantitative analyses demonstrate the robust generalizability of 3F dynamics, highlighting highly conserved empirical cumulative distributions of CPM, a substantial multi-fold elevation of flow magnitude in front cells across both modalities, and a consistent positive correlation between regional front load and median expansion momentum.

Because neither transcriptomic pseudotime nor physical coordinates alone captures continuous evolutionary history, we constructed CPM as a dual-clock metric grounded in the ecological principle of “space-for-time substitution”: normalized global transcriptional pseudotime serves as an internal clock of molecular progression, while a direction-normalized spatial distance acts as an external clock in which outward expansion encodes history. The two are integrated through a weighting hyperparameter α (Fig. 1c, Fig. 2b and Extended Data Fig. 3d). The macroscopic dynamic metrics underpinning all downstream analyses—front load and front flow magnitude—remained stable across α ∈ [0.5, 0.95] and preserved their inter-regional hierarchy, indicating that our conclusions are robust to rather than contingent on this weighting; we therefore fixed α = 0.85, at which CPM aligns closely with the transcriptional trajectory (Spearman *ρ* ≈ 0.97) while the spatial term contributes a targeted directional correction (Fig. 2b).

Empirical cumulative distribution analysis confirmed that CPM forms an independent evolutionary axis distinct from either component alone (Fig. 2b and Extended Data Fig. 3d). Benchmarked against the diffusion pseudotime (DPT) of the graph-based tool STlearn, CPM resolved broad, continuous progression gradients within individual STlearn clusters and retained discriminative resolution across regions where graph-based pseudotime saturated (Fig. 2b and Extended Data Fig. 3f), supporting CPM as a more continuous and spatially faithful coordinate of tumour evolution.

### Field–Flow–Front resolves tumour evolution as a continuous physical field rather than discrete clonal competition

While CPM defines a unified spatiotemporal metric embedded in tissue space at the single-cell level, a scalar metric alone cannot capture the higher-order spatial organization and dynamic patterns of tumour evolution. We therefore formalized the 3F model, projecting CPM into a spatial framework to decode the spatiotemporal organization of tumour evolution (Fig. 1d, Fig. 2c and Extended Data Fig. 4a). CPM is treated as a continuous scalar field over the tissue space, generating an Evolutionary Field Map (EFM) of evolutionary states within CMZs. Quantitative analysis revealed substantial inter-regional variation: the spatial proportion of the dominant evolutionary domain (Top-1) ranged from 0.21 to 0.44, with corresponding Shannon entropy values of 1.51 to 1.77, spanning a structural continuum from single-domain dominance to multi-domain coexistence.

From this field we derived the evolutionary flow, capturing the directionality and intensity of evolutionary progression. Flow strengths were markedly heterogeneous across regions: some CMZs exhibited concentrated, directionally coherent flows, whereas others showed dispersed or weakly directional expansion (Fig. 2c and Extended Data Fig. 4a), indicating that even cells occupying similar CPM intervals differ widely in the physical dynamics of their spatial expansion.

Finally, we defined the evolutionary front as regions of advanced CPM or steep local gradients and quantified their expansion capacity using the front load. Front load varied strikingly between regions—exceeding 30% of tumour cells in highly active regions while remaining much lower elsewhere (Fig. 2c and Extended Data Fig. 4a)—indicating that evolutionary activity is concentrated in specific physical niches rather than spatially uniform. Comparing core 3F metrics between BMZ and CMZ, CMZs exceeded BMZs on overall progression (CPM), local gradient (CPM slope), baseline expansion momentum (flow magnitude) and front-cell momentum (front flow magnitude) (all p < 0.0001; Extended Data Fig. 4a,b), establishing higher active expansion capacity at the tumour front.

Within this continuous framework, subclones emerge naturally as spatially contiguous regions occupying similar CPM isobands, allowing subclonal structure to be delineated without imposing rigid, tree-like phylogenies (Fig. 2e and Extended Data Fig. 4c). This continuous representation advances beyond discrete clonal sweeps, supporting tumour evolution as a continuous, population-level process of spatial and phenotypic adaptation rather than strict mutational competition, in line with models that concentrate selection at invasive fronts and tumour–stroma interfaces.^30–33^ Independent cross-platform validation in single-cell-resolution Xenium data closely reproduced the continuous trajectories and core micro-dynamics of the Visium HD baseline, confirming a multi-fold elevation of expansion momentum at front cells and establishing the 3F framework as a resolution-agnostic paradigm for decoding spatial tumour evolution (Fig. 2d and Extended Data Fig. 5a).

### Three-dimensional reconstruction confirms cancer microzones as sealed reactors compartmentalizing tumour evolutionary dynamics

Building on the cell-level subclonal reconstruction, we applied the 3F spatial clustering strategy across independent tumour samples, identifying 40 CMZ-level subclones with clearly defined boundaries. Macro-CPM correlated positively with front load—more advanced subclones contained a higher proportion of front cells (Fig. 2e)—suggesting that microenvironmental selection favours expansive, invasive phenotypes. Regardless of evolutionary stage, these subclones formed contiguous, boundary-defined patches rather than scattered cells (Fig. 2e and Extended Data Fig. 4c).

Assessing neighbouring CMZs by the evolutionary fitness index (EFI), higher-EFI subclones preferentially dominated adjacent CMZs and established sharper boundaries and competitive interfaces (Fig. 2e and Extended Data Fig. 3e), indicating localized clonal competition. CMZs thus function not merely as physical units but as discrete evolutionary compartments, with progression driven by spatially coherent, patch-like expansions rather than random single-cell diffusion.^34^

To test whether these two-dimensional patches have a three-dimensional structural basis, we reconstructed topological structures from serial sections. The CMZ-level subclones identified in 2D corresponded closely to anatomically independent units physically encapsulated by stroma in 3D, with high Z-axis continuity: most extended across more than seven consecutive sections to form spatially closed columnar or island-like volumes, ruling out artifactual inter-slice noise (Fig. 2e and Extended Data Fig. 4c).

Crucially, the 3D models directly visualize the isolation mechanisms between subclones of distinct evolutionary state: dominant high-Macro-CPM subclones remained segregated from neighbouring low-potential subclones by intact barriers of dense extracellular matrix (ECM), with no observable fusion or channel-like connections.^35^ CMZs are therefore not only structural building blocks of tumour growth but compartmentalized evolutionary reactors; by limiting clonal mixing, this physical isolation lets different CMZs evolve in parallel—much like independent petri dishes—generating macroscopic functional heterogeneity within the tumour.

### Spatiotemporal trajectories uncouple tumour progression from proliferation toward a low-dependency high-invasion front

The CPM axis resolves tumour transcriptional evolution into reproducible, stage-specific module switching rather than a homogeneous continuum. Stratifying CMZ-level populations into sequential subclones along the CPM axis (g1–g6), we found distinct transcriptional modules enriched within specific intervals, reflecting a phenotypic convergence in which cells decouple from environmental nutrient dependencies—tumour evolution propelled by discrete state transitions trading rapid proliferation for resilient survival and invasion. Across regions the core conserved modules were dominated by ECM and stromal remodelling, whereas immune-related modules emerged in a context-dependent, “inserted” manner; regional heterogeneity arose mainly from shifts in the timing and intensity of these modules rather than a redefinition of subclonal identity (Fig. 3a and Extended Data Fig. 5b).

**Fig. 3:**
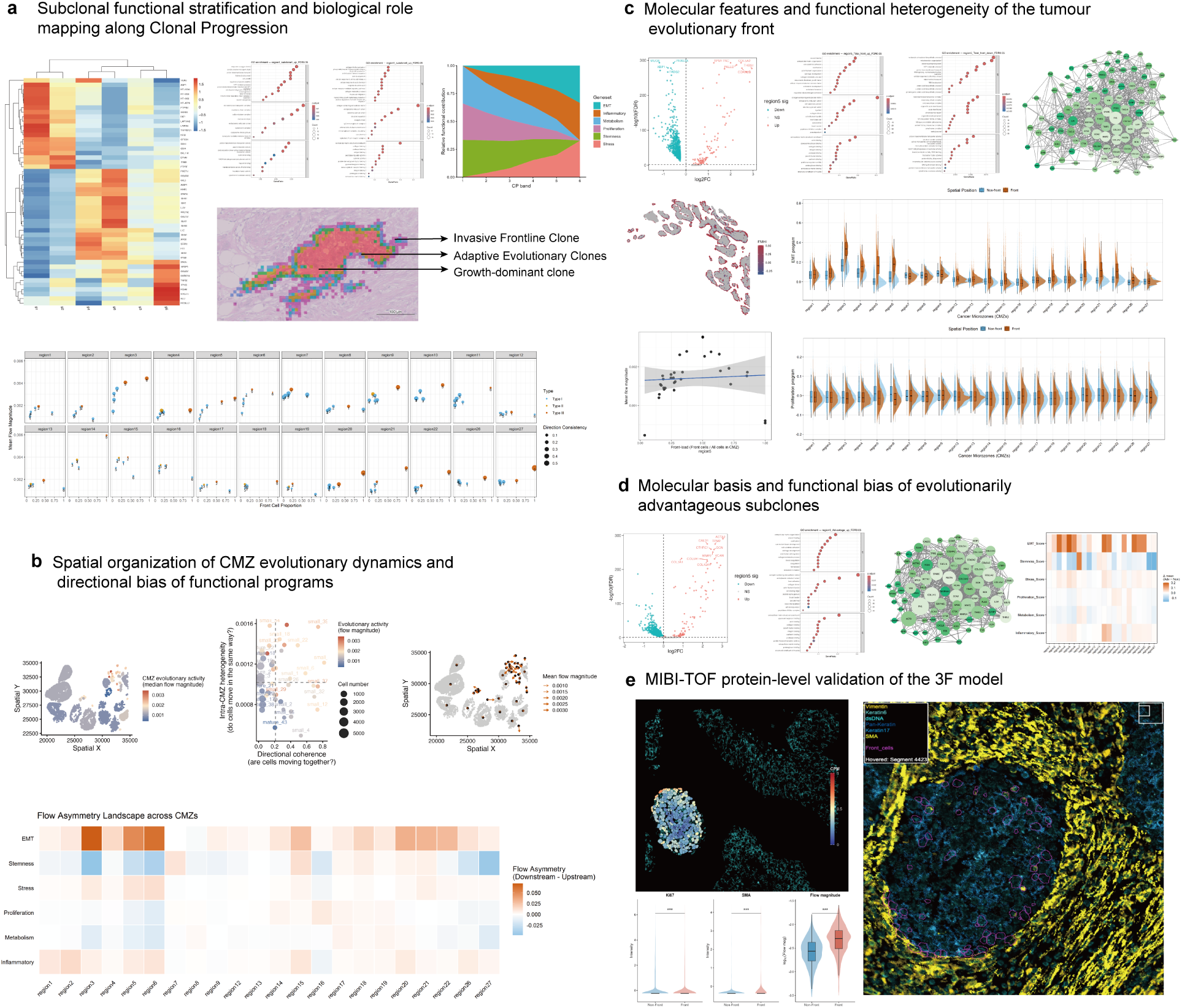
Core-to-front reprogramming decouples invasion from proliferation. **a,** Subclonal functional stratification along the CPM. Transcriptional trajectories resolve subclones (g1–g6) into distinct functional states, with heatmaps and gene ontology (GO) enrichment indicating stage-specific program transitions. Spatial mapping delineates growth-associated, adaptive and invasive frontline niches, and advanced CPM stages show increased flow magnitude and directional coherence. **b,** Spatial organization of evolutionary flow at the CMZ scale. Evolutionary activity (median flow magnitude) forms discrete spatial hotspots. Phase diagrams integrating directional coherence and intra-CMZ heterogeneity, together with mean flow vectors, define dominant expansion trajectories. Functional program distributions show upstream enrichment of stemness and downstream enrichment of EMT and stress programs along the evolutionary axis. **c,** Molecular characteristics of the invasive front. Differential expression analysis and protein–protein interaction (PPI) networks identify ECM-, adhesion- and migration-associated modules in front cells. Spatial regression links front load to flow magnitude, supporting the dynamic and adaptive properties of the evolutionary front. **d,** Molecular features of advantageous subclones. Subclones with high CPM values, steep gradients and strong flow exhibit consistent transcriptional advantages. Cross-regional comparisons and network analyses indicate that this advantage is associated with sustained extracellular matrix and stromal remodeling programs. **e,** Cross-platform validation of the 3F model on an independent MIBI-TOF triple-negative breast cancer cohort. A representative CMZ (Point 5) shows the “inner-low, outer-high” CPM-pp topology (left) and peripherally distributed front cells (magenta, right). Cohort-level plots show reduced Ki67, elevated SMA and ~2-fold higher flow magnitude in front versus non-front cells (Wilcoxon; \*\*\**p <* 0.001).

Trajectory and differential expression analyses resolved these subclones into three functional archetypes—growth-dominant, adaptive evolutionary, and invasive frontline—which mapped back onto histological space as a layered evolutionary architecture (Fig. 3a). Along the CPM axis, EMT and stress responses escalated while stemness, metabolism and proliferation were suppressed and inflammatory programs intensified with regional variability, delineating a transition from a “stemness/metabolism-dominant” state to a “stress/inflammation/EMT-dominant” state—a shift from localized proliferative expansion to a resource-conserving phenotype of adaptive survival, metabolic austerity and active migration (Fig. 3a). This stratification was tightly coupled to physical dynamics: late-CPM subclones were spatially enriched at the invasive front with amplified flow magnitudes and elevated directional coherence, and vector-field analysis assigned the terminal CPM stages (CPM5–6) to high-flow, directionally coherent (Type III) dynamics despite their diminished proliferative drive and low metabolic dependency (Fig. 3a).

At the CMZ level, evolutionary flow organized into discrete hotspots of high activity, and functional programs established a stable, spatially ordered hierarchy along the flow direction: stemness and metabolism upstream (low CPM), and EMT, stress and inflammatory programs downstream (high CPM) at the invasive front, with proliferation occupying an orthogonal, drift-like dimension (Fig. 3b and Extended Data Fig. 5b). Advancing clones thus shed their reliance on continuous nutrient influx and growth signalling, redirecting limited resources into a resilient, invasive, matrix-remodelling phenotype through a suppressed metabolic and proliferative baseline coupled with region-specific adaptive modules—pronounced mitochondrial and metabolic suppression accompanied by either extensive ECM remodelling or localized immune-interaction programs, consistent with metabolic reprogramming under microenvironmental stress^36^ and with tumour–immune shaping of progression.^37^ PPI analysis converged on a central ECM–adhesion–migration hub, anchoring the dynamic and functional identity of this environmentally resilient front (Fig. 3c and Extended Data Fig. 6a).

Synthesizing these insights, we defined advantage subclones as populations simultaneously exhibiting advanced CPM, steep local gradients and high flow magnitudes—the late-stage, persistently advancing units driving progression. Orthogonal differential expression, GO enrichment and PPI analyses identified ECM and stromal remodelling as their indispensable backbone, and cross-regional comparison showed this evolutionary advantage to be consistently conserved across single-gene, pathway and higher-order program levels, defining a highly invasive, structurally remodelling, environmentally decoupled consensus state (Fig. 3d and Extended Data Fig. 6b).

To validate these spatiotemporal architectural features at the protein level on an orthogonal platform, we projected the 3F framework onto an independent MIBI-TOF imaging cohort of triple-negative breast cancer, in which a protein-based CPM-pp was constructed using Vimentin and Beta-catenin as proxies for the EMT–Wnt invasion axis. Across CMZs, CPM-pp consistently recapitulated the “inner-low, outer-high” spatial topology observed in Visium HD data, with MIBI-defined front cells distributing along the CMZ periphery and stromal interface rather than within the proliferative core (Figure 3E and S6C). Independent of the markers used in CPM-pp construction, front cells exhibited reduced Ki67 ( *ρ* = −0.124, p < 0.001), elevated SMA ( *ρ* = +0.218, p < 0.001), and an approximately two-fold increase in evolutionary flow magnitude (1.87×, p < 0.001)(Figure 3E). These cross-platform observations confirm at the protein level that advancing tumour fronts simultaneously suppress proliferation, acquire mesenchymal–matrix-remodelling features, and accelerate spatial expansion momentum, providing orthogonal validation of the low-dependency, high-invasion state predicted by the 3F model.

### A non-cell-autonomous ECM programme drives the evolutionary front beyond clonal selection

To biologically validate the 3F model at the mechanistic level, we sought the core molecular programmes sustaining evolutionary front formation, integrating CPM-resolved transcriptional switching, network-based driver inference, genetic dependency profiling, spatial interaction analysis, protein-level imaging and therapeutic validation. Across these complementary analyses, the results consistently converged on a conserved front-associated programme as the principal engine of tumour evolutionary progression.

To map the transcriptional events propelling the front, we performed CPM-GeneSwitch analysis across regions, annotating genes progressively deactivated (ON→OFF) or activated (OFF→ON) along the CPM axis, which showed profound functional convergence. Genes silenced with advancing CPM corresponded to intrinsic homeostasis and metabolic–secretory programs (mitochondrial respiration, lipid metabolism, ER/secretory stress, cell cycle, basal epithelial markers), declining in a layered pattern in which epithelial identity fell early while metabolic and mitochondrial pathways persisted to higher CPM. Conversely, genes activated along the axis were enriched in extracellular matrix (ECM) remodelling with amplified inflammatory and stress responses, following a temporally choreographed cascade from *COL1A1* at early stages (switch cpm ≈ 0.05–0.12), through *SPARC* -related modules at intermediate-to-high CPM (≈ 0.4–0.8), to *MMP2* peaking at the extreme front (≈ 0.85–0.95) (Fig. 4a and Extended Data Fig. 7a–b).

**Fig. 4:**
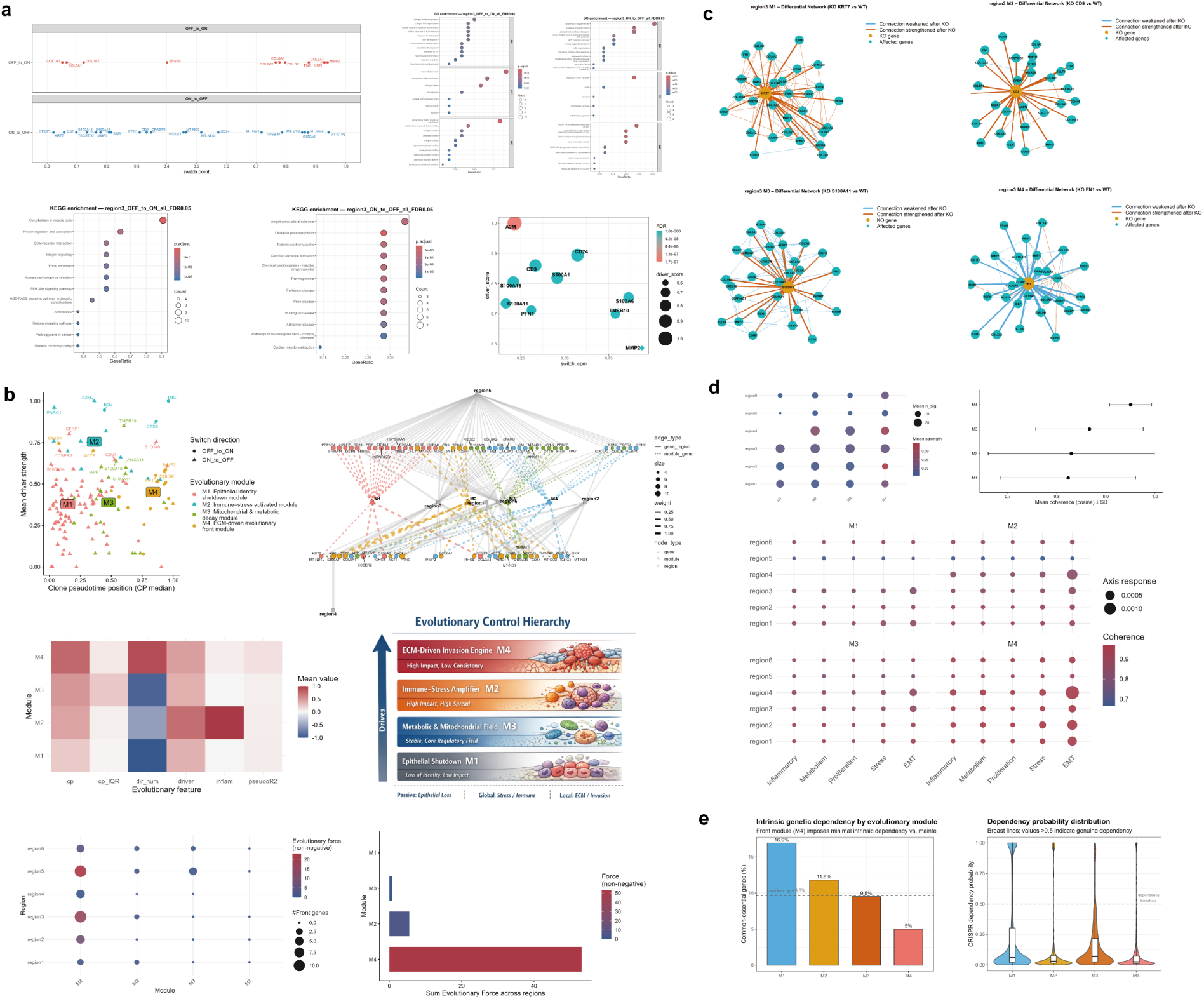
An ECM-driven front module (M4) drives progression independently of cell-autonomous gene essentiality. **a,** Temporal dynamics of evolutionary driver genes along the CPM. CPM-GeneSwitch analysis identifies activation (OFF→ON) and repression (ON→OFF) of transcriptional programs across CPM (region3 shown as a representative example). Gene ontology (GO) and KEGG enrichment analyses indicate stage-specific functional transitions. Integration of protein–protein interaction (PPI) networks with a graph attention network (GAT) identifies candidate driver genes based on switch position and driver score. **b,** Identification and organization of driver modules. Cross-regional integration groups driver genes into four modules (M1–M4) distributed along the CPM axis. Bipartite networks and feature heatmaps indicate cross-regional conservation, and module positioning reflects progressive transitions from early epithelial programs to late extracellular matrix (ECM) and stress-associated programs. **c,** Perturbation analysis of driver genes using in silico knockout. scTenifoldNet-based analysis evaluates regulatory network changes following removal of candidate drivers. Network representations show perturbation of downstream targets and alterations in network connectivity. **d,** Quantification of perturbation effects and cross-regional consistency. Bubble plots summarize the extent and magnitude of network perturbations, and cosine similarity analysis indicates consistent effects across regions. Module-specific responses are observed across multiple functional programs, including inflammation, metabolism, proliferation, stress and epithelial–mesenchymal transition (EMT). **e,** Intrinsic genetic dependency of evolutionary modules in DepMap. Across breast cancer cell lines, the proportion of common-essential genes declines from maintenance modules M1/M3 to the M4 ECM/front module (5.0%, ∼half of a size-matched random background of ≈9.6%; left), and CRISPR dependency probabilities for M4 fall almost entirely below the 0.5 dependency threshold (right), indicating that M4 genes are not required for tumour-cell survival in isolation.

To distinguish drivers from passengers, we developed EvoDrGK (Evolutionary Driver Gene identification based on GeneSwitch kinetics), inferring network-aware driver scores from STRING PPI networks with a graph attention network (Fig. 1f). High-scoring drivers overwhelmingly converged on ECM structural components and inflammation–stress coupling nodes, with ECM genes establishing the physical scaffold at low-to-intermediate CPM and inflammation/stress regulators enriched subsequently, coupling matrix remodelling to inflammatory amplification^38^ (Fig. 4a and Extended Data Fig. 7c). Clustering drivers across regions yielded four evolutionary modules (M1–M4) (Fig. 4c and Supplementary Table 2), whose front-directed evolutionary force was quantified within the 3F framework. M4 (the ECM-driven front module) exerted by far the strongest positive force (total force = 53.23 versus 5.56 for the M2 immune–stress module), peaking in highly active regions, whereas M3 (mitochondrial/metabolic decay) and M1 (epithelial-identity shutdown) contributed minimally (Fig. 4b). In silico knockout with scTenifoldNet established a causal hierarchy: targeting M4 drivers (*COL1A1*, *FN1*, *MMP2*) collapsed ECM, cytoskeletal and migration networks, whereas disrupting M2 (*B2M*, *PNRC1*, *A2M*), M3 (*MT-ND1*, *TMSB10*, *LMNA*) or M1 (*EIF4EBP2*, *UBA52*, *TRIB3*) perturbed inflammation/stress, mitochondrial/nuclear stability, or transcriptional–metabolic regulation respectively without dismantling the ECM architecture; M4 showed the most profound and coherent systemic effects across EMT, stress and inflammatory programs (Fig. 4c,d and Extended Data Fig. 7d).

To elucidate the spatial inputs sustaining these front states, we applied a distance-stratified, sender-centred NicheNet analysis. The cancer-associated fibroblast (CAF)→Tumour axis was the most conserved interaction across regions (ligand–receptor strength 0.12–0.38, peaking at 0.35–0.38 in region3, region5 and region6), with tumour-proximal CAFs inducing ECM and invasive responses through ligands enriched in ECM, stromal and growth-factor families (collagen ligands, TGF-β/FGF/IGF), whereas Immune→Tumour and Tumour→stroma interactions were weaker and context-dependent (Fig. 5a and Extended Data Fig. 8). Mapped onto the evolutionary modules, CAF→Tumour interactions were overwhelmingly dominated by M4 in interaction strength (mean ∼0.25 versus <0.02 for other modules), indicating that conserved stromal inputs preferentially reinforce the tumour-intrinsic M4 ECM/front programme and stabilize the invasive front (Fig. 5a). To corroborate this at the protein level, we performed CODEX proteomic imaging across manually delineated CMZs (23 ROIs from 10 patients), scoring an invasiveness/ECM programme (VIM, COL4A2, PDPN, inverted KRT19) and an inflammatory-stress programme (HIF1A, HLA-DRA) along the core-to-periphery axis. The invasiveness/ECM programme increased markedly toward the periphery while the inflammatory-stress programme showed only a weak gradient (Fig. 5b and Extended Data Fig. 8b), confirming that front enrichment is predominantly M4- rather than M2-driven. Projected onto genome-wide CRISPR dependency data (DepMap), the proportion of common-essential genes declined along the evolutionary axis, from 16.9% in M1 to 9.5% in M3 and only 5.0% in M4, roughly half of a size-matched random background (≈9.6%); M1 and M3 retained high-dependency tails toward 1.0 whereas M4 collapsed almost entirely below the 0.5 threshold (Fig. 4e). Thus, although M4 is the top causal driver of front formation, its genes are not required for tumour-cell survival in isolation, mirroring the division of labour seen by knockout and implying that the M4 front advantage is coupled to the surrounding tissue ecology rather than encoded by cell-autonomous survival genes.

**Fig. 5:**
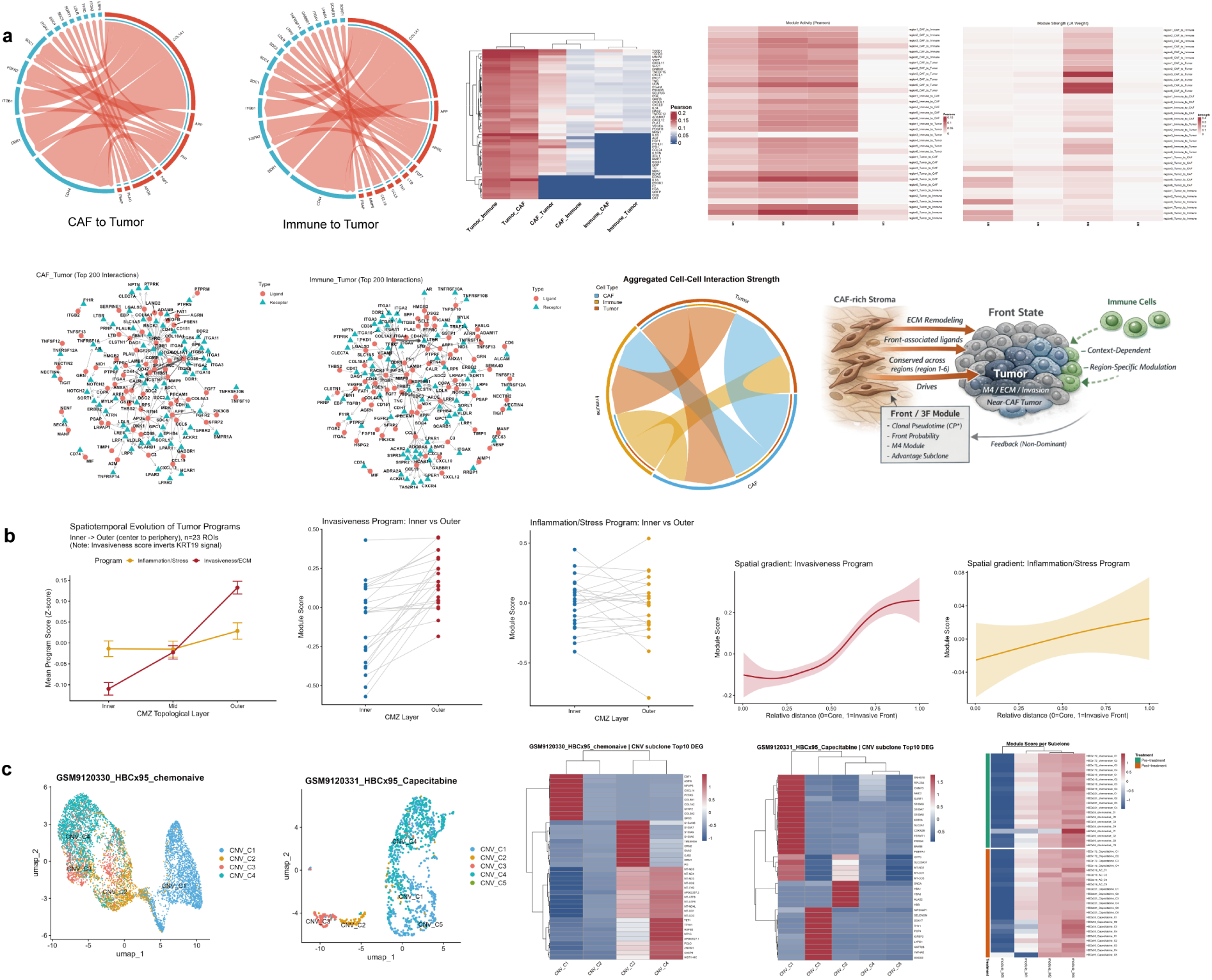
CAF-mediated interactions and protein-level validation establish the M4-driven evolutionary front and its therapeutic resilience. **a,** Spatial cell–cell communication at the evolutionary front. Distance-stratified interaction networks show dominant signalling from cancer-associated fibroblasts (CAFs) to tumour cells. Ligand activity analysis and interaction strength heatmaps indicate enrichment of collagen–integrin and growth factor signalling associated with the M4 (ECM/front) module across regions. Immune-to-tumour interactions are comparatively weaker and vary across spatial contexts. A schematic summarizes the hierarchical organization of microenvironmental interactions linking CAF activity to ECM remodeling and front-associated states. **b,** CODEX protein-level validation across CMZs: along the core-to-periphery (Inner→Outer) axis, the invasiveness/ECM score (*VIM*, *COL4A2*, *PDPN*, inverted *KRT19*) rises markedly toward the invasive front, whereas the inflammatory-stress score (*HIF1A*, *HLA-DRA*) shows only a weak gradient, confirming at the protein level that front enrichment is predominantly M4- rather than M2-driven. Error bars, s.e.m.; shaded bands, 95% CI. **c,** Evolutionary dynamics under therapeutic selection. Single-cell profiling of paired pre- and post-chemotherapy PDX models (e.g., HBCx95, chemonaive versus capecitabine-treated) shows transcriptional remodeling of copy-number-defined subclones. Subclone tracking and differential expression analyses indicate persistence of ECM-associated transcriptional programs in post-treatment subclones across samples.

Finally, to test robustness under therapeutic selection, we analysed paired pre- and post-chemotherapy PDX samples from five TNBC patients (10 samples), scoring signed module activity (M1–M4) at the copy-number (CNV) subclone level (Supplementary Table 3). M4 remained the sole consistently positive module across surviving subclones while M1–M3 were generally suppressed, indicating that residual tumour expansion continues to rely on ECM-driven front dynamics.^39^ Rather than uniformly scaling M4, therapy redistributed which CNV subclones carried the M4 programme while rewiring associated stress-response networks, through two reproducible “rescue” trajectories:^1^ a Triage–Heterogeneity pattern, in which pre-existing M4-carrying subclones shifted or converged alongside compensatory stress and proliferation responses, and a Plasticity/Front-Rewiring pattern, in which new M4 carriers emerged after treatment through transcriptional reprogramming of pre-existing subclones, frequently coupled with M2 (Fig. 5c and Extended Data Fig. 9). These models establish M4 as a transferable, cross-sample engine of tumour evolution whose clonal carriers are reshuffled under treatment.

### Evolutionary Advantage Load translates local spatial dynamics into cross-scale clinical outcome prediction

To extend the 3F model from tissue-level dynamics to clinical outcome and test its cross-scale validity, we analysed two independent breast cancer cohorts, TCGA-BRCA^40^ and I-SPY1 (GSE32603),^41^ which differ in population origin and platform but are comparable in age and molecular-subtype composition (Extended Data Fig. 10b). Because I-SPY1 is better suited to recurrence and TCGA to overall survival (OS), we adopted an endpoint-matched design, evaluating recurrence-free survival (RFS) and OS in their respective optimal contexts.

From the spatial transcriptomics data we constructed a weakly supervised Top80 advantage signature (Fig. 6a, Extended Data Fig. 10a,c and Supplementary Table 2). Projected onto scRNA-seq data (GSE161529), it identified advantage-like states within tumour epithelial cells, from which we defined the sample-level evolutionary advantage load (E-load) as the proportion of advantage-like cells (Fig. 6b and Supplementary Table 4). To scale E-load to bulk cohorts, we trained a model on routine clinical markers (ER, PR, HER2, Ki67), selecting a quasibinomial model to predict E-load in bulk RNA-seq samples (Supplementary Table 5); unsupervised K-means (k = 3) stratified samples into Low, Mid and High E-load groups (Fig. 6c and Extended Data Fig. 10a).

**Fig. 6:**
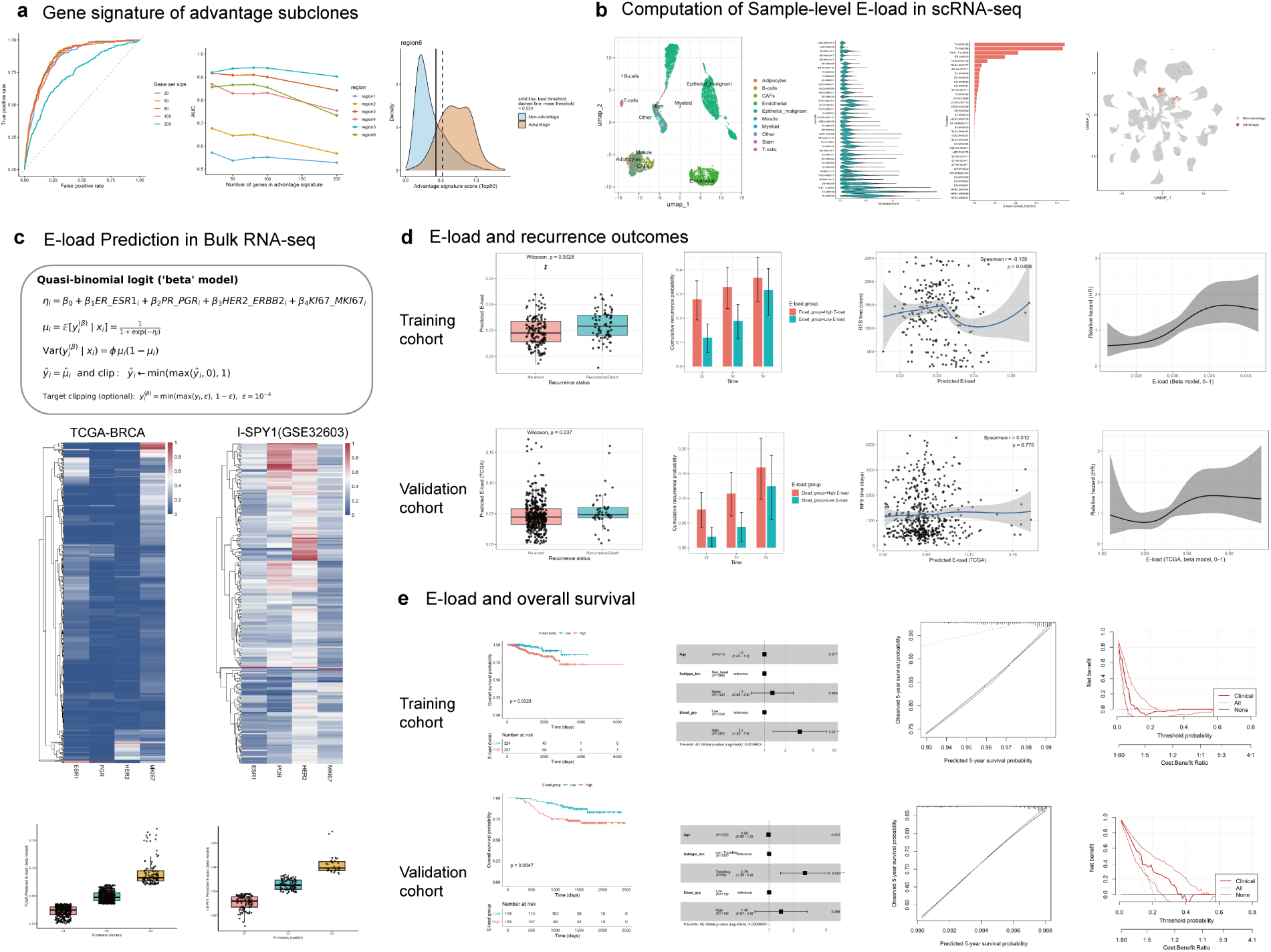
Macroscopic Evolutionary Advantage Load enables cross-scale prediction of recurrence and survival. **a,** Derivation of the advantage signature associated with advantage subclones, with a unified threshold (0.524) applied across spatial regions. **b,** Single-cell quantification of E-load. The spatial signature identifies advantage-like malignant states in scRNA-seq data, and E-load is defined as the sample-level fraction of these cells; E-load showed a right-skewed distribution across 38 samples (range 0–0.438; median 0.002; IQR 0–0.022). **c,** Cross-scale projection to bulk cohorts. E-load is estimated in bulk transcriptomic datasets from routine clinical markers (ER, PR, HER2 and Ki67), stratifying patients into three risk groups. **d,** Association between E-load and recurrence risk. Across I-SPY1 and TCGA cohorts, higher baseline E-load is associated with increased recurrence probability (I-SPY1: *r* = −0.13, *p* = 0.046; 2-year 27.8% vs 11.9% and 3-year 33.0% vs 18.9% for high vs low E-load), with a continuous E-load–hazard relationship. **e,** Prognostic performance for overall survival. Elevated E-load is associated with adverse overall survival (TCGA-BRCA: log-rank *p* = 0.0023; adjusted HR = 3.21) and improves risk prediction when added to clinical covariates (C-index 0.685 → 0.717; 3-year OS AUC 0.675 → 0.712), with consistent performance on external validation in I-SPY1 (log-rank *p* = 0.0047; *p* = 0.04 for added value).

We then evaluated the prognostic value of E-load. In I-SPY1, E-load was negatively correlated with RFS (r = −0.13, p = 0.046), with recurrence risk rising monotonically; high-E-load patients showed substantially higher recurrence (2-year 27.8% vs 11.9%; 3-year 33.0% vs 18.9%) (Fig. 6d and Extended Data Fig. 10e). A congruent trend held in TCGA, where restricted cubic spline Cox models confirmed an increasing hazard ratio with higher E-load despite fewer recurrence events. For OS in TCGA-BRCA, high E-load was significantly associated with worse survival (log-rank p = 0.0023; adjusted HR = 3.21), and incorporating E-load improved prognostic performance over clinical covariates alone, raising the C-index from 0.685 to 0.717 and 3-year OS AUC from 0.675 to 0.712, with well-calibrated 5-year survival predictions and no evidence of overfitting (Fig. 6e and Extended Data Fig. 10e). External validation in I-SPY1 corroborated these findings: despite smaller size and treatment heterogeneity, high E-load remained associated with worse survival (log-rank p = 0.0047), and likelihood-ratio testing confirmed significant additional explanatory value beyond standard clinical metrics (p = 0.04) (Fig. 6e and Extended Data Fig. 10e).

Collectively, the prognostic detriment of E-load was reproducible across heterogeneous populations and divergent platforms, affirming both its cross-scale transferability and the clinical validity of the 3F model. The cumulative macroscopic burden of these advantage-like states thus constitutes a quantifiable evolutionary dimension linking tissue-level front dynamics to patient outcome.

## Discussion

Conventional models equate malignant advance with proliferative capacity, treating fast-cycling clones as the fittest units of selection.^30, 31^ Our spatiotemporal trajectories overturn this at the invasive front: along the CPM axis, proliferation is orthogonal to evolutionary progression, and the most advanced front states are the least proliferative and least metabolically dependent yet show the highest directional expansion momentum. Tumour evolution thus resolves not into a homogeneous continuum but into reproducible, stage-specific module switching that converges—across regions and at the protein level—onto a single low-dependency, high-invasion state defined by ECM remodelling and motility rather than growth. Fitness at the front is better understood as sustained spatial expansion under resource austerity than as rapid division.

What drives an advance uncoupled from the cells’ own survival machinery? Conventional frameworks assume the genes conferring evolutionary advantage are those required for survival; our data suggest otherwise. The ECM programme (M4) we identify as the principal causal driver of the front is, by CRISPR dependency profiling, among the least cell-essential of all modules, indicating that its selective advantage is encoded not in tumour-cell-autonomous survival genes but in ecological coupling to the stroma. Extending models that concentrate selection at invasive fronts and tumour–stroma interfaces,^32–34^ we resolve the front into a stromal-reinforced ECM programme that, critically, is preserved under therapy while its clonal carriers are reshuffled—arguing that the operative unit of front evolution is a transferable programme rather than any single clone, a program-centric view that in this respect takes precedence over the clone-centric paradigm.

These conclusions rest on reconstructing evolution within native tissue. The CMZ—a stroma-encapsulated region whose cells approximate a clonal expansion from a common ancestor—grounds the ecological space-for-time substitution and helps overcome the clonal illusion that follows when spatial context is discarded.^42, 43^ On this unit the CPM becomes interpretable, anchoring a predominantly transcriptional clock with a direction-normalized spatial prior (r*^dir^*); macroscopic dynamics remain stable across a wide range of α, indicating robustness to rather than dependence on this weighting. The design exploits the complementary error structures of its two coordinates—dense, deterministic spatial geometry versus sparse, directionally unstable pseudotime under Visium HD asymmetry—stabilizing molecular ordering while preventing spatial signals from over-smoothing genetic heterogeneity.^25, 26^ Together these let the 3F model render Field, Flow and Front as a self-consistent physical map that corresponds directly to tissue histology.^33, 44^

Finally, the framework yields a clinically transferable measure: the E-load captures the cumulative burden of this conserved front-driving process and projects local evolutionary advantage to the patient level, linking mesoscopic dynamics to outcome across platforms.^45, 46^ E-load thus substantiates, rather than stands apart from, the cross-scale interpretability of the 3F model, supporting a shift from static molecular classification towards evolution-informed prognosis and intervention.^34, 47, 48^ Overall, we establish a tissue-embedded cartographic paradigm in which clonal behaviour is read through spatial position, structural context, and progression state—a generalizable basis for understanding, and ultimately intercepting, how solid tumours advance.

Our study has several limitations. First, temporal ordering is inferred computationally from spatial and transcriptional states rather than observed longitudinally, and requires prospective validation. Second, current spatial platforms are limited in field of view and coverage, constraining recovery of lesion-wide trajectories. Third, the stromal mechanisms implicated in sustaining the front—particularly CAF-reinforced ECM dynamics—remain computationally inferred and await direct functional perturbation. Fourth, the therapeutic and prognostic analyses, though orthogonal, are limited in cohort size, especially the paired chemotherapy PDX models and the cohorts underlying the E-load associations, constraining subclone-level power and E-load precision. Resolving these will require whole-slide, multi-region longitudinal profiling with lineage tracing, in situ perturbation of the M4 programme, and validation in larger, uniformly treated cohorts.

## Methods

### Experimental methods

#### Specimens and sample processing

Five formalin-fixed paraffin-embedded (FFPE) tumour samples from patients with triple-negative breast cancer (TNBC) were included in this study (Supplementary Table 1). All samples were obtained from Sichuan Cancer Hospital with written informed consent from all patients, and the study was approved by the institutional ethics committee (approval number: SCCHEC-02-2023-049). Eligible patients underwent surgical resection between January and December 2023, with histopathological confirmation of TNBC. TNBC status was defined according to immunohistochemical criteria, including oestrogen receptor (ER) and progesterone receptor (PR) expression < 1% and absence of HER2 overexpression. All samples were independently reviewed and confirmed by two pathologists. For inclusion, samples were required to have preserved tissue architecture and tumour cell content greater than 60%. Patients with a history of other malignancies or who had received neoadjuvant therapy prior to surgery, and samples exhibiting severe degradation or extensive necrosis, were excluded.

### Serial sectioning

Serial consecutive FFPE sections were generated for each sample. Sections were prepared using a rotary microtome (Leica RM2016, Leica Microsystems) at a thickness of 5 µm with consistent orientation and sectioning conditions. Sections were sequentially numbered and preserved in order. Adjacent sections were collected at uniform intervals to maintain spatial correspondence across tissue layers. These serial sections were used for histological comparison, spatial registration and downstream three-dimensional reconstruction analyses.

### Tissue fixation, embedding and H&E staining

Breast cancer tissue samples were fixed in 10% neutral buffered formalin, followed by graded dehydration and paraffin infiltration using an automated tissue processor (Leica ASP300S, Leica Microsystems), and subsequently embedded in paraffin. Paraffin-embedded tissues were sectioned into consecutive 5 µm-thick slices using a rotary microtome (Leica RM2235, Leica Microsystems) and mounted onto charged glass slides to enhance tissue adhesion. Paraffin sections were deparaffinized in xylene and rehydrated through a graded ethanol series (100%, 95%, 85%, 75%) to distilled water. Hematoxylin and eosin (H&E) staining was then performed at room temperature. Sections were stained with hematoxylin solution (Sigma-Aldrich), rinsed under running water, differentiated in 1% acid alcohol, and subsequently blued in a weak alkaline solution. Sections were then counterstained with eosin (Eosin Y, Sigma-Aldrich) to visualize cytoplasmic and stromal structures. After staining, sections were dehydrated through graded ethanol, cleared in xylene, and mounted using neutral balsam (Neutral Balsam, Solarbio). All sections were processed under identical conditions.

### Whole-slide scanning and histological imaging

H&E-stained and coverslipped sections were digitized using whole-slide imaging systems, including the SQS-600P (SQray Technology, Shenzhen, China) and the Panoramic 250 digital slide scanner (3DHISTECH, Hungary). Slides were first scanned at 20× magnification to generate overview images covering the entire tissue section, followed by imaging at 40× resolution for representative regions. The acquired images were stored as whole-slide images (WSIs) in proprietary multi-resolution pyramid formats, including .sdpc and .mrxs. Digital slides were visualized and managed using CaseViewer (3DHISTECH) and ImageViewer (SQray), which were used for histological inspection, pathological feature annotation and assessment of spatial correspondence across serial sections. Selected WSIs were converted into standard TIFF image format without loss of spatial resolution to enable downstream computational analysis and three-dimensional reconstruction.

### RNA quality control for Visium HD

To ensure the robustness and reliability of Visium HD FFPE spatial transcriptomics experiments, RNA quality was assessed for all breast cancer FFPE samples prior to library preparation. RNA was extracted using the RecoverAll™ Total Nucleic Acid Isolation Kit for FFPE (Thermo Fisher Scientific, AM1975).^49^ RNA purity, concentration and fragment distribution were evaluated using a NanoDrop spectrophotometer (Thermo Fisher Scientific)^50^ and an Agilent Bioanalyzer (Agilent Technologies).^51, 52^ Because RNA derived from FFPE samples is often fragmented, the percentage of RNA fragments longer than 200 nucleotides (DV200) was used as the primary quality control metric. Additional criteria included RNA concentration and absorbance ratios (A260/280 and A260/230). Sample quality thresholds were defined according to the manufacturer’s recommendations for Visium HD and CytAssist FFPE spatial transcriptomics workflows (10x Genomics), including: (1) DV200 > 30%; (2) A260/280 ratio between 1.8 and 2.2; and (3) sufficient RNA input for library construction. Bioanalyzer electropherograms were used to assess RNA fragment size distribution and degradation profiles. Only samples meeting all quality criteria were retained for downstream analyses.

### FFPE sectioning and Visium HD slide preparation

Breast cancer FFPE tissues containing target tumour regions were serially sectioned at a thickness of 5 µm and mounted onto glass slides (Sigma-Aldrich, P0425). Sections were air-dried at room temperature. Tissue sections were placed within the Capture Area of the Visium HD slide to ensure sufficient coverage of the target regions for spatial transcriptomic signal capture.

### Deparaffinization and decrosslinking

Deparaffinization, rehydration, hematoxylin and eosin (H&E) staining, and decrosslinking of FFPE sections were performed according to the 10x Genomics FFPE sample preparation guide (CG000684).^53^ Following staining, brightfield whole-slide imaging was performed to obtain high-resolution histological images. These images were used for tissue region identification, histopathological annotation and spatial registration in downstream spatial transcriptomic analyses.

### Probe hybridization and ligation

Tissue sections were processed using 10x Genomics Visium HD slides (6.5 mm, PN-2000970) and Visium HD Spatial Gene Expression Reagent Kits (human whole transcriptome, 6.5 mm, PN-1000675). Probe hybridization was performed at 50^◦^C overnight (16–24 h) to allow binding of gene-specific probes to target RNA. Following hybridization, post-hybridization washes were conducted at 50^◦^C, after which probe ligation was performed to generate amplifiable ligation products.

### CytAssist transfer and spatial barcode capture

After probe ligation, tissue sections were aligned with the capture slide using the Visium CytAssist system. Through enzymatic reactions, ligation products were released from the tissue and transferred onto the spatially barcoded oligonucleotide array on the capture slide. The Visium HD capture area consists of densely arranged ∼2 µm barcoded squares, enabling near single-cell resolution spatial transcriptomic profiling. Each barcoded square contains a unique molecular identifier (UMI) and a spatial barcode, allowing precise assignment of transcript spatial origin.

### Visium HD library construction and high-throughput sequencing

Captured spatially barcoded ligation products were extended, eluted and pre-amplified for library construction. Libraries were generated by sample index PCR and purified using SPRIselect beads for fragment selection. Library quality control, including fragment size distribution and concentration, was performed prior to sequencing. Sequencing was performed on an Illumina platform using paired-end sequencing (PE150 mode), where Read 1 captures spatial barcode and UMI information and Read 2 sequences probe-derived transcript sequences. Sequencing depth followed 10x Genomics recommendations, with approximately 275 million read pairs per Capture Area for FFPE samples to ensure sufficient coverage and data quality. All experimental procedures were conducted according to the Visium HD Spatial Gene Expression User Guide (CG000685).^54^

### Analytical methods

#### Sequencing data processing and spatial gene quantification

Raw sequencing data were generated in FASTQ format and processed using the 10x Genomics Space Ranger pipeline (v3.0.1), including image alignment, tissue detection, read alignment and gene expression quantification. Brightfield histological images were first aligned to the capture array to identify tissue-covered regions. Sequencing reads were then aligned to the reference genome (human: GRCh38), and spatial assignment was performed based on spatial barcodes. In Visium HD data, each 2 × 2 µm barcoded square represents the minimal spatial unit for gene expression quantification. To improve signal robustness, neighbouring squares were aggregated to generate spatial bins at 8 × 8 µm and 16 × 16 µm resolutions. During downstream data analysis, to balance signal stability and spatial precision, the original spatial signals were primarily integrated to an effective analytical resolution of 8 µm. For each resolution (2 µm, 8 µm and 16 µm), quality metrics including total reads per bin, number of detected genes and unique molecular identifier (UMI) counts were computed to assess sequencing depth and data quality (Supplementary Table 1).

### Quality control and preprocessing of gene expression matrices

Gene expression matrices generated by Space Ranger were further processed using Seurat (v5.1.0).^55^ Spatial bins at 8 × 8 µm resolution were selected for downstream analyses. Low-quality bins were filtered based on gene counts and UMI counts to remove potential background or low-quality signals. Data normalization was performed using the NormalizeData function in Seurat with default log-normalization, correcting for differences in sequencing depth across bins.^56^

### Dimensionality reduction and unsupervised clustering

Highly variable genes (HVGs) were identified using the FindVariableFeatures function in Seurat, and the top 3,000 HVGs were selected for downstream analysis. Principal component analysis (PCA) was performed based on the expression matrix of HVGs for dimensionality reduction, and the top principal components were retained. A shared nearest neighbour (SNN) graph was constructed in PCA space, followed by unsupervised clustering to identify spatial regions with similar transcriptional profiles. Uniform Manifold Approximation and Projection (UMAP) was applied for nonlinear dimensionality reduction and visualization in two-dimensional space.

### Spatial cell type annotation using robust cell type decomposition (RCTD)

To infer the cellular composition within each spatial unit, we used the breast cancer single-cell RNA sequencing dataset (GSE176078) published by Wu et al. in Nature Genetics (2021)^57^ as the reference dataset to perform cell type annotation on Visium HD spatial transcriptomics data. Spatial cell type deconvolution was carried out using the spacexr (version 2.2.0) package. This method leverages cell type-specific transcriptional signatures derived from single-cell RNA-seq data to robustly decompose mixed signals in spatial transcriptomics data, while correcting for systematic differences between sequencing platforms. In the implementation, the create.RCTD function was used with default parameters, with additional constraints that each cell type must contain at least one single cell and each spatial bin must contain at least 11 unique molecular identifiers (UMIs). Subsequently, in the run.RCTD function, the mode parameter was set to “doublet” to infer the major cell type composition within each spatial bin.^58^

### Construction of cancer microzones

To visualize the spatial cell-type distribution predicted by RCTD and to provide a basis for subsequent microregion-level analyses, we used Loupe Browser (v8.0.0, 10x Genomics) to visualize the spatial transcriptomics data. Based on tumour-related cell-type annotations, Cancer.Basal.SC, Cancer.Cycling, Cancer.Her2.SC, Cancer.LumA.SC, and Cancer.LumB.SC were selected as the primary markers of tumour cells and used to define candidate tumour regions, thereby avoiding contamination from immune or stromal cells.

On this basis, integrating spatial mapping information from serial sections and histopathological features, cancer microzones (CMZs) were systematically defined and delineated. The criteria for CMZ identification were as follows: (1) clear tumour epithelial identity, confirmed by RCTD mapping; (2) relative local independence, containing at least two spatially adjacent tumour cells while remaining spatially separable from surrounding tumour-cell populations; (3) distinct stromal encapsulation; and (4) well-defined histological boundaries. To ensure the biological relevance of the constructed CMZs, regions with high tumour-cell density were preferentially annotated, and multiple rounds of manual inspection and consistency verification were performed to avoid omission of key regions or inclusion of non-tumour cells.

The same identification criteria were applied across platforms. For single-cell-resolution Xenium data, CMZs were delineated directly from single-cell tumour annotations together with histopathological priors, following the same criteria used for Visium HD. For the protein-based MIBI-TOF and CODEX platforms, tumour cells, cancer-associated fibroblasts (CAFs) and lymphocytes were first localized by marker-based protein staining, after which stroma-encapsulated tumour-cell aggregates were outlined as CMZs (see the corresponding MIBI-TOF and CODEX sections for platform-specific details). Across all platforms, this yielded 296 CMZs in total (Visium HD, Xenium, MIBI-TOF and CODEX CMZs; Supplementary Table 1). The final CMZ regions were used for subsequent subclone modelling at both the cellular and CMZ levels, as well as for spatial topology analysis and inference of tumour evolutionary dynamics.

### Determination of the cell evolutionary root state

To enable objective and reproducible determination of the root state in pseudotime analysis, we developed a multi-criteria strategy. The core concept of Monocle2 pseudotime analysis is to reconstruct a relative temporal ordering of cells based on continuous changes in gene expression patterns.^59^ This analytical paradigm establishes temporal relationships through ordered transitions of state-defining features. Therefore, Monocle2 provides an appropriate computational framework for temporal analysis at the scale of cancer microzones (Cancer Microzone, CMZ).

At the spatial level, CMZs represent locally bounded and relatively independent structural units within tumor tissues. At the lineage level, tumor cells within a CMZ are typically derived from a common ancestral cell, resulting in a relatively homogeneous genetic background and a clear evolutionary relationship. These characteristics effectively reduce confounding from multi-origin or mixed lineage populations, making CMZs suitable units for reliable pseudotime inference using Monocle2.

After CMZ construction, pseudotime analysis was performed separately for tumor cells within each spatial region. The trajectory in low-dimensional space represents continuous relationships among cell states and enables partitioning of cells into discrete states, each corresponding to a group of transcriptionally similar cells. Pseudotime values were then assigned by ordering cells along the trajectory, reflecting their relative positions rather than absolute time. Because the direction of pseudotime depends on the choice of the initial state (root state), determining the root state is a critical step in pseudotime analysis.

To avoid subjective or arbitrary selection of the root state, we implemented a multi-evidence strategy. First, the states identified by Monocle2 were mapped back to their original spatial coordinates, and their spatial distribution was evaluated at the CMZ level. Within each CMZ, states located closer to the microzone center were considered candidate early-stage states, whereas those distributed toward the periphery or outward-expanding regions were considered candidate late-stage states. Second, we quantified the number of tumor cells in each state (measured by the number of spatial barcodes), and considered larger states as supportive evidence for earlier structural stages, while smaller states were regarded as indicative of later stages. By integrating spatial localization and state size, an initial root state was determined for each trajectory.

Following spatial and size-based determination, we further evaluated transcriptional consistency to validate the inferred root state. Specifically, epithelial–mesenchymal transition (EMT)-related gene set scores, stemness-related gene set scores, and differentially expressed genes across states were systematically analyzed to assess whether their variation patterns along the pseudotime trajectory were consistent with spatial organization and state ordering. These analyses were used to evaluate the biological plausibility of pseudotime directionality, rather than as prior criteria for root state determination. By integrating spatial structure, state size, and transcriptional consistency, we determined stable and reproducible root states for Monocle2 pseudotime analysis in each region, providing a robust basis for subsequent temporal modeling and evolutionary analysis.

### Pseudotime trajectory inference based on Monocle2

Pseudotime analysis was performed using Monocle2 (version 2.24.1). First, epithelial cell data from the Seurat object were converted into a Monocle2 CellDataSet object. Data were then normalized and dispersion parameters were estimated to correct for differences in transcript abundance and variability across cells. Genes expressed in at least 10 cells were retained for downstream analysis. To identify genes associated with biological processes, differential expression analysis was performed using cell clustering results as grouping variables. Genes with a false discovery rate (q value < 0.01) were selected as ordering genes for trajectory construction. Dimensionality reduction was then performed using reversed graph embedding, and cells were projected into a two-dimensional space. Based on this embedding, cells were ordered to obtain pseudotime values and partitioned into discrete cell states. Trajectory structures were visualized by coloring cells according to state, pseudotime, or cluster identity. The root state was specified to ensure that the inferred pseudotime direction was consistent with biological interpretation.

### EMT and stemness-related gene set scoring analysis

To systematically evaluate transcriptional program features across different pseudotime states, gene set scoring was performed using the module scoring approach implemented in Seurat (version 5.1.0) (AddModuleScore function).^60^ This method computes the average expression of genes within a target gene set and normalizes it against control genes with similar expression levels, thereby providing robust functional scores at the level of individual spatial units (bins). The epithelial–mesenchymal transition (EMT) program was assessed using the HALLMARK EPITHELIAL MESENCHYMAL TRANSITION gene set from the Molecular Signatures Database (MSigDB) (v2025.1.Hs). The stemness-related gene set was curated based on reported stemness-associated features in human triple-negative breast cancer. After calculating gene set scores for each spatial bin, scores were aggregated and compared at the state level to analyze dynamic changes of EMT and stemness features along the pseudotime trajectory and to evaluate their consistency with state transitions. In addition, differential expression analysis was performed between states to identify state-specific marker genes, thereby providing further validation of state assignments at the transcriptional level.

### State-based differential expression analysis

To further characterize transcriptional differences between pseudotime states and validate the biological relevance of state assignments, differential expression analysis was performed using Seurat (version 5.1.0). For each state, a one-versus-rest comparison strategy was applied, in which the target state was compared against all other states to identify state-specific differentially expressed genes. Differential expression analysis was conducted using the FindMarkers function, based on the non-parametric Wilcoxon rank-sum test. P values were adjusted for multiple testing using the Benjamini–Hochberg method to control the false discovery rate. Genes were selected based on statistical significance (adjusted P < 0.05) and effect size (log fold change threshold) to identify biologically meaningful state-specific genes. The resulting differentially expressed genes were used to characterize transcriptional features of each state and, together with gene set scoring results (EMT and stemness scores), to validate the directionality of the pseudotime trajectory and the ordering of states.

### Cancer Progression Metric: a tissue-embedded spatiotemporal coordinate for quantifying tumour evolution

To characterize the continuous evolutionary dynamics of tumors within the Cancer Microzone (CMZ), we propose a space-time integrated quantitative framework that captures tumor progression as a continuous process rather than a discrete clonal lineage structure. Spatial transcriptomic data represent a snapshot of the tumor tissue’s organizational state at a single time point, in which each CMZ can be regarded as a spatial cell cluster derived from multiple rounds of proliferation of a common ancestral cell. Within this context, each tumor cell constitutes both the minimal structural and evolutionary unit, and single-cell transcriptomic profiles reflect instantaneous state projections rather than true dynamic evolutionary trajectories. This framework therefore integrates transcriptional variation and spatial organization to quantitatively infer the relative evolutionary progression of tumor cells from single-snapshot data.

To integrate transcriptional temporal ordering with spatial progression within CMZs, we introduce the Cancer Progression Metric (CPM) as a unified spatiotemporal metric. CPM provides a mathematical framework that integrates transcriptional variation and spatial organization into a single measure. Rather than directly measuring absolute time, CPM infers the relative evolutionary ordering of cells from their combined transcriptional states and spatial distributions. In this formulation, genetic (transcriptional) variation and spatial positioning are not treated as independent variables but are coupled within a unified metric space, enabling a geometric representation of tumor evolutionary progression.

In this framework, CPM is constructed by integrating pseudotemporal and spatial information. First, the raw global pseudotime *t_i_* for each cell *i* is linearly normalized using Min-Max scaling and mapped to a unified scale in the range of [0, 1], yielding the normalized temporal component *t_i_^global^*:

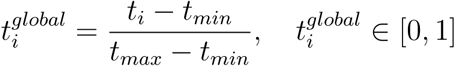

where *t_min_* and *t_max_* represent the minimum and maximum raw pseudotime values across all cells in the sample, respectively.

Considering that solid tumors generally exhibit irregular shapes and direction-dependent growth patterns in tissue space, spatial normalization is performed independently within each CMZ. Specifically, let (*c_x_*, *c_y_*) be the robust geometric center of the CMZ, defined by the median spatial coordinates (*x, y*) of all cells within it. We calculate the spatial position of each cell *i*(*x_i_, y_i_*) relative to this center, and estimate the maximum radial expansion of the CMZ in the corresponding angular direction *θ_i_* to define the directionally normalized spatial component *r_i_^dir^*:

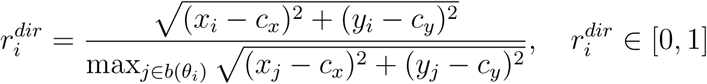

where the numerator denotes the Euclidean distance *d_i_* from cell *i* to the CMZ center, and the denominator represents the maximum radial boundary *R*(*θ_i_*) defined by the furthest cell j within the same angular bin *b*(*θ_i_*). This definition characterizes the cell’s position relative to the growth front in its respective direction, explicitly incorporating directional spatial progression differences.

On this basis, the Cancer Progression Metric is mathematically formulated as a convex combination of multi-modal state functions, integrating the temporal and spatial components:

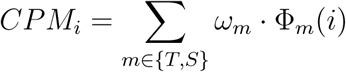

where *m* ∈ {*T, S*} denotes the underlying data modalities representing the transcriptomic/temporal (*T*) and spatial (*S*) features, respectively. The state function Φ*_T_* (*i*) corresponds to the normalized global pseudotime *t_i_^global^*, and *Φ_S_*(*i*) corresponds to the directionally normalized spatial component *r_i_^dir^*. The modal contributions are balanced by the weight vector ω, subject to ∑ω*_m_* = 1. Specifically, we parameterize the transcriptomic weight as ω*_T_* = α and the spatial weight as ω*_S_* = 1 − α. The parameter α thereby controls the relative contribution of transcriptional variation and spatial positioning in the metric construction. To determine its optimal value, we employed a Spatiotemporal Trade-off Calibration strategy. This approach systematically evaluates a predefined grid of *α* values to quantify the balance between molecular fidelity and spatial topological constraints, precisely isolating the equilibrium (*α* = 0.85) that ensures highly robust macroscopic evolutionary dynamics. The resulting CPM provides a unified quantitative measure of the relative evolutionary state of tumor cells within a single time snapshot, rather than representing their progression along absolute time.

### A spatiotemporal evolutionary dynamics model: Field–Flow–Front (3F)

To characterize the spatial organization of tumor evolution within single time-slice spatial transcriptomic data, we further extend CPM from a relative process coordinate at the single-cell level to a continuous evolutionary description defined in physical space. Based on this modeling approach, tumor evolution is not described as a discrete clonal lineage but rather as a continuous evolutionary process embedded in the tissue space, with its states, directions of change, and active regions being characterized within a unified framework. In this context, we construct the Field–Flow–Front (3F) model, which formalizes the continuous evolutionary dynamics of tumors in space by combining evolutionary state fields, directional evolutionary flows, and evolutionary fronts.

Field In the 3F model, CPM is viewed as a continuous scalar field defined in physical space, used to characterize the relative evolutionary state heights of cells in space. Formally, the evolutionary state field is defined as:

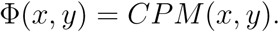

This scalar field assigns a continuous evolutionary state value to each cell in tissue space, providing the basis for subsequent quantification of spatial dynamics.

Flow To characterize the directional changes in evolutionary state within space, we construct a k-nearest neighbors (KNN) graph based on the spatial coordinates of cells. On this adjacency structure, for each cell i, we calculate the average absolute difference between its CPM and the CPMs of its neighboring cells, which estimates the local evolutionary field gradient intensity (i.e., the evolutionary slope):

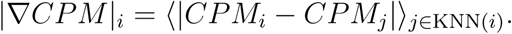

This measure characterizes the magnitude of evolutionary state changes within local spatial neighborhoods.

On this basis, to describe the directional changes of evolutionary states within space, we project the CPM differences onto the normalized spatial displacement direction and define the evolutionary flow vector for each cell *v⃗_i_* = (*v_x_*, *v_y_*) as:

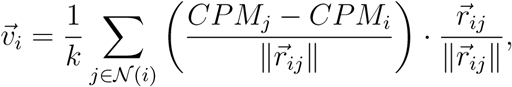

where *r⃗_ij_* represents the spatial displacement vector between cell i and its neighbor j. The magnitude of this vector characterizes the intensity of local evolutionary changes, while its direction indicates the main direction of evolutionary state changes in space.

Front The evolutionary front is defined as the spatial region where the evolutionary state is leading or undergoing the most significant changes within the evolutionary field. Specifically, when a cell’s CPM value or its local evolutionary slope |∇CPM| exceeds the 85th percentile of the distribution in that region, the cell is marked as part of the evolutionary front. This criterion is used to identify spatial regions with leading evolutionary stages and local areas undergoing the most rapid evolutionary changes.

Subclone Stratification Constrained by CPM Isobands In the evolutionary field model (EFM) framework, subclones are defined as stable regions in the evolutionary potential field that are spatially coherent. To achieve automatic subclone stratification, we apply the optimal k-means clustering algorithm (Ckmeans.1d.dp) to the CPM values for data-driven grouping and determine the optimal stratification result by the curvature of the within-group sum of squares curve, resulting in integer-labeled CPM isobands. Each CPM isoband corresponds to a finite-width evolutionary potential interval, representing a relatively stable evolutionary stage. Further, within each CPM isoband, we identify spatially connected components based on the spatial adjacency relationships between cells and define each spatially connected isoband region as an independent evolutionary subclone. This definition ensures the consistency of subclones in both evolutionary state and spatial structure.

Comparative Evolutionary Dynamics of BMZ and CMZ To further investigate the macro-environmental influences on tumor evolution, we utilized the 3F framework to compare the evolutionary dynamics between the BMZ and CMZ. Single-cell 3F metrics were extracted and aggregated across predefined regions belonging to these two macro-environments. We systematically evaluated differences in four core evolutionary indicators: (1) the overall evolutionary state (CPM), (2) the local evolutionary gradient (CPM slope), (3) the global evolutionary flow magnitude, and (4) the localized driving power of the evolutionary front (quantified as the flow magnitude exclusively within cells identified as the evolutionary front). To ensure robust quantification and mitigate the effect of extreme outliers, flow magnitudes were truncated at the 99th percentile prior to analysis. Statistical comparisons between the BMZ and CMZ for all continuous metrics were performed using the Wilcoxon rank-sum test.

In summary, the 3F model provides a systematic analytical framework for characterizing the continuous evolutionary dynamics of tumors in spatial transcriptomic data from a single time snapshot by integrating evolutionary state fields, directional evolutionary flows, and evolutionary fronts.

### Sensitivity analysis of the weighting parameter α

To evaluate the robustness of CPM with respect to the weighting parameter *α*, we systematically varied *α* across a predefined grid {0.50, 0.60, 0.70, 0.75, 0.80, 0.85, 0.90, 0.95} and recomputed CPM independently for all 49 regions. For each *α* value, four quantitative metrics were assessed: (i) the Spearman correlation between CPM and normalized pseudotime ( *ρ*_CPM,pt_), reflecting the fidelity of temporal encoding; (ii) the Spearman correlation between CPM and the direction-normalized spatial progression ( *ρ*_CPM,*r*_*_dir_*), reflecting the retention of spatial information; (iii) the mean evolutionary flow magnitude at the front (*v̄*_front_), capturing local evolutionary activity at the leading edge; and (iv) the front load, defined as the proportion of cells classified as evolutionary front cells.

Across all regions, *ρ*_CPM,pt_ increased monotonically with *α*, reaching 0.972 ± SD at *α* = 0.85, while *ρ*_CPM,*r*_*_dir_* decreased monotonically, retaining a value of approximately 0.49 at the same point. This monotonic divergence is an analytically expected property of any convex combination of the two components and does not by itself define an optimum. Critically, front load and *v̄*_front_ exhibited only marginal variation across the full *α* range and preserved their inter-regional ordering, indicating that the identification of the evolutionary front and the downstream dynamics derived from it are insensitive to the precise weighting. The mean and standard deviation of all four metrics across regions are reported as a function of *α* in Fig. 2b. We therefore interpret this analysis as establishing robustness rather than identifying an optimum: because the downstream metrics are stable across [0.5, 0.95], the principal conclusions of this study do not depend on a particular value of *α*. On this basis we fixed *α* = 0.85 as a representative setting for all downstream analyses, a value that keeps the molecular clock as the dominant determinant of ordering while retaining spatial position as an auxiliary directional constraint. The high CPM–pseudotime correlation at this setting ( *ρ* ≈ 0.97) reflects this intended design: the spatial term contributes a small, directionally informative correction rather than a wholesale reordering, consistent with its role as a regularizer rather than a co-equal coordinate.

### Pseudotime normalization and directional spatial standardization

After obtaining pseudotime and spatial coordinates for each cell, pseudotime values were linearly rescaled to the interval [0, 1] using min–max normalization, *t_i_^global^* = (*t_i_* −*t_min_*)/(*t_max_* − *t_min_*), where *t_min_* and *t_max_* denote the minimum and maximum pseudotime values within each region. This transformation removes regional differences in pseudotime range and ensures comparability across regions for subsequent analyses. Normalization was performed independently within each region. Regions with zero pseudotime range or containing non-finite values (for example NA or Inf) were excluded from downstream analyses.

Directional spatial standardization was performed independently within each spatial group (for example CMZs or other microzone units defined within a region). To improve robustness to irregular shapes and local outliers, the median spatial coordinates of cells within each group were used as the centre, and the angular position of each cell relative to this centre was calculated. The angular space [−π, π] was discretized into fixed bins (72 bins in the implementation), and within each angular bin the maximal radial extent was estimated as the directional scale. Each cell’s radial distance was subsequently normalized by this directional scale, producing a standardized radial coordinate that preserves directional progression (consistent with the definition used in the CPM construction step). Groups containing fewer than 10 cells or lacking a defined directional scale were excluded from directional normalization and subsequent analyses.

### Construction of K-nearest neighbor graph and computation of local evolutionary quantities

To robustly estimate local spatial variation from discretely sampled cells, a k-nearest neighbor (KNN) graph was constructed based on two-dimensional spatial coordinates (with k = 10 in the implementation), using the FNN package to identify local neighborhoods. Regions with insufficient cell numbers to support KNN construction (n ≤ k) were excluded from further analysis. Within the KNN neighborhood, we computed (i) a local variation magnitude metric by aggregating the absolute differences in CPM values between each cell and its neighbors and taking the mean, serving as an estimate of local evolutionary variation (consistent with the local slope definition in the 3F model, but emphasized here in terms of numerical implementation); and (ii) a local directional variation vector by combining spatial displacement directions with CPM differences and averaging across the k neighbors to obtain a cell-level vector. To avoid numerical instability caused by zero distances, a small constant (10^−6^ in the implementation) was added to the distance term. The magnitude of the resulting vector was used to quantify flow strength and was further applied in visualization filtering (e.g., displaying only vectors above a high quantile threshold and applying random subsampling to control arrow density).

### CPM isoband stratification using Ckmeans.1d.dp

To achieve automatic stratification of CPM isobands, we applied one-dimensional optimal k-means clustering (Ckmeans.1d.dp) to finite CPM values.^61^ To avoid subjective selection of the cluster number, models were fitted across a predefined range (*k* = 1–7 in the implementation), and the within-cluster sum of squares (SSE) was calculated for each k:

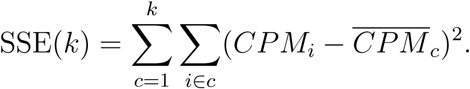

The optimal k was determined from the curvature of the SSE curve using second-order differences, Δ^2^SSE(*k*) = SSE(*k* + 1) − 2SSE(*k*) + SSE(*k* − 1), and selecting the value corresponding to the most pronounced curvature change (elbow point). The resulting clustering produced integer-labelled CPM isobands (CPM bands), which were subsequently used to identify spatially connected components and define subclones. Non-finite CPM values were excluded from clustering and assignment.

### Spatial trajectory inference using stLearn

To highlight the advantages of our CPM in resolving continuous spatial trajectories, we contrasted our model against conventional discrete inference using the Python package stLearn (v1.4.0).^27^ We emphasize that the stLearn diffusion pseudotime is an external, independently computed ordering, distinct from the Monocle2 global pseudotime that serves as the internal transcriptional clock of CPM; it therefore provides an unbiased reference against which to benchmark CPM. The standard stLearn workflow was applied independently to each delineated CMZ. First, Spatial Morphological gene Expression (SME) normalization was performed to integrate spatial coordinates, matched H&E morphological features, and transcriptomic profiles, thereby smoothing the gene expression data based on tissue architecture. Subsequently, dimensionality reduction (PCA) and neighbourhood graph construction were executed, followed by Louvain clustering to partition the cells into discrete spatial sub-clusters. For trajectory inference, Partition-based Graph Abstraction (PAGA) was utilized to construct the global connectivity network among these discrete clusters. Diffusion Pseudotime (DPT) was then calculated to infer the evolutionary progression. Crucially, to ensure a rigorous and fair quantitative comparison with CPM, the root cell for stLearn’s DPT calculation was assigned to the cluster corresponding to the ancestral root region identified in our spatially informed ordering strategy.

For comparative analysis, the resulting DPT values and discrete cluster assignments for each spot were extracted alongside the corresponding CPM values. Two complementary contrasts were quantified. First, within individual stLearn clusters we computed the CPM range spanned by their constituent cells, where a broad spread (for example ΔCPM > 0.5 within a single cluster) indicates continuous micro-evolutionary heterogeneity that discrete clustering collapses into an apparently homogeneous state. Second, across regions we compared the dynamic range of CPM against that of stLearn DPT, identifying cells compressed into a narrow DPT interval yet spanning a wide range of CPM as evidence that CPM retains discriminative resolution where graph-based pseudotime saturates.

### Reconstruction of region-level subclones via 3F-guided spatial clustering

To characterize tumour heterogeneity at the macro-architectural level, we extended the 3F framework from single cells to the cancer microzone (CMZ) level. Each CMZ was treated as a discrete evolutionary meta-unit, and a Spatial–Evolutionary Graph (SEG) clustering framework was developed to identify region-level subclones exhibiting both spatial connectivity and evolutionary synchrony.

#### Construction of CMZ evolutionary fingerprints

For each CMZ *C_k_* comprising *N_k_* cells, we extracted a 3F evolutionary fingerprint vector 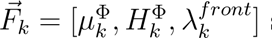 summarizing its dynamic state across three dimensions. Field (macro-evolutionary state, 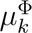) characterizes the consensus evolutionary stage of the CMZ. Given that CPM is globally comparable within a region, the median CPM was used to represent the core evolutionary position of the CMZ, 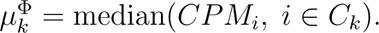

Flow (evolutionary entropy, 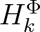) quantifies intra-CMZ evolutionary heterogeneity. The Shannon entropy of the CPM distribution was calculated as

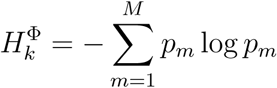

where the CPM range was discretized into M = 20 bins and *p_m_* denotes the probability of cells falling into the mth bin.

Front (front load, 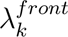) represents the expansion activity of the CMZ and was defined as the proportion of cells classified as evolutionary front cells, 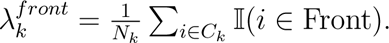

Prior to graph construction, the feature vector *F⃗_k_*was Z-score standardized to ensure equal contribution from all three evolutionary dimensions.

#### Spatial–evolutionary graph construction and clustering

We constructed an undirected graph *G* = (*V, E*) in which nodes represent CMZ centroids. Spatial adjacency was defined using a k-nearest neighbour algorithm (*k* = 6) based on physical coordinates. To incorporate evolutionary similarity, edges were weighted according to the similarity between 3F fingerprints of neighbouring CMZs using a Gaussian kernel

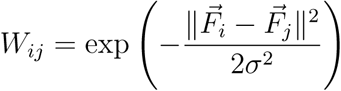

with bandwidth parameter σ = 0.5.

Region-level subclones were then identified using the Louvain community detection algorithm,^62^ which partitions the graph by maximizing modularity and thereby delineates spatially coherent tissue domains governed by shared evolutionary dynamics without pre-specifying the number of clusters.

### Z-score standardization of CMZ features

To eliminate scale differences among evolutionary features, feature vectors at the CMZ level were standardized prior to the construction of the spatial–evolutionary graph. For each feature variable *x*, Z-score normalization was applied as *z* = (*x* − µ)/σ, where µ and σ denote the mean and standard deviation of the feature across all CMZs, respectively. In implementation, macro-evolutionary state (Median CPM), evolutionary entropy (Entropy), and front load (Front Load) were standardized independently. This ensures that different feature dimensions contribute on a comparable scale in subsequent distance calculations, preventing any single feature from dominating the clustering results.

### CMZ-level subclone identification based on the Louvain algorithm

To identify CMZ-level subclones on the spatial–evolutionary graph, the Louvain algorithm was applied for community detection.^62^ This method partitions the network by maximizing modularity, thereby identifying groups of nodes that are densely connected and exhibit high internal similarity. Modularity is defined as:

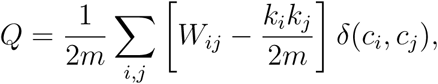

where *W_ij_* denotes the edge weight, *k_i_* and *k_j_* are the weighted degrees of nodes, m is the total sum of edge weights in the graph, and δ(*c_i_*, *c_j_*) is an indicator function (equal to 1 if two nodes belong to the same community, and 0 otherwise). In implementation, modularity optimization was performed on a weighted undirected graph, and random initialization was used to ensure reproducibility. The resulting communities correspond to region-level subclones that are spatially contiguous and evolutionarily coherent.

### Evolutionary fitness index-based local clonal competition analysis

To quantify local competitive interactions between cancer microzone (CMZ)-level subclones, we performed a two-dimensional competition analysis based on the Region Subclone assignments generated by the 3F spatial clustering procedure. Cell-level outputs from each tissue region were aggregated at the Region Subclone level. For each subclone, we calculated centroid coordinates, cell number, median Cancer Progression score (Macro-CPM), front load, Cancer Progression entropy, mean flow magnitude, and directional coherence. Front load was defined as the fraction of cells annotated as front cells. Cancer Progression entropy was calculated as the Shannon entropy of CPM values after binning CPM into 20 equal-width intervals over the range [0,1]. Directional coherence was defined as the magnitude of the mean flow vector divided by the mean flow magnitude, 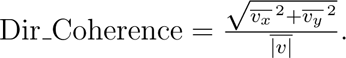

To summarize the local evolutionary advantage of each subclone, we defined an evolutionary fitness index (EFI). For each feature, values were standardized by z-score across all subclones. EFI was then calculated as a weighted linear combination of Macro-CPM, front load, mean flow magnitude, and directional coherence: EFI = 0.35 Z(Macro-CPM)+0.35 Z(Front Load)+ 0.20 Z(Flow Magnitude) + 0.10 Z(Dir Coherence). When flow fields were unavailable, EFI was reduced to an equally weighted combination of Macro-CPM and front load. CPM entropy was retained as an additional descriptive variable but was not included in the main EFI definition. To infer local neighbourhood relationships between subclones within each region, we constructed a centroid-based k-nearest-neighbour (KNN) graph using subclone centroids in two-dimensional tissue space. For each subclone, edges were drawn to its three nearest neighbouring subclones (or all available neighbours when fewer than three existed), and duplicated edges were removed. To avoid biologically implausible long-range connections, edges with distances above the 90th percentile within each region were excluded. The resulting graph was used as the local competition network.

For each neighbouring subclone pair, competitive outcome was determined using a hierarchical rule intended to minimize circularity with EFI. The subclone with higher front load was assigned as the winner; ties were resolved first by higher Macro-CPM and then by higher mean flow magnitude. For each pair, we calculated the EFI difference (ΔEFI), the absolute EFI difference, and a boundary sharpness score defined as the absolute Macro-CPM difference divided by centroid distance, 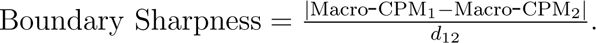

Neighbouring pairs were further stratified into near and far groups within each region using the region-specific median centroid distance as the cutoff. To evaluate whether EFI predicts local dominance, we defined a binary variable indicating whether the higher-EFI member of each pair was also the winner. This variable was analysed using logistic regression with absolute EFI difference, distance group, and their interaction as predictors. Boundary sharpness was analysed by linear regression against absolute EFI difference and centroid distance. In addition, paired Wilcoxon signed-rank tests were used to compare EFI, Macro-CPM, front load, entropy, flow magnitude, and directional coherence between winners and losers. Spearman correlation analyses were used to assess the relationships between EFI and front load, Macro-CPM, entropy, flow magnitude, and directional coherence. For visualization, EFI landscapes were generated by plotting subclone centroids in tissue space with node colour representing EFI and node size representing subclone size. Competition networks were plotted using the same centroids, with edge width proportional to the absolute EFI difference between neighbouring subclones.

### Image registration and tumor region segmentation

To validate whether region-level subclones identified by the 3F model correspond to physically real spatial structures, we constructed a three-dimensional topological model of cancer microzones (CMZs) based on consecutive H&E-stained histological sections. After digitization of serial FFPE sections, the original slide files were stored in the MRXS format of CaseViewer and subsequently batch-converted into TIFF format using 3DHISTECH Slide Converter. Each section was then manually cropped through interactive selection to ensure that comparable tumor regions were retained across adjacent slices. A multi-stage registration pipeline was applied to reconstruct the volumetric tissue architecture. First, coarse alignment was performed using manual rigid transformations (rotation and translation) on a large unified canvas (with consistent global dimensions and white background padding) to correct for global positional variations introduced during slide scanning. Second, the coarse-aligned images were refined using sequential automatic registration based on the enhanced correlation coefficient (ECC) algorithm under Euclidean motion constraints (MOTION EUCLIDEAN), which estimates the optimal transformation between adjacent sections.^63^ The ECC algorithm aligns images by maximizing the normalized correlation of grayscale intensities between the reference and moving images, and the optimization objective can be formulated as:

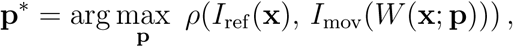

where *I*_ref_ and *I*_mov_ denote the grayscale images of adjacent sections, and W (·) represents a Euclidean rigid transformation (rotation and translation).

To improve robustness against staining variation and local noise, preprocessing steps including grayscale conversion, background inversion, and Gaussian smoothing were applied prior to ECC estimation, allowing the optimization to focus on tissue contours and global structures rather than local color differences. Registration was performed sequentially using adjacent sections as references (sequential registration), and the ECC correlation coefficient was recorded as a quality metric. When the correlation coefficient was low or the optimization diverged, the coarse alignment result was retained to prevent propagation of misregistration. After alignment, the registered sections were imported into QuPath for tumor region annotation.^64^ Binary segmentation masks were generated from manually delineated Tumor annotation objects on a per-slice basis and stacked in a unified coordinate system to form a voxel-based tumor volume, which was subsequently used for three-dimensional topological modeling and CMZ identification.

### Graph-based topological modeling and CMZ identification

To quantitatively identify physically independent CMZ structures in three-dimensional space, we constructed a directed acyclic graph (DAG) model G = (V, E) from the stacked tumor volume masks, where nodes represent connected tumor regions within each slice and edges represent inter-slice continuity. Model parameters were independently optimized for each sample to account for variations in slice resolution, inter-slice spacing, and tissue heterogeneity. (1) Node definition: For each slice z, connected component analysis (8-connectivity) was performed on the binary mask. Each connected component corresponds to a candidate node v ∈ V, with its area and centroid coordinates computed. To remove nonspecific staining and digitization noise, a sample-specific minimum area threshold *A*_min_ was applied to exclude small fragments. (2) Edge construction: To model tumor continuity along the sectioning axis, edges were only allowed from layer *z* to *z* + 1, ensuring an acyclic graph structure. For nodes *u* (at layer *z*) and *v* (at layer *z* + 1), a directed edge (*u, v*) ∈ E was established if either of the following criteria was satisfied: (i) morphological overlap criterion, where the mask of node u was morphologically dilated to form an envelope *D*(*u*), and an edge was created if *D*(*u*) overlapped with the mask of node v; the dilation kernel size was adaptively determined based on residual registration error tolerance; (ii) centroid proximity criterion, where an edge was established if the Euclidean distance between centroids was smaller than a search radius R, which was adjusted according to slice thickness and local tumor cell density. To quantify the confidence of inter-slice continuity, edge weights could be defined as a distance-decay function (e.g., *w_uv_* = 1/(1 + *d_uv_*)) for downstream analysis. (3) CMZ identification: In the constructed DAG, connected components in the undirected version of the graph were defined as candidate three-dimensional structural units. To eliminate spurious connections caused by incidental overlap or transient contact, only connected subgraphs spanning more than a minimum number of slices *S*_min_ (set to *S*_min_ = 7 in this study) were retained as valid physical CMZs. Additional features, including layer span, total volume (summed area across slices), and splitting/fusion events (based on nodes with in-degree or out-degree greater than one), were computed to characterize the three-dimensional topological complexity of each CMZ.

### Three-dimensional surface reconstruction and geometric analysis (Marching Cubes)

For each topologically validated CMZ, three-dimensional surface geometry was reconstructed from its voxelized mask to assess whether 3F-defined subclones correspond to spatially discrete architectural units within the physical tumor volume. Specifically, the binary volumetric representation of each CMZ was treated as a scalar field, and an isosurface was extracted at the threshold level of 0.5 using the Marching Cubes algorithm to generate a triangular mesh representation.^65^ To correct for inconsistencies between inter-slice spacing and in-plane resolution, a scaling factor was applied along the z-axis during reconstruction (e.g., based on slice thickness or empirically defined z-stretch), ensuring that the reconstructed geometry reflects the true spatial proportions of the sampled tissue. The resulting triangular meshes were used for three-dimensional visualization and quantitative analysis, and were further compared with region-level subclones identified by the 3F model in spatial transcriptomics data to validate their spatial correspondence.

### Subclone markers, differential gene annotation, functional program scoring, and CPM-band trends

Within each spatial region (region), we performed biological characterization by grouping cells into six subclones (g1–g6) defined by CPM bands. Subclone markers and differentially expressed genes (DEGs) were identified at the bin level using the Seurat framework for differential expression testing (for example, one-vs-rest or specified pairwise comparisons), followed by FDR correction for multiple testing. Marker heatmaps for visualization were generated based on the average expression within each subclone and further standardized by gene (e.g., z-score) to highlight relative expression modules. Functional annotation of significant DEGs was performed using GO/KEGG enrichment analysis (with FDR correction), with the background gene set defined as the set of genes detectable in the expression matrix of the corresponding region.^66, 67^ To enable unified comparisons of tumor functional states across subclone, flow, front, and advantageous subclones, we used six core functional program scores: inflammatory response (Inflammatory), metabolism (Metabolism), proliferation (Proliferation), stress response (Stress), EMT, and stemness (Stemness). Gene sets for Inflammatory/Metabolism/Proliferation/Stress/EMT were derived from MSigDB Hallmark modules (imported in GMT format), whereas the Stemness gene set was obtained from our previously constructed stemness-related gene set; all gene symbols were standardized, and only genes detectable in the expression matrix were retained for downstream analysis. The mapping from Hallmark gene sets to functional modules was as follows: **Metabolism:** HALLMARK CHOLESTEROL HOMEOSTASIS, HALLMARK FATTY ACID METABOLISM, HALLMARK GLYCOLYSIS; **Proliferation:** HALLMARK DNA REPAIR, HALLMARK E2F TARGETS (if HALLMARK G2M CHECKPOINT and HALLMARK MITOTIC SPINDLE are additionally used, they are explicitly incorporated into the proliferation/cell cycle axis); **Stress:** HALLMARK UNFOLDED PROTEIN RESPONSE, HALLMARK REACTIVE OXYGEN SPECIES PATHWAY, HALLMARK HYPOXIA; **Inflammatory:** HALLMARK ALLOGRAFT REJECTION, HALLMARK TNFA SIGNALING VIA NFKB, HALLMARK COMPLEMENT, HALLMARK IL2 STAT5 SIGNALING, HALLMARK IL6 JAK STAT3 SIGNALING, HALLMARK INFLAMMATORY RESPONSE, HALLMARK INTERFERON ALPHA RESPONSE, HALLMARK INTERFERON GAMMA RESPONSE; **EMT:** HALLMARK EPITHELIAL MESENCHYMAL TRANSITION. At the bin level, we used Seu-rat::AddModuleScore to compute scores for the six programs, and aggregated them within each region by CPM-band to obtain subclone-level trajectories. Monotonic trends of functional programs along clonal evolution were assessed using Spearman rank correlation (correlation coefficient *ρ* and p value between program scores and CPM-band). To derive cross-region consensus trajectories, we averaged CPM-band program scores across regions, performed z-score standardization for each program, normalized them into relative contribution proportions, and visualized them as stacked area plots (river plots) to depict continuous functional reorganization along CPM bands.

### Dynamic coupling analysis of subclones

To quantitatively characterize the coupling between subclone stages and dynamic/front phenotypes, we summarized the cell-level table output from EFM (including CPM, CPM slope, flow vx, flow vy, flow mag, is front, etc.) across region1–6. For each region × CPM-band, we computed the number of bins *n*_cells_, front proportion front ratio, mean CPM and CPM slope, mean flow mag, and mean flow vector components mean vx and mean vy. To quantify directional consistency, we defined

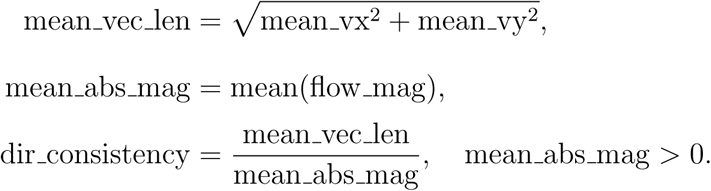

In addition, within each region, CPM-bands were classified into Type I–III based on quantile thresholds of mean flow mag and front ratio (for example, using the 0.75 quantile of mean flow mag as the high-flow threshold and combining the 0.75/0.50 quantiles of front ratio to define Type III/II/I), enabling comparisons of front participation, flow strength, and directional coordination across subclone stages within a unified coordinate system, and supporting visualization of front ratio trajectories/heatmaps, flow metric trajectories, joint distributions of front ratio and flow strength, and CPM-band mean flow vector maps.

### Flow dynamics metrics and phase diagram construction (magnitude, coherence, heterogeneity)

At the CMZ scale, we summarized local evolutionary dynamics using bin-level flow vectors (flow vx, flow vy) and their magnitudes (flow mag) from EFM output. Non-finite values of flow vx, flow vy, and flow mag were first filtered. CMZ was used as the grouping unit (CMZ labels obtained from the input data), and the number of bins *n*_cells_ and spatial center coordinates (cmz x, cmz y) were computed. Summary metrics included

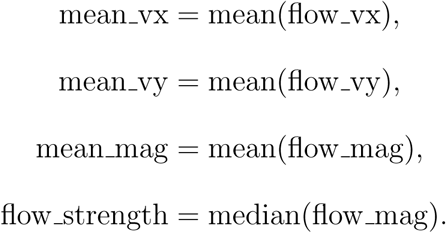

Directional coherence was defined as

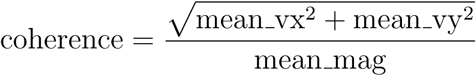

and internal heterogeneity was defined as

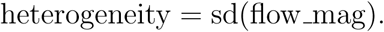

which quantifies directional consistency within CMZ; internal heterogeneity was defined as heterogeneity = sd(flow mag), reflecting dispersion of flow magnitude within CMZ. Based on these metrics, three visualizations were constructed: (1) spatial projection of CMZ mean flow vectors, plotting arrows at (cmz x, cmz y) with direction (mean vx, mean vy) and width mapped to mean mag; (2) CMZ dynamic phase diagram, with coherence on the x-axis and heterogeneity on the y-axis, point size mapped to *n*_cells_, and color mapped to flow strength; (3) CMZ evolutionary activity map, mapping flow strength back to bins within each CMZ and projecting onto tissue coordinates to reveal the structured spatial distribution of CMZ activity.

### Flow-aligned functional asymmetry (Down–Up)

To test whether functional programs exhibit directional organization along CPM-defined evolutionary flow, we computed flow-aligned asymmetry (Down–Up). Within each region, for each bin, a K-nearest neighbour local neighbourhood was constructed based on two-dimensional spatial coordinates. Let the local evolutionary flow vector be *v⃗_i_* with unit direction *v̂_i_* = *v⃗_i_*/ ∥ *v⃗_i_*∥. For any neighbouring bin *j*, the displacement vector Δx*_ij_* = (x*_j_* − *x_i_*, *y_j_* − *y_i_*) was projected onto the flow direction as *s_ij_* = Δx*_ij_* · v̂*_i_*, whereby neighbours with *s_ij_* > 0 were defined as downstream and those with *s_ij_* < 0 as upstream. For a functional program score f, downstream and upstream neighbourhood means 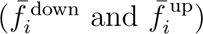 were calculated, and bin-level asymmetry was defined as

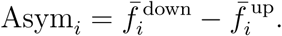

Averaging Asym*_i_* across bins yielded the region-level directional bias of each functional program along the evolutionary flow. Cross-region comparisons were then performed to assess the consistency of upstream–downstream functional asymmetry.

### Molecular characterization of front (DEG/enrichment/PPI; functional program comparison)

The definition and computation of front and Front-load have been described in the 3F model section. Here we focus on their biological characterization. Within each region, we compared differentially expressed genes (DEGs) between front and non-front cells (with FDR correction), and performed GO/KEGG enrichment analysis (with FDR correction) for significantly upregulated or downregulated genes, respectively. Interaction networks of front-upregulated genes were constructed using publicly available PPI databases to identify hub modules at the invasive front.^68^ In parallel, we compared the scores of six functional programs between front and non-front cells in a region-stratified manner, and visualized the differences using distribution plots (e.g., raincloud plots or violin plots combined with scatter points), thereby providing cross-region comparable evidence at the program level.

### Definition and spatial mapping of Front Molecular Heterogeneity Index

To quantify molecular heterogeneity at the tumor front at the CMZ scale, we constructed the Front Molecular Heterogeneity Index (FMHI), integrating three aspects: expression complexity, functional diversity, and population structural dispersion. At the single-bin level, three heterogeneity metrics were computed: (1) expression entropy, defined by constructing a probability distribution based on highly variable genes (HVGs) and calculating Shannon entropy; (2) pathway diversity, defined by constructing a distribution of functional program activities based on module scores and calculating Shannon entropy; (3) PC dispersion, defined as the mean squared deviation of bins within front CMZs in the space of the first 20 principal components. The three metrics were averaged at the CMZ level, followed by z-score standardization within each region; FMHI was defined as the arithmetic mean of the three standardized metrics. FMHI was then mapped back to tissue space to identify CMZ hotspots with high FMHI and to characterize the spatial organization of front heterogeneity.

### Identification of advantageous subclones and downstream analysis

To delineate spatial cell populations with selective evolutionary advantage, three evolutionary metrics were computed for all spatial barcode cells within each region: CPM position, local gradient slope along the CPM direction (CPM slope), and the magnitude of local evolutionary flow (flow mag), defined as flow mag = flow vx^2^ + flow vy^2^. Within each region, cells were classified as advantageous if their values of CPM, CPM slope and flow mag were all greater than or equal to the 0.85 quantile of the respective distributions.

Advantageous cells were subsequently stratified by CPM-band. For each CPM-band, a k-nearest-neighbour spatial graph (*k* = 5) was constructed using two-dimensional spatial coordinates (x, y), and spatially contiguous clusters were identified by connectivity analysis. Each connected component was considered a candidate advantageous subclone; clusters containing at least three advantageous cells were retained and labelled as advantageous subclones with identifiers in the format “B*band* A*index* ”.

For downstream biological analysis, differential expression between advantageous and non-advantageous populations was computed within each region (FDR correction applied). Gene ontology (GO) and KEGG enrichment analyses were performed on significantly upregulated genes to annotate functional characteristics of the advantageous state. Protein–protein interaction (PPI) networks were constructed using significantly upregulated genes, and hub modules were identified based on node degree.

To quantify functional biases associated with advantageous states, we calculated the difference in mean program activity between advantageous and non-advantageous populations as Δmean = mean(Advantage) − mean(Non-advantage). These values were summarized across regions as heatmaps to assess cross-region consistency and regional variation of functional programs in advantageous subclones.

### GO/KEGG enrichment analysis

At each region × subclone (or front vs non-front comparison) level, functional enrichment analysis was performed on differential expression results. Genes for enrichment were first selected from DEG tables: by default, genes with FDR (p val adj < 0.05) were used; when the number of significant genes was insufficient to support enrichment (threshold set to 3), the TopN genes ranked by ascending FDR (N=50) were used as a fallback set, and gene sets could be constructed separately for upregulated and downregulated genes when log2FC was available. Gene symbols (SYMBOL) were then mapped to ENTREZID (org.Hs.eg.db)(v3.22.0), and only successfully mapped genes were retained for downstream analysis. GO enrichment was performed using clusterProfiler(v4.18.4)::enrichGO, covering BP/CC/MF (ont = “ALL”), with multiple testing correction by the Benjamini–Hochberg method; to reduce redundancy, semantic similarity-based simplification was applied when feasible (simplify, Wang measure, cutoff=0.6, selecting representative terms by p.adjust). KEGG enrichment was performed using clusterProfiler::enrichKEGG (organism = “hsa”), with multiple testing control applied similarly. Enrichment results were exported as tables and visualized using dotplots for significant terms.

### Spearman rank correlation for CPM-band trend assessment

To evaluate monotonic trends of functional program scores along CPM-band, correlation analysis was performed within each region using CPM-band-aggregated program scores. Specifically, region-level CPM-band mean tables were used, treating CPM-band as an ordered numeric variable, and Spearman rank correlation was computed between each functional program score (columns denoted as * Score) and CPM-band. Correlation analysis was implemented using stats::cor.test (method = “spearman”), yielding the Spearman correlation coefficient *ρ* and corresponding p value, and generating independent trend result tables for each region × program. For visualization, CPM-band–program score trajectories were converted into long-format tables, and *ρ* values (with significance annotations) were displayed in the legends of multi-region overlaid trajectory plots.

### PPI network construction and visualization

PPI networks were constructed using filtered differentially expressed genes as input. Within each region, front-upregulated genes (avg log2FC > 0) were first extracted from front vs non-front DEG results and ranked by adjusted P value (p val adj) to generate input gene lists. These gene lists were then submitted to the STRING database to obtain protein interaction edge tables, which were exported in TSV format. The exported interaction tables were imported into Cytoscape to construct undirected networks, and DEG statistics (e.g., p val adj, significance labels, and log2FC when available) were mapped to node attributes. Network topology metrics were computed at the node level, with node size mapped to degree to reflect hubness, and node color encoding statistical significance (e.g., stratified or continuous mapping based on p val adj). Final network layouts and visual parameters were interactively adjusted in Cytoscape to generate PPI network figures for presentation.

### Protein-level validation of the 3F model using MIBI-TOF imaging

To validate the 3F model at the protein level on an orthogonal platform, we analysed a publicly available MIBI-TOF imaging dataset of triple-negative breast cancer. Single-cell expression matrices with spatial coordinates and cell-type annotations (tumour, CAF, and CD8^+^ T cells) were obtained, and pixel coordinates were converted to physical units(0.39 µm/pixel).

CMZs were delineated manually in Mantis Viewer. For each sample, segmented tumour cells were overlaid on the Pan-Keratin channel, and spatially contiguous, stroma-encapsulated tumour-cell aggregates were outlined as individual CMZs following the same criteria used for the Visium HD data(clear epithelial identity, local independence, and well-defined boundaries). In total, 59 CMZs were delineated across 32 samples and used for downstream analysis.

Within each CMZ, a protein-based Cancer Progression Metric (CPM-pp) was constructed as a convex combination of an EMT–Wnt invasion proxy and the direction-normalized spatial coordinate, 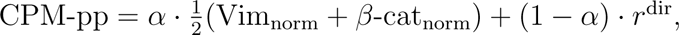 where Vimentin and Beta-catenin intensities were min–max normalized within each CMZ as a protein surrogate for transcriptomic pseudotime, *r*^dir^ was computed as in the Visium HD analysis (radial distance to the CMZ centroid normalized by the maximal radius within 36 angular bins), and *α* = 0.85 was retained from the primary model. Evolutionary flow vectors and flow magnitude were derived from local CPM-pp gradients over a k-nearest-neighbour graph (*k* = 10), and front cells were defined as tumour cells exceeding the 85th percentile of either CPM-pp or local CPM-pp slope within each CMZ. The distance from each tumour cell to its nearest CAF was computed per sample, and tumour cells were classified as CAF-near (< 30 µm) or CAF-far.

To avoid circular validation, statistical comparisons between front and non-front cells were performed using markers independent of the CPM-pp construction—Ki67, SMA, and flow magnitude—using two-sided Wilcoxon rank-sum tests; Vimentin and Beta-catenin were used only as a signature-consistency check. CMZs containing fewer than 11 cells were excluded from flow and front analyses.

### Evolutionary Driver Gene identification based on GeneSwitch kinetics

To elucidate the dominant molecular forces driving tumour evolutionary progression from a single spatial transcriptomics snapshot and to provide biologically grounded validation, we developed a stepwise analytical framework spanning from a continuous evolutionary coordinate system to driver identification, perturbation validation and cross-region consistency, termed Evolutionary Driver Gene identification based on GeneSwitch kinetics (EvoDrGK). We first defined CPM within the 3F (Field–Flow–Front) framework and constructed the evolutionary field map (EFM), thereby representing tumour evolution as a continuous dynamical system embedded in physical space. We then modelled gene switching dynamics along the CPM axis (CPM-GeneSwitch), transforming static differential signals into monotonic switching events along the evolutionary trajectory. Building on this, we incorporated protein–protein interaction (PPI) priors and applied a graph attention network (GAT) to extract candidate driver genes from switch genes that are more likely to govern regulatory propagation.^69, 70^ Cross-region drivers were subsequently aggregated into stable driver modules, and their contributions were projected onto the Front within the 3F framework to quantify their evolutionary impact. Finally, pseudo-spot aggregation and scTenifoldNet((v1.3) in silico knockout were used for network-level counterfactual validation, and functional axis projection together with cosine similarity analysis was performed to assess whether downstream responses of driver modules exhibit reproducible patterns across regions.^71^

### CPM-GeneSwitch: gene switching kinetics along the evolutionary axis

If a gene contributes to clonal evolution, its activation state is expected to exhibit systematic switching along CPM. To characterise transcriptional program switching along the CPM axis under the 3F framework, we performed CPM-aligned gene state switching analysis independently in all regions. Spatial transcriptomics data from each region were integrated with corresponding CPM annotations derived from the Evolutionary Field Map (EFM), establishing a one-to-one correspondence between gene expression matrices and CPM vectors at the spot level. Gene expression states were binarised at the spot level as ON (log-normalized expression > 0) or OFF (otherwise). To avoid instability caused by extremely sparse or ubiquitously expressed genes, only genes with at least 50 ON and 50 OFF spots within each region were retained.

For each gene, we fitted a logistic regression model:

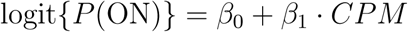

where *β*_1_ captures the systematic change in gene activation probability along the CPM axis. A positive *β*_1_ indicates an OFF→ON activation trend during evolutionary progression, whereas a negative *β*_1_ indicates an ON→OFF repression trend. The switching position (switch point) is defined as the CPM value where the predicted probability of the ON state equals 0.5 (switch cpm = −*β*_0_/*β*_1_).

Statistical significance was assessed using the Wald test for *β*_1_. P values were adjusted within each region using the Benjamini–Hochberg procedure to control the false discovery rate (FDR). We further computed McFadden’s pseudo-R^2^ (1 −deviance/null deviance) to quantify the explanatory power of CPM for gene ON/OFF states. Only genes with switch cpm within the interval [0, 1] were retained for downstream analyses. For each region, a complete CPM-GeneSwitch result table was generated, and significantly switching genes were extracted based on FDR thresholds (< 0.05 and < 0.20) for pathway enrichment and cross-region evolutionary comparison.

### Driver gene inference using PPI-based graph neural networks

Because gene switching alone does not necessarily imply causal driving, we further incorporated protein–protein interaction (PPI) network structure as a prior scaffold for regulatory propagation. For each region, a subgraph was constructed using switching genes as nodes. Node features were derived from CPM-GeneSwitch kinetic parameters, including switch cpm, slope (*β*_1_), pseudo-R^2^, and ON/OFF statistics (for example frac on, *n*_on_, *n*_off_), and were standardized prior to model training.

To ensure cross-region comparability and avoid instability inherent to purely unsupervised ranking, we introduced weak supervision anchors using the inflammatory Hallmark gene set. Genes belonging to this gene set were assigned positive labels, whereas all other genes were treated as unlabeled.

Driver inference was performed using a two-layer graph attention network (GAT) trained on the STRING PPI graph G = (V, E). For each gene g with feature vector **x***_g_*, the model predicts a continuous score:

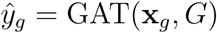

which reflects the likelihood that the gene functions as an evolutionary driver. Predicted scores were subsequently min–max normalized to obtain the final driver score (driver score). Candidate driver genes within each region were then ranked according to this score.

By embedding GeneSwitch-derived kinetic signals into the protein interaction network, this approach enables network-aware inference of evolutionary drivers while incorporating prior biological knowledge from PPI topology.

### Cross-region driver module construction and quantification of “evolutionary force”

To elevate driver genes from the gene level to interpretable dominant forces, we performed cross-region integration and modular modelling of driver gene results. For each gene, we constructed a feature vector **f***_g_* = (dir*_g_*, *CPM*_median_, *CPM*_IQR_, *R*^2^ _mean_, driver_mean_, inflam frac) summarising cross-region occurrence frequency, switching stage characteristics and driver strength.

Clustering in the **f***_g_* feature space was performed using k-means (*k* = 4) to obtain modules *M*_1_–*M*_4_. For each module, we summarised the proportion of OFF→ON genes, CPM positions, inflammatory enrichment, housekeeping gene fraction and driver score levels, and interpreted the modules as immune/stress front modules, metabolic–mitochondrial attenuation modules, epithelial identity shutdown modules and ECM-driven evolutionary front modules.

To quantify the effective driving contribution of functional modules to tumour front expansion, we defined a module-level evolutionary force metric. The front contribution of gene g in region r was defined as

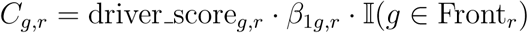

and the evolutionary force of module M in region r was calculated as

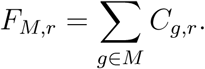

This formulation yields module-specific evolutionary forces across regions, enabling identification of dominant driver modules in front progression and their regional heterogeneity.

### Causal validation via in silico knockout

To provide biologically grounded validation closer to causal perturbation, we performed in silico knockout (KO) analysis for the top-ranked driver genes within each module and region. To mitigate sparsity and technical noise in spatial transcriptomic spots, raw spots were first aggregated into N = 150 pseudo-spots using principal component analysis followed by k-means clustering, and counts were summarised for each pseudo-spot. Regulatory networks were then reconstructed using scTenifoldNet from the wild-type expression matrix (X) and from a perturbed matrix (X^(−^*^g^*^)^) in which gene g was set to zero. The KO-induced network perturbation was defined as the difference between the perturbed and wild-type networks (Δ*W_g_* = W ^(−^*^g^*^)^ − W). For each gene, perturbation strength was quantified as the norm of the differential regulatory vector for that gene. To further characterise KO-induced network rewiring, we visualised strengthened and weakened regulatory connections and quantified perturbation breadth and magnitude based on the number of significantly affected genes and the cumulative regulatory distance. Finally, KO-induced differential regulatory vectors were projected onto predefined functional axes (inflammation, metabolism, proliferation, stress and EMT) to construct region-level response profiles. Cosine similarity across regions within each module was used to quantify response coherence, thereby evaluating the consistency and robustness of dominant functional drivers across spatial regions.

### DepMap dependency analysis of evolutionary modules

To assess whether the four evolutionary driver modules (M1–M4; comprising 83, 17, 21, and 20 genes, respectively) constitute tumour-cell-intrinsic dependencies, we analysed genome-wide CRISPR–Cas9 loss-of-function dependency data from the Cancer Dependency Map (DepMap, release 26Q1 – Portal update), covering 18,531 genes across 1,208 profiled cell lines. Breast cancer cell lines were identified from the model annotation table by OncotreeLineage == "Breast", excluding non-malignant entries (fibroblast and immortalized breast lines); of 92 candidate breast models, 53 lines with available CRISPR dependency profiles were retained for analysis. Gene identifiers in all matrices were stripped of their Entrez suffixes to match module gene symbols. For each module, we quantified two complementary metrics: (i) the proportion of constituent genes annotated as pan-essential in the inferred common-essential list, and (ii) the distribution of per-cell-line gene dependency probabilities across the 53 breast lines, where values >0.5 denote a genuine dependency. To provide a reference baseline, a size-matched random background (≈9.6%) was estimated by randomly sampling equally sized gene sets from all assayed genes over 1,000 iterations and averaging the resulting common-essential proportion. All analyses were performed in R using the data.table package, and results were visualized with ggplot2.

### Spatial cell–cell interaction and directional signalling inference

Following module-level evolutionary driver analysis, we selected a representative cancer microzone (CMZ) from each region together with surrounding cancer-associated fibroblasts (CAF) and lymphocytes (immune cells) to perform spatial cell–cell interaction analysis. Expression matrices were obtained from the log-normalized “data” layer of the Seurat Spatial assay. Cells were classified into Tumor, CAF and Immune groups based on the CCI Label annotation.

For each sender–receiver pair, spatial proximity was quantified using a k-nearest-neighbour approach (*k* = 10). Sender cells were partitioned into “near” and “far” groups according to the minimum spatial distance to receiver cells, using a fixed threshold of 50 µm. When the fixed threshold failed to meet the minimum cell number requirement (≥ 30 cells), an adaptive quantile-based strategy (q = 0.15–0.40) was applied to ensure statistical robustness. Differential expression between near and far sender cells was performed using the Wilcoxon rank-sum test (min.pct = 0.05). Genes upregulated in the near group were ranked by p value and the top 200 genes were selected as the candidate target gene pool.

Ligand–target regulatory activity was inferred using the NicheNet framework (nichenetr v2.2.1), incorporating the pre-trained ligand target matrix and lr network resources.^72^ Candidate ligands were defined as ligands expressed in sender cells and present in the NicheNet ligand database. For each ligand, regulatory activity was quantified as the Pearson correlation between the ligand-specific regulatory weight vector **w***_l_* and the binary target gene vector **y***_T_*, activity(l) = corr(**w***_l_*, **y***_T_*). Only ligands with positive activity scores were retained and ranked.

To estimate ligand–receptor interaction strength, average expression levels of ligands and receptors were calculated in sender and receiver cells, respectively. Interaction strength for each ligand–receptor pair was defined as LR strength(l, r) = mean*_send_*(l) × mean*_recv_*(r). All ligand activity scores and LR interaction strengths were computed independently for each region and sender–receiver direction.

To integrate spatial signalling with the evolutionary framework, active ligands and LR interaction strengths were mapped to the previously defined cross-region driver modules (M1–M4). For each region and interaction direction, average ligand activity and LR strength were calculated for genes within each module to generate module-level directional interaction scores. These scores were visualised using ordered heatmaps (ComplexHeatmap) to characterise module-specific signalling patterns across regions and interaction directions.

### CODEX-based spatial validation of the 3F model

CODEX multiplexed imaging data of breast cancer samples (HTAPP series) were obtained from the Human Tumor Atlas Network (HTAN) Data Portal(https://humantumoratlas.org), generated as part of the study by Iglesia et al.^73^ Single-cell protein-expression matrices and spatial coordinates were extracted directly from .h5ad files using the hdf5r R package, and cell types (tumour, cancer-associated fibroblasts [CAFs], and immune subsets) were assigned by marker-based gating on z-scored protein intensities, using *KRT19* for tumour cells, *FAP* /*VIM* for CAFs, and canonical lineage markers (*PTPRC*, *CD4*, *CD8A*, *FOXP3*, *PDCD1*, *HLA-DRA*, *ITGAX*, *CD274*, *CD19*) for immune populations. CMZs, defined as tumour–CAF interface regions, were manually delineated using an interactive lasso-selection tool built in Shiny and plotly, yielding 23 CMZ ROIs from 10 patients; cells left unassigned within a CMZ were classified as tumour cells based on histological context. For each CMZ, tumour cells were stratified into concentric Inner/Mid/Outer layers along the core-to-periphery axis, defined by the relative Euclidean distance of each cell to the tumour centroid, and a per-cell invasiveness/ECM score (mean z-score of *VIM*, *COL4A2*, *PDPN*, with epithelial *KRT19* inverted) and an inflammatory-stress score (mean z-score of *HIF1A* and *HLA-DRA*) were computed. Spatial gradients of each program were assessed by linear mixed-effects models (program score ∼ relative distance + (1 | patient)) and by paired Wilcoxon tests comparing per-ROI inner versus outer layers, with all analyses performed in R using the hdf5r, data.table, lme4, lmerTest, and ggplot2 packages.

### Characterization of therapy-induced subclonal changes and their association with evolutionary driver modules

Single-cell transcriptomic data were obtained from the publicly available dataset GSE303201 (Gene Expression Omnibus), which contains single-cell RNA sequencing profiles of TNBC patient-derived xenograft (PDX) models before and after chemotherapy treatment (Capecitabine or Anthracyclines), including 10 samples from five models (HBCx39, HBCx95, HBCx172, HBCx221 and HBCx218). The dataset provides paired pre- and post-treatment tumor-derived single-cell transcriptomes from the same PDX models, enabling analysis of therapy-induced subclonal evolution; therefore, all cells were treated as tumor cells without specifying an external normal reference. To infer intratumoral copy number variation (CNV) structure and define subclonal lineages, raw count matrices from each sample were analyzed using inferCNV (v1.26.0). Genes were ordered according to hg38 genomic annotation to construct inferCNV objects, and CNV smoothing was performed without specifying reference normal populations (cutoff = 0.1, HMM = FALSE, denoise = FALSE). The resulting CNV expression profiles were used for subclonal stratification. Unsupervised clustering was performed on CNV profiles to identify subclonal populations. Pairwise Euclidean distances between cells were computed, followed by K-means clustering with candidate cluster numbers ranging from k = 2 to 6 (or k = 4 to 6). The optimal number of clusters was selected based on the mean silhouette coefficient. Each cell was assigned a CNV subclone label (for example, CNV C1, CNV C2) for downstream analysis (Supplementary Table 3). To characterize transcriptional features of each subclone, standard single-cell preprocessing was performed for each sample, including normalization, variable feature selection, scaling and principal component analysis. Differential expression analysis was conducted using subclone labels as grouping variables (Seurat FindAllMarkers, min.pct = 0.25, logfc.threshold = 0.25) to identify subclone-specific marker genes. To quantify the activity of evolutionary driver programs at the subclonal level, signed module scores were computed based on previously defined cross-region driver gene modules (M1–M4). For each module, genes were assigned directional weights according to CPM-GeneSwitch kinetics (OFF-to-ON assigned +1, ON-to-OFF assigned −1), and weighted mean expression across module genes was calculated for each cell to obtain module activity scores. Module scores were subsequently averaged within each subclone to quantify the extent to which different subclones carried evolutionary programs related to ECM/front activity, inflammatory stress, metabolic attenuation and epithelial identity suppression. Comparisons of subclonal composition, differential expression signatures and module activity distributions before and after treatment were used to assess therapy-induced subclonal reorganization and its association with evolutionary driver modules.

### SingleR reference-based cell type annotation (GSE161529)

In the breast cancer single-cell RNA-seq dataset GSE161529, we applied a reference-based cell type annotation strategy. Briefly, the merged Seurat object was loaded and multi-layer structures were integrated to ensure consistency across expression layers. The RNA assay was then normalized, and highly variable genes (HVGs, n = 2000) were identified. The object was subsequently converted into a SingleCellExperiment(v1.32.0) to facilitate downstream annotation. Cell type inference was performed in parallel using three independent reference datasets: BlueprintEncode, Monaco Immune, and HumanPrimaryCellAtlas, with their respective main label systems used as output annotations. The prediction was conducted in the HVG-restricted log-normalized expression space, with internal marker gene identification based on a Wilcoxon framework. A pruning strategy was enabled to reduce the impact of uncertain or conflicting annotations. The inferred labels were written into the cell metadata and further consolidated into 11 major cell types (celltype major) based on a predefined mapping table, with unmatched labels assigned to “Other”. We exported per-sample annotation summaries at both the raw label level and the 11-class aggregated level, and saved the final annotated object.^74^

### inferCNV for malignant epithelial cell identification and CNV subclone inference

To identify malignant epithelial cells at the sample level while controlling for sample-specific variability in CNV inference, inferCNV analysis was performed independently for each sample. The reference population was defined as global T-cells, with a minimum of 200 reference cells required to ensure stable background estimation. For each sample s, a subset was constructed containing epithelial cells from that sample together with global T-cells, and the raw count matrix was used as input for inferCNV. A cell annotation file and a genomic gene order file were provided to enable CNV inference based on chromosomal positioning. Key inferCNV parameters included cutoff=1, denoise=TRUE, cluster by groups=TRUE, and HMM=TRUE to obtain discrete CNV state predictions at the regional level. For each sample, the HMM output table was parsed, and cells exhibiting any non-neutral CNV state were defined as candidate malignant cells. The intersection of this set with epithelial cells yielded the final malignant epithelial population, and malignancy status was recorded in the cell metadata (e.g., Epithelial malignant and Epithelial normal). For downstream CNV subclone analysis, the observation groupings output from inferCNV was further processed to map cells to their corresponding subclone indices. These indices were reformatted into standardized labels (e.g., Epithelial s1, Epithelial s2) and stored in the metadata for subsequent stratified analyses.

### Construction and Prediction of Evolutionary advantage subclone load

In this study, we propose E-load (Evolutionary Advantage Load), a quantitative metric designed to characterize the proportion of tumor cells exhibiting “advantage-like” transcriptional features and their associated evolutionary burden, and to evaluate its inferability and prognostic relevance in larger patient cohorts. Methodologically, evolutionary advantage subclones defined from spatial transcriptomics were used as weakly supervised ground truth. Through differential expression analysis between **advantage** and **non-advantage** cells, we constructed an Advantage core gene set. A continuous AdvantageScore was then derived in single-cell transcriptomic data, and advantage-like tumor cells were identified using a spatially calibrated threshold. At the sample level, E-load was subsequently defined and further projected across scales to bulk RNA-seq cohorts for clinical and prognostic analyses.

### Construction of an evolutionary advantage–associated gene pool

Based on the definition of evolutionary advantage subclones within the 3F framework, we first generated binary advantage annotations at the cellular level: advantage = 1 indicates cells in an evolutionarily advantageous state, whereas advantage = 0 indicates non-advantage states. These cell-level labels served as weak supervision for downstream gene selection and model calibration. Subsequently, within each region, we compared transcriptional differences between advantage and non-advantage cells, and identified genes significantly upregulated in advantage cells (adjusted P < 0.05, avg log_2_FC > 0). To obtain robust advantage-associated molecular features across regions, we integrated significantly upregulated genes from all regions to construct a unified advantage gene pool. For each gene in the pool, we quantified its occurrence across spatial regions and corresponding expression effects, including the number of regions in which it was identified as advantage-upregulated, as well as its cross-region mean and median log2 fold change. To avoid the loss of potentially relevant signals due to overly stringent filtering at early stages, no hard filtering was applied to the gene pool; instead, all candidate genes were retained and subsequently ranked.

### Determination of the Advantage core gene set, construction of Advantage Score, and threshold learning

Based on the ranked advantage gene pool, we iteratively selected top-ranked gene subsets of varying sizes to evaluate their discriminative ability in distinguishing advantage from non-advantage subclones. For each candidate gene set, we computed the average expression of the gene set in tumor cells from spatial transcriptomics data and defined it as the Advantage Score, a continuous metric reflecting the advantage-associated transcriptional state of individual tumor cells. Using cell-level advantage annotations as ground truth, we performed ROC analysis and applied the Youden index to determine the optimal classification threshold, thereby systematically evaluating the performance of gene sets of different sizes in distinguishing advantage and non-advantage states. Balancing discriminative performance and cross-region robustness, we ultimately selected the top 80 ranked genes as the Advantage core gene set (Top80). Accordingly, the Advantage Score was defined as the mean expression of the Advantage core gene set (Top80) for each tumor cell. Based on thresholds learned from spatial data, we averaged the optimal thresholds across all regions to derive a unified decision criterion. The final global threshold was 0.523833, which was used to classify tumor cells into advantage-like and non-advantage states. This threshold was fixed in subsequent analyses to enable cross-modality projection and comparison of evolutionary advantage states.

### Calculation of E-load in single-cell RNA-seq data

We analyzed the breast cancer single-cell transcriptomic dataset GSE161529. After loading the Seurat object, we performed reference-based cell type annotation using SingleR: following normalization and highly variable gene selection (HVG, n=2000), the data were converted to a SingleCellExperiment object. Subsequently, cell type inference was conducted in parallel using three reference datasets—BlueprintEncode, Monaco Immune, and HumanPrimaryCellAtlas (label.main)—on the HVG-restricted log-normalized expression matrix (prune=TRUE). Predicted labels were then consolidated into 11 major cell types (celltype major). To identify malignant epithelial cells, we performed inferCNV analysis within each sample, using global T-cells as the reference population to infer CNV profiles for epithelial cells. Cells annotated as *Cancer Epithelial* were selected as tumor epithelial cells for downstream analysis. At the single-cell level, the AdvantageScore for each tumor cell was computed using the Advantage core gene set (Top80) derived from spatial data, and a unified threshold was applied to classify cells into advantage-like and non-advantage states. The proportion of advantage-like cells within each sample was then calculated and defined as E-load (Evolutionary Advantage Load), representing the overall burden of evolutionarily advantageous tumor cells at the sample level (Supplementary Table 4).

### Construction of a four-gene E-load prediction model and projection to bulk cohorts

To generalize E-load from single-cell spatial data to conventional bulk transcriptomic cohorts, we constructed an E-load prediction model based on 38 breast cancer samples from GSE161529 with annotated ER/PR/HER2/Ki-67 status. For each sample, the mean expression of ESR1, PGR, ERBB2 and MKI67 was extracted and min–max normalized to obtain four features: ER ESR1, PR PGR, HER2 ERBB2 and KI67 MKI67. The sample-level E-load (proportion of advantage-like cells) was used as the continuous response variable and combined with the four-gene expression features to form the training dataset. We fitted four types of regression models (Supplementary Table 5): (1) linear regression (Eload ~ linear terms of the four genes); (2) polynomial regression (including quadratic terms); (3) random forest regression; (4) quasi-binomial regression with a logit link, used to model “beta-like” E-load constrained within the [0,1] interval (Eload beta, with small-value truncation to avoid exact 0 or 1). For the quasi-binomial model, the continuous proportion response was first truncated to ensure numerical stability:

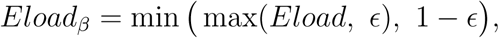

where ɛ = 10^−4^. A generalized linear model with a logit link was then fitted:

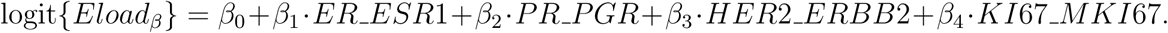

Model parameters were estimated using quasi-likelihood, allowing the variance of the response to deviate from the standard binomial assumption and thereby better accommodating continuous proportion data. Predicted values were obtained through inverse-logit transformation:

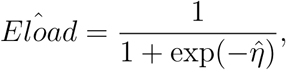

where η̂ denotes the linear predictor. This modeling strategy preserves probabilistic interpretability while improving stability and generalization for outcomes bounded between 0 and 1. Model performance was evaluated using the correlation between predicted and observed E-load and the root mean square error (RMSE), and predictions were constrained within the [0,1] interval. Given its suitability for proportional outcomes and favorable performance, the quasi-binomial (“beta-like”) model was selected as the primary model for downstream bulk analysis. The same min–max normalized four-gene expression features were then extracted in two bulk cohorts: the TCGA-BRCA first-line chemotherapy cohort and the I-SPY1 (GSE32603) baseline tumor microarray dataset (with microarray signals adapted to ER/PR/HER2/Ki-67 expression proxies and normalized to 0–1). The trained beta model was applied to these cohorts to obtain sample-level predicted E-load (Predicted E-load, range 0–1), which was used for downstream recurrence and survival analyses. For visualization and stratification, Predicted E-load values were further partitioned using K-means clustering (K=3), and their distributions were displayed using boxplots overlaid with individual data points.

### Cohort selection and patient filtering

TCGA-BRCA data were obtained from the GDC Data Portal, including clinical annotations and Illumina RNA-seq expression matrices. We included patients who received first-line chemotherapy and had valid RNA-seq data, and further excluded samples lacking follow-up time, age, or molecular subtype information, yielding the final TCGA cohort for overall survival (OS) analyses (Supplementary Table 6). The I-SPY1 dataset was retrieved from GEO (GSE32603), profiled on the GPL14668 microarray platform using baseline tumor specimens. GEO-provided GSM identifiers were matched to patient IDs in the I-SPY1 clinical database, and age, HR/HER2 status, pathologic complete response (pCR), and long-term follow-up information were integrated. Only patients with clearly defined OS or recurrence-free survival (RFS) endpoints were retained (Supplementary Table 7). Because RFS captures early treatment-related failure whereas OS reflects longer-term disease burden and event accumulation, training cohorts were selected according to endpoint-specific characteristics. For RFS analyses, I-SPY1—featuring standardized treatment and high-quality recurrence annotation—was used as the training cohort and TCGA-BRCA as the validation cohort. For OS analyses, TCGA-BRCA—providing longer follow-up and more OS events—served as the training cohort and I-SPY1 as the validation cohort. Bidirectional cross-cohort validation was performed to assess directional consistency and generalizability across cohorts.

### Modeling and validation of recurrence-free survival (RFS)

#### Training cohort: I-SPY1 (GSE32603)

In the I-SPY1 cohort, recurrence-free survival time (RFS time) and recurrence/progression event indicators (RFS event) were extracted from the curated clinical table and merged with beta model–predicted E-load based on GSM sample IDs, retaining only patients with non-missing RFS time, RFS event, and Eload. We first examined the crude association between Predicted E-load and recurrence events in a two-dimensional manner: Wilcoxon rank-sum tests were used to compare E-load distributions between the no-event group and the recurrence/death group, and Spearman correlation analysis was performed to assess the monotonic relationship between E-load and RFS time, with LOESS-smoothed scatter plots for visualization. Subsequently, using the median Predicted E-load within the cohort as the cutoff, patients were stratified into High E-load and Low E-load groups. Kaplan–Meier analysis was applied to estimate RFS curves for each group, and cumulative recurrence probabilities (1 − S(t)) at 2, 3, and 5 years were calculated along with their 95% confidence intervals, yielding E-load–stratified bar plots of 2/3/5-year cumulative recurrence risk.^75^ To characterize the nonlinear relationship between continuous E-load and recurrence risk, we fitted a restricted cubic spline Cox model in the I-SPY1 cohort using the rms(v8.1-0) package: Surv(RFS time, RFS event) ~ rcs(E-load, 4 knots). The model returns hazard ratios (HRs) and 95% confidence intervals across different E-load levels, enabling visualization of a smooth E-load–risk curve. Furthermore, using the fitted spline Cox model, we predicted individualized RFS curves S(t—E-load) over 0–5 years at given E-load values (e.g., 0.3 and 0.6), to illustrate dynamic differences in recurrence risk across E-load levels.^76^

#### Validation cohort: TCGA-BRCA first-line chemotherapy subset

As an external validation of the RFS association, we constructed a first-line chemotherapy cohort within TCGA-BRCA by integrating clinical and follow-up data, selecting patients who completed first-line treatment, received chemotherapy, had RNA-seq data available, and possessed complete OS/DFS information. Recurrence/progression status was defined based on progression or recurrence and related time fields from follow-up records, combined with death information to derive DFS/RFS events and time. For recurrence analysis, Predicted E-load (beta model) was merged with patient id, retaining only samples with complete RFS/DFS time, event, and E-load. The same analytical workflow as in I-SPY1 was applied: (1) comparison of E-load between no-event and recurrence/death groups (Wilcoxon test); (2) assessment of the Spearman correlation between E-load and RFS time; (3) stratification into High/Low groups using the cohort median E-load and estimation of 2/3/5-year cumulative recurrence probabilities with confidence intervals; (4) fitting a restricted cubic spline Cox model with continuous E-load and plotting HR curves. These analyses were used to evaluate the transferability of the E-load–recurrence association and potential signal attenuation in a heterogeneous real-world cohort.

### Construction, external validation, and internal calibration of the overall survival (OS) model

#### Training cohort: TCGA-BRCA first-line chemotherapy cohort

In the TCGA-BRCA first-line chemotherapy + RNA-seq cohort, we merged beta model–predicted E-load with OS data (OS time, Event OS), and further incorporated age (Age) and molecular subtype (Subtype bin, Basal vs Non basal) as covariates. Using the median Predicted E-load of the entire cohort as a fixed cutoff, patients were stratified into Low E-load and High E-load groups (Eload grp), with the cutoff held constant in all TCGA analyses. Kaplan–Meier analysis was performed to estimate OS curves for each group, and log-rank tests were used to compare survival differences between High and Low groups. Risk tables over time were also generated to provide a descriptive overview of the relationship between E-load and survival outcomes, forming the basis for subsequent Cox modeling. On this basis, we constructed the following Cox models: (1) univariable models with Age, Subtype bin, and Eload grp as predictors, respectively; (2) multivariable clinical model: Surv(OS time, Event OS) ~ Age + Subtype bin; (3) multivariable extended model: Surv(OS time, Event OS) ~ Age + Subtype bin + Eload grp. For each model, regression coefficients, hazard ratios (HRs), 95% confidence intervals, and p values were recorded. Likelihood ratio tests were used to compare models with and without Eload grp to assess whether inclusion of E-load significantly improved model fit. Harrell C-index and AIC were calculated to quantify discrimination and overall model performance.^77, 78^ The proportional hazards assumption was evaluated using cox.zph, and multicollinearity among covariates was assessed using variance inflation factors (VIF).^79^ To compare time-dependent predictive performance between the clinical model and the clinical + E-load model, timeROC(v0.4) was applied to compute time-dependent AUCs at 3, 4, and 5 years, based on linear predictors from the Clinical and Clinical + Eload grp models, respectively, and AUC–time curves were plotted.^80^ Decision curve analysis (DCA) was further performed to construct net benefit–threshold probability curves at 5-year OS, comparing the Clinical and Clinical + Eload grp models against “treat all” and “treat none” strategies, thereby evaluating the potential clinical utility of incorporating E-load.^81^ To mitigate overfitting and obtain robust performance estimates, we fitted a CPH model using the rms package: Surv(OS time, Event OS) ~ Age + Subtype bin + Eload grp, and performed internal validation via bootstrap resampling (B = 200).^82^ The validate function was used to obtain Dxy and its optimism-corrected values, which were further converted to optimism-corrected C-index. Additionally, bootstrap-based calibrate was used to generate calibration curves at 5 years, comparing predicted and observed OS probabilities to assess calibration accuracy.

#### External validation cohort: I-SPY1 (GSE32603)

In the I-SPY1 cohort, beta model–predicted E-load from microarray expression data was merged with OS time, OS event, and clinical covariates (Age, HR HER2 STATUS). HR HER2 STATUS was further binarized into TripleNeg and non TripleNeg (Subtype bin), and patients were stratified into High/Low E-load groups (Eload grp) using the cohort-specific median cutoff. The same KM and Cox analysis workflow as in TCGA was then applied: (1) Kaplan–Meier OS curves stratified by Eload grp with log-rank testing; (2) multivariable Cox modeling Surv(OS time, OS event) ~ Age + Subtype bin + Eload grp, comparing HRs and confidence intervals with those from the TCGA training cohort; (3) computation of time-dependent AUCs for Clinical and Clinical + E-load models over 3–5 years and plotting AUC–time curves; (4) decision curve analysis at 5-year OS. These analyses were used to assess the generalizability of the E-load–based prognostic model trained in TCGA to an independent clinical trial cohort.

### Survival analysis

Survival curves were estimated using the Kaplan–Meier method and compared between groups using the two-sided log-rank test.^75^ Hazard ratios were estimated using the Cox proportional hazards model.^76^ Statistical inference for covariates was based on Wald tests and 95% confidence intervals. Model fitting and inference were performed using the survival and rms packages in R.

### Model performance evaluation

Model discrimination was evaluated using Harrell’s concordance index (C-index).^77^ Time-dependent receiver operating characteristic analysis was performed using the timeROC package to estimate AUC values at prespecified time points (3, 4 and 5 years).^80^ Model calibration and clinical usefulness were further assessed using decision curve analysis.^81^ Model complexity and goodness-of-fit were compared using the Akaike information criterion (AIC).^78^ Multicollinearity among covariates was evaluated using variance inflation factors (VIF).^79^

### Internal validation

Internal validation was performed using bootstrap resampling with B = 1000 iterations.^82^ In each bootstrap sample, the model was refitted and evaluated on both the bootstrap sample and the original dataset to estimate optimism in model performance. Optimism-corrected discrimination metrics were obtained using the validate function from the rms package, which reports Somers’ *D_xy_* and the corresponding C-index after optimism correction.

### Computational methods and software versions

All computational analyses were performed in R (version 4.5.1) and Python (version 3.13.9). Key R packages included Seurat (v5.4.0), Monocle2 (v2.24.1), spacexr/RCTD (v2.2.0), SingleR (v2.12.0), infercnv (v1.26.0), clusterProfiler (v4.18.4), scTenifoldNet (v1.3), survival (v3.8.3), survminer (v0.5.1), rms (v8.1-0), timeROC (v0.4), randomForest (v4.7-1.2), lme4 (v2.0.1), lmerTest (v3.2.1), mgcv (v1.9.3), hdf5r (v1.3.12), shiny (v1.11.1), plotly (R; v4.11.0), dplyr (v1.1.4), tidyr (v1.3.1), data.table (v1.17.8), and ggplot2 (v4.0.1). Key Python libraries included opencv-python (v4.13.0), numpy (v2.3.5), tifffile (v2026.1.14), nibabel (v5.3.3), networkx (v3.1), matplotlib (v3.10.8), plotly (v6.5.2), scipy (v1.16.3), and scikit-image (v0.26.0). Spatial and imaging data were processed and visualized using Space Ranger (v3.0.1), Loupe Browser (v8.0.0), Xenium Explorer (v4.1.1), Mantis Viewer (v1.2.4-alpha.3), QuPath, CaseViewer (3DHISTECH), ImageViewer (SQray), and the 3DHISTECH Slide Converter; protein–protein interaction networks were retrieved from STRING and visualized in Cytoscape. CRISPR gene-dependency validation used the DepMap dataset (Broad Institute). The in-house frameworks developed in this study (CMZ reconstruction, CPM, 3F, EFM, EvoDrGK, and E-load) were implemented in R and Python.

### Quantification and statistical analysis

All statistical analyses were performed in R (version 4.5.1) unless otherwise specified. No statistical methods were used to predetermine sample size. All analyses were conducted on available samples that passed predefined quality control criteria. Unless otherwise indicated, statistical tests were two-sided.

#### Differential expression analysis

Differential expression analysis was performed using Seurat (v5.1.0 or v5.4.0) with the FindMarkers or FindAllMarkers functions, based on the non-parametric Wilcoxon rank-sum test. P values were adjusted for multiple comparisons using the Benjamini–Hochberg method to control the false discovery rate (FDR). Genes were considered statistically significant at adjusted P < 0.05 unless otherwise specified.

#### Gene set scoring and functional analysis

Gene set activity scores (e.g., EMT, stemness, and other functional programs) were calculated using the AddModuleScore function in Seurat. Differences in functional scores between groups were assessed using non-parametric tests where appropriate. Functional enrichment analyses (GO and KEGG) were performed using clusterProfiler (v4.18.4), with multiple testing correction using the Benjamini–Hochberg method. Enrichment results were considered significant at adjusted P < 0.05.

#### Correlation analysis

Associations between continuous variables (e.g., CPM, pseudotime, functional scores) were evaluated using Spearman rank correlation. Correlation coefficients ( *ρ*) and corresponding P values were reported.

#### Regression and modeling

Logistic regression models were used for binary outcome analyses, and linear regression models were applied for continuous outcomes. For survival analyses, Cox proportional hazards regression models were used to estimate hazard ratios (HRs) and 95% confidence intervals. The proportional hazards assumption was assessed using Schoenfeld residuals. Restricted cubic spline Cox models were fitted using the rms package to evaluate nonlinear associations between continuous variables (e.g., E-load) and survival outcomes.

#### Survival analysis

Survival curves were estimated using the Kaplan–Meier method and compared using the two-sided log-rank test. Time-to-event endpoints included overall survival (OS) and recurrence-free survival (RFS). Time-dependent receiver operating characteristic (ROC) analysis was performed using the timeROC package to estimate area under the curve (AUC) values at predefined time points.

#### Model evaluation and validation

Model performance was evaluated using Harrell’s concordance index (C-index), root mean square error (RMSE), and Akaike information criterion (AIC). Decision curve analysis (DCA) was used to assess clinical utility. Internal validation was performed using bootstrap resampling (typically B = 200 or B = 1000), and optimism-corrected performance metrics were reported.

#### Multiple testing correction

Unless otherwise specified, multiple hypothesis testing was controlled using the Benjamini–Hochberg procedure. Adjusted P values (FDR) were used to determine statistical significance.

#### Data filtering and exclusion criteria

Data points (e.g., spatial bins, cells, or samples) were excluded only based on predefined quality control criteria, including low gene counts, low UMI counts, insufficient cell numbers for analysis, or non-finite values (e.g., NA or Inf). No additional outlier removal was performed unless explicitly stated.

#### Reproducibility

All analyses were performed using publicly available software packages with specified versions. Default parameters were used unless otherwise indicated in the Method details section.

## Data availability

Publicly available datasets analysed in this study include the breast cancer single-cell RNA sequencing datasets GSE176078, GSE303201 and GSE161529 obtained from the Gene Expression Omnibus (GEO); the TCGA-BRCA cohort obtained from The Cancer Genome Atlas (TCGA) via the Genomic Data Commons (GDC) Data Portal; and the I-SPY1 cohort (GSE32603) obtained from GEO. Additional publicly available spatial and imaging datasets were collected as follows: 29 Visium HD and 16 Xenium breast cancer samples from the 10x Genomics dataset repository (https://www.10xgenomics.com/datasets); 32 multiplexed ion beam imaging (MIBI) breast cancer samples from the Angelo Lab (https://www.angelolab.com/mibi-data); 10 breast cancer CODEX datasets from the Human Tumor Atlas Network (HTAN) Data Coordinating Center Data Portal at the National Cancer Institute (https://data.humantumoratlas.org/, under the HTAN WUSTL Atlas); and CRISPR gene-dependency data for 53 breast cancer cell lines from the DepMap portal (https://depmap.org/portal/data_page/?tab=allData). The GRCh38 reference files used for Visium HD spatial transcriptomics analysis are freely available from the 10x Genomics website (https://support.10xgenomics.com). The MSigDB Hallmark gene sets are available from the GSEA database (https://www.gsea-msigdb.org). Protein–protein interaction (PPI) data were obtained from the STRING database (https://string-db.org). The spatial transcriptomic data generated in this study (five Visium HD breast cancer samples) have been deposited in the Genome Sequence Archive for Human (GSA-Human) at the National Genomics Data Center (NGDC), China National Center for Bioinformation (CNCB), under accession number HRA017537 (project accession PRJCA060688), and are publicly available. Any additional information required to reanalyse the data reported in this paper is available from the corresponding authors upon reasonable request.

## Code availability

Custom scripts used for data processing and analysis in this study were implemented in R (v4.5.1) and Python (v3.13.9). The core frameworks for mapping tumour evolutionary dynamics — the Cancer Progression Metric (CPM) and the Field–Flow–Front (3F) model — together with example data and documentation, are publicly available at https://github.com/Euphonni/spatiotemporal-tumor-evolution under the MIT License.

## Acknowledgements

We thank the patients and clinical staff at Sichuan Cancer Hospital for providing tumour samples and clinical information. We thank OE Biotech Co., Ltd. (Shanghai, China) for spatial transcriptomics services, and Hongyan Liu, Lanying Wang and Sheng Pan for technical assistance and bioinformatics support. This work was supported by the Clinical Scientist Program of Sichuan Cancer Hospital (YB2022003), the Sichuan Natural Science Foundation (2022NSFSC0051), the Chengdu Technology Innovation R&D Project (2021YF0501659SN) and the National Natural Science Foundation of China (11375124) to W.W.; the Chongqing Natural Science Foundation (CSTC2021jcyj-msxm3803) to R.L.; the National Natural Science Foundation of China Young Scientists Fund (81402784) to S.L.; the Outstanding Young Investigator Program of Sichuan Cancer Hospital (YB2024016) to J.J.; and the Sichuan Provincial Natural Science Foundation (25NSFSC0425) to Y.L.

## Author contributions

W.W., R.L. and Y.L. conceived and designed the study. Y.Z., J.J., Y.Z., H.Y., J.W. and H.J. collected tumour samples and performed experiments. Y.Z., J.J., Y.P., Y.Y., L.L., S.L., Y.T., L.L., M.G., Z.W. and C.W. performed computational analyses. W.W., R.L., Y.L., Y.Z., J.J. and S.L. interpreted the results. W.W. and Y.Z. wrote the manuscript. W.W., R.L. J.L and Y.L. revised the manuscript. W.W., R.L., Y.L., J.J. J.L and S.L. supervised the study and acquired funding. All authors reviewed and approved the final manuscript.

## Competing interests

The authors declare no competing interests.

---

**Extended Data Fig. 1.**
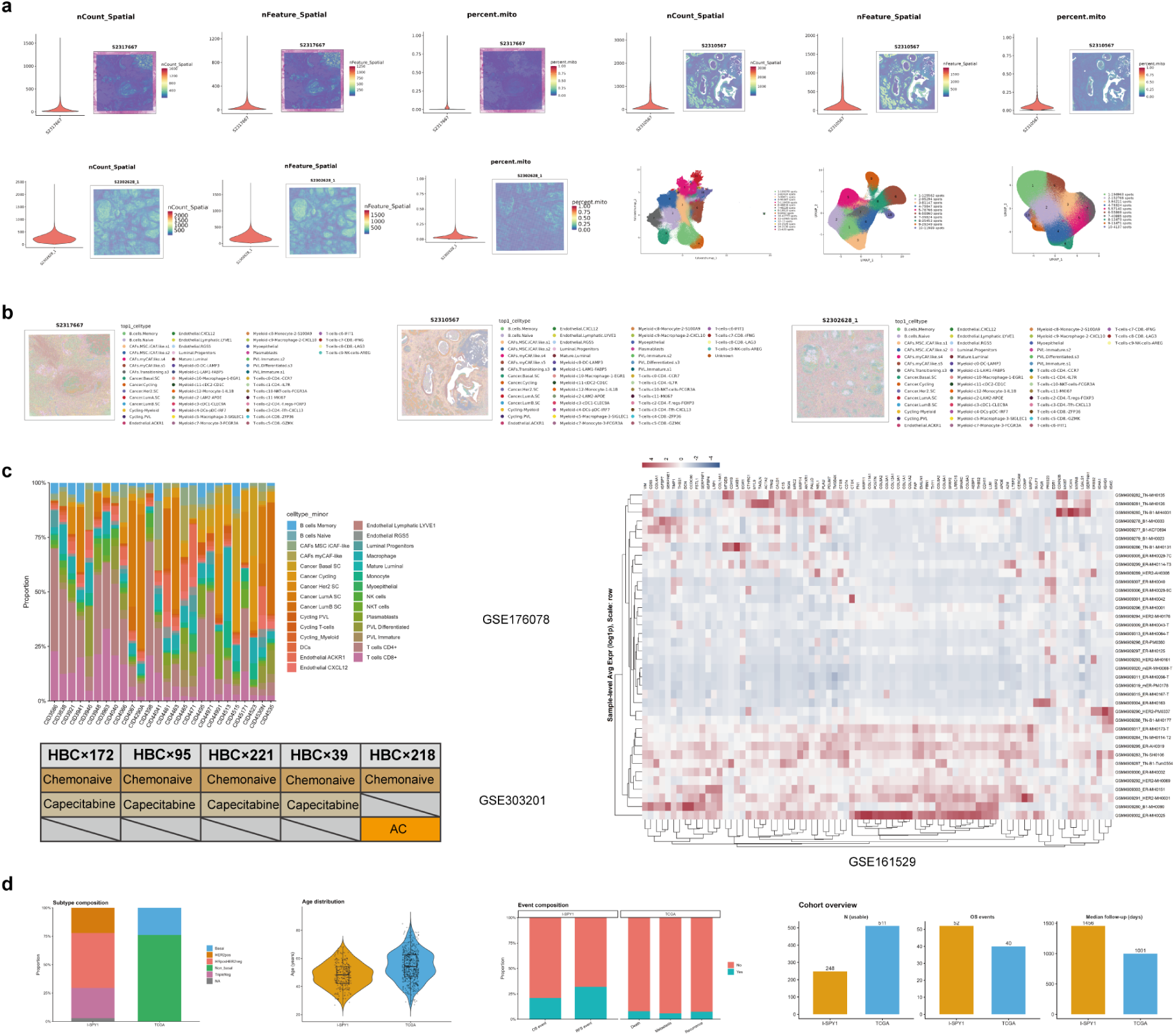
Quality control and overview of spatial, single-cell and bulk transcriptomic datasets. **a,** Quality control and unsupervised clustering of Visium HD spatial transcriptomic data. Data quality was assessed using distributions and spatial feature maps of UMI counts, numbers of detected genes, and mitochondrial transcript proportions. UMAP-based dimensionality reduction and clustering were performed at the spot level to visualize transcriptional heterogeneity and its spatial organization within tissues. **b,** Cell-type deconvolution of spatial transcriptomic data using RCTD. Reference scRNA-seq data were used to infer the dominant cell type at each spatial spot, and spatial maps were generated to display the distribution of malignant, stromal, and immune cells across tissue sections. **c,** Visualization of three published breast cancer scRNA-seq datasets. GSE176078 shows cell-type composition across 26 breast cancer patients; GSE303201 presents sample grouping of patient-derived xenograft (PDX) models derived from five patients; and GSE161529 displays gene expression profiles from 38 breast cancer patients, visualized using scaled average expression heatmaps. **d,** Transcriptomic overview of two breast cancer bulk RNA-seq cohorts. Summaries include molecular subtype composition, age distribution, clinical event profiles, sample availability, numbers of outcome events, and median follow-up time, providing population-level context for spatial and single-cell analyses.

**Extended Data Fig. 2.**
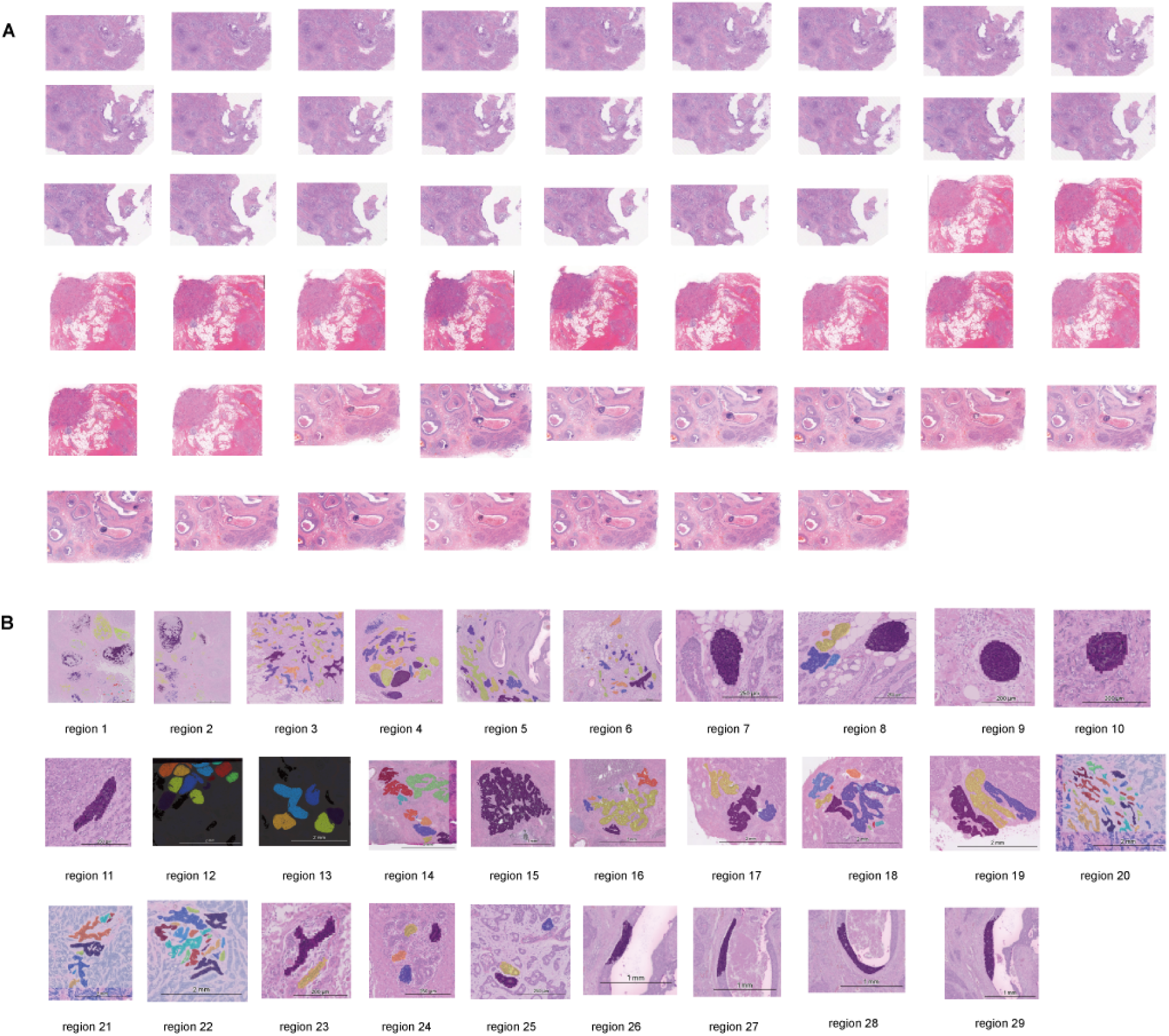
Consecutive histological sections and CMZ/BMZ distribution across regions. **a,** Consecutive whole-slide histological sections from three representative breast cancer samples (sections 1–25 for sample 628; sections 26–38 for sample 667; and sections 30–52 for sample 567). Serial sections illustrate the spatial continuity of tumour architecture across adjacent slices. **b,** Regional selection and spatial distribution of microzones across 29 regions (regions 1–29). Regions 23, 24, 25, 28 and 29 represent normal breast tissue (benign microzones, BMZs), whereas the remaining regions represent tumour tissue (cancer microzones, CMZs). Microzone annotations are overlaid on histological images.

**Extended Data Fig. 3.**
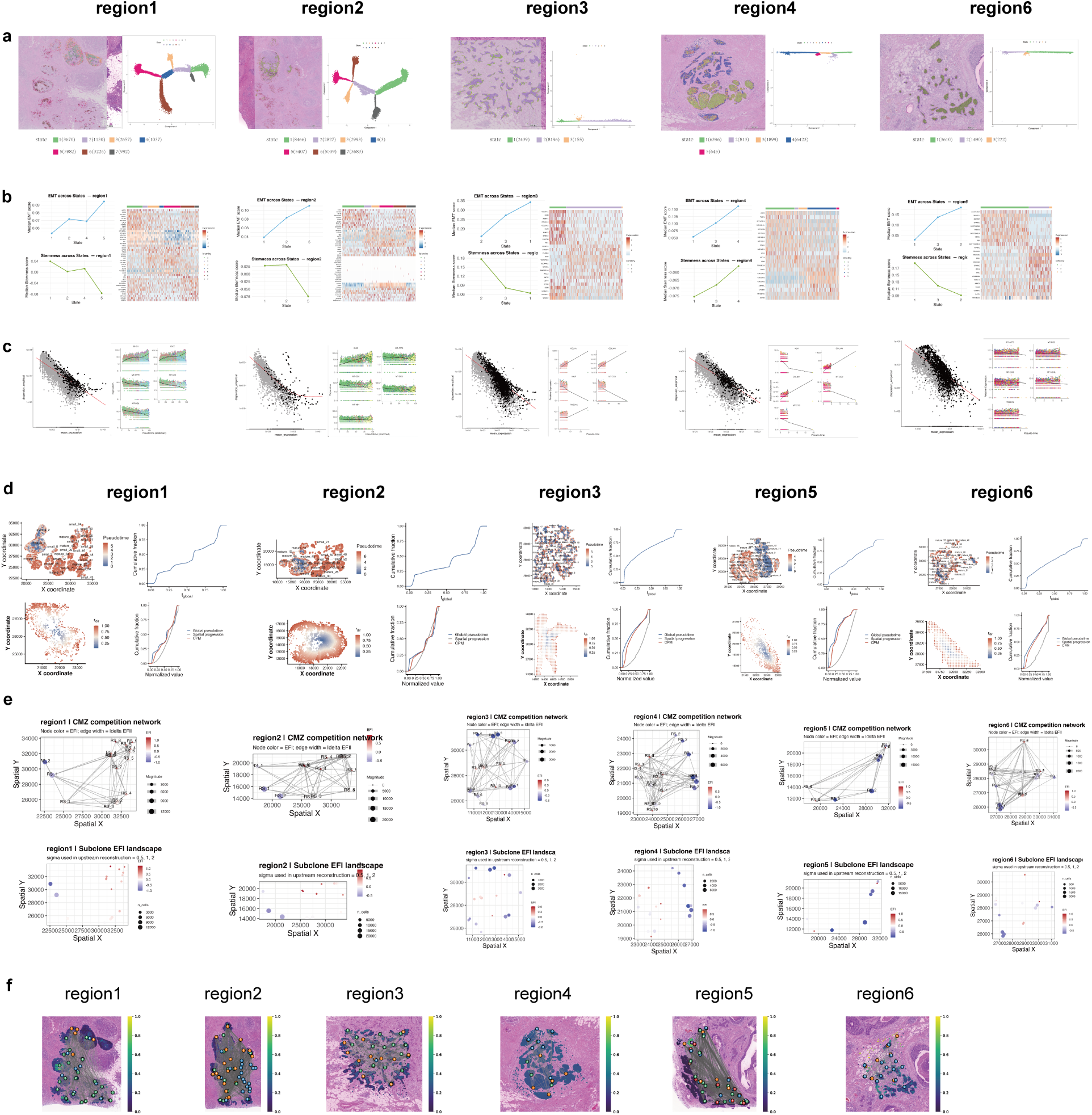
Cross-regional consistency of root state inference and the Cancer Progression Metric. **a,** Monocle2 pseudotime trajectories and spatial mapping of states across regions 1–4 and region6. For each region, tumour cells are projected onto the Monocle2 trajectory and mapped back to their histological locations. Distinct states exhibit coherent spatial organization within CMZs, and the number of tumour cells in each state is indicated, highlighting region-specific differences in state composition and scale. **b,** Validation of root state ordering across regions. States are ordered according to the predefined root-to-late sequence, and median EMT and stemness scores are shown as line plots for each region. Heatmaps display differentially expressed genes across states, illustrating progressive transitions in transcriptional programs along the inferred pseudotime direction. **c,** Quality control of Monocle2 pseudotime analysis across regions. Dispersion-based selection of ordering genes is shown for each region, demonstrating the relationship between mean expression and dispersion used to identify highly variable genes. These distributions indicate that pseudotime reconstruction is driven by robust and informative transcriptional variation across regions. **d,** Construction and cross-regional consistency of CPM. For each region (region1, region2, region3, region5 and region6), spatial maps of global pseudotime (top left) and directionally normalized spatial progression (*r*_dir_, bottom left) are shown together with their empirical cumulative distribution functions (top right). The lower-right panels compare ECDFs of global pseudotime, spatial progression and their weighted integration into CPM. CPM consistently displays a distributional profile distinct from either temporal or spatial variables alone, indicating that it represents an independent continuous evolutionary coordinate reproducibly recovered across spatial regions. **e,** Spatial competition networks and EFI landscapes across tumour regions. Upper panels show CMZ competition networks in each region (node colour, EFI; node size, subclone size; edge width, absolute EFI difference). Lower panels show the corresponding spatial EFI landscapes of CMZ-level subclones. Across regions, higher-EFI subclones form localized^93^clusters and competitive interfaces with neighbouring CMZs. **f,** Spatial trajectory inference generated by STlearn, depicting discrete cellular clusters (nodes) and their connectivity overlaid on the H&E-stained tissue. The colour gradient represents the diffusion pseudotime from the inferred root (dark purple) to the terminal states (yellow).

**Extended Data Fig. 4.**
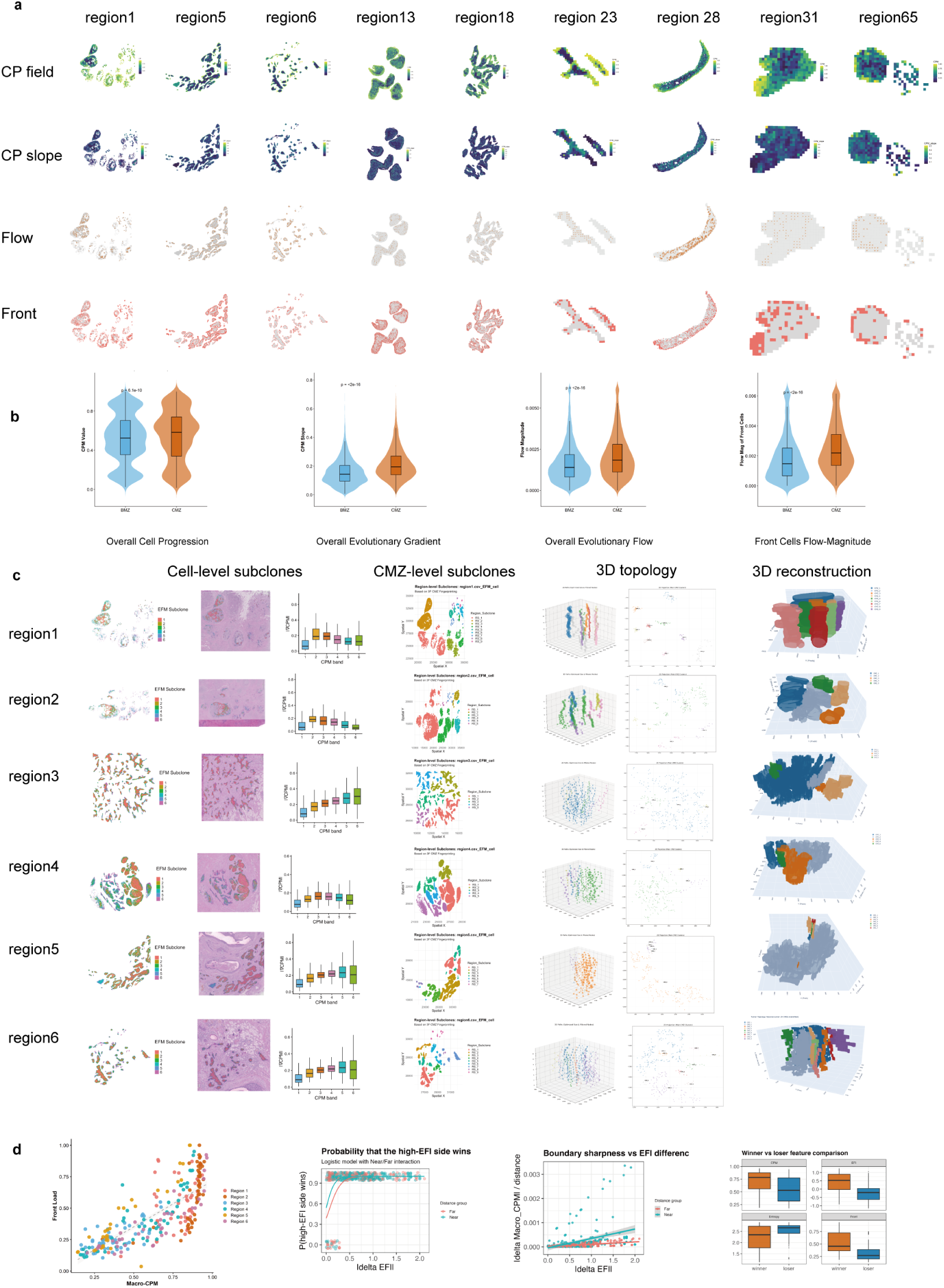
Spatial heterogeneity of 3F dynamics and subclonal architecture across regions. **a,** Spatial distribution of CP field, CP slope, flow and evolutionary front across different tumour regions and normal breast tissue. Shown are the spatial maps of four key 3F model components—CP field, CP slope, flow and evolutionary front—across 8 representative tumour regions (regions 1, 2, 3, 5, 6, 13, 18) and normal breast tissue (regions 23 and 28). Each row corresponds to one of these metrics, visualizing their distribution across these regions. These maps highlight the heterogeneity in tumour progression dynamics and illustrate the differences between tumour regions and normal breast tissue in their evolutionary processes. **b,** Quantitative comparisons of overall CPM, overall evolutionary gradient (CPM slope), overall evolutionary flow (flow magnitude) and front-cell flow magnitude between CMZ and BMZ. Statistical comparisons indicate significant differences between CMZ and BMZ across all these metrics (*p <* 0.0001). **c,** Spatial subclone analysis and 3D reconstruction. Cell-level subclones: spatial distribution of subclones based on CPM isobands (left), spatial mapping in H&E tissue sections (middle) and box plots of subclone gradient distributions (right). CMZ-level subclones: spatial clustering of CMZ-level subclones based on 3F fingerprints. 3D topological analysis: CMZ 3D growth trajectories where nodes represent centroids of tumour foci with node size proportional to lesion area and edges indicating continuity between adjacent slices. 3D reconstruction: CMZ structures. **d,** Macro-CPM correlates with front load, and pairwise evolutionary fitness index (EFI) comparisons between cancer microzones indicate localized clonal competition.

**Extended Data Fig. 5.**
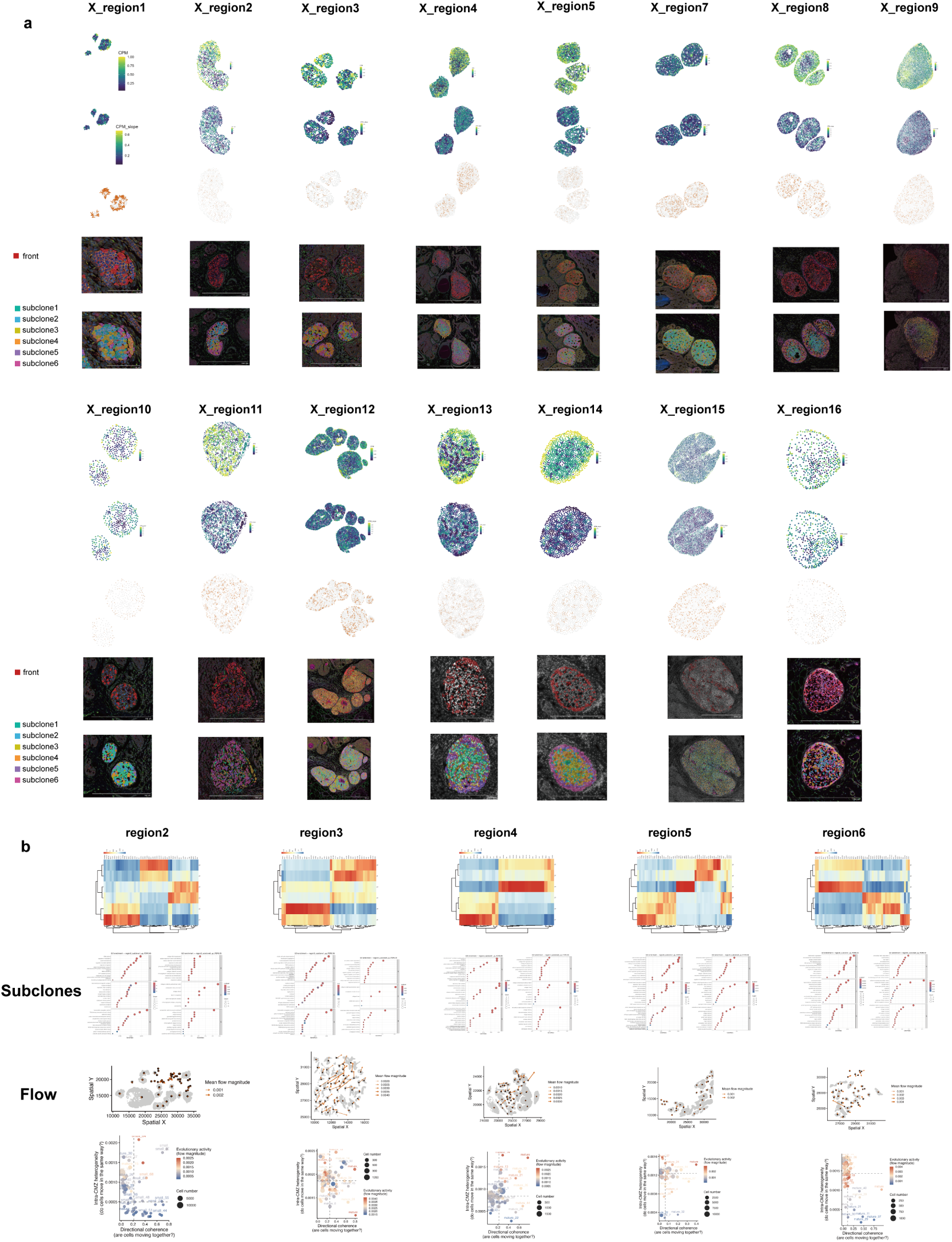
Independent Xenium validation of 3F dynamics, subclonal functional stratification and CMZ evolutionary flow dynamics. **a,** Independent validation of 3F dynamics in single-cell resolution Xenium tumour regions (X region1–16; X region6 shown in the main figure). For each region, the top three rows show the 3F spatial maps (CPM, CPM slope and flow), and the bottom two rows show the corresponding front and subclone (subclone1–6) assignments mapped onto the tissue in Xenium Explorer. **b,** Subclonal functional architecture and spatial organization of CMZ evolutionary flow. Left panels show heatmaps of average marker gene expression and functional enrichment across subclones defined along CPM in multiple spatial regions, illustrating region-specific reorganization of stage-associated transcriptional programs. Right panels depict CMZ-level evolutionary dynamics, including spatial mapping of CMZ evolutionary activity quantified by median flow magnitude; a CMZ evolutionary phase diagram defined by directional coherence (x axis) and intra-CMZ heterogeneity (y axis), with point size indicating CMZ cell number and colour representing evolutionary activity; and spatial projection of mean CMZ evolutionary flow vectors, where arrow direction indicates the dominant evolutionary tendency and arrow width is proportional to mean flow magnitude.

**Extended Data Fig. 6.**
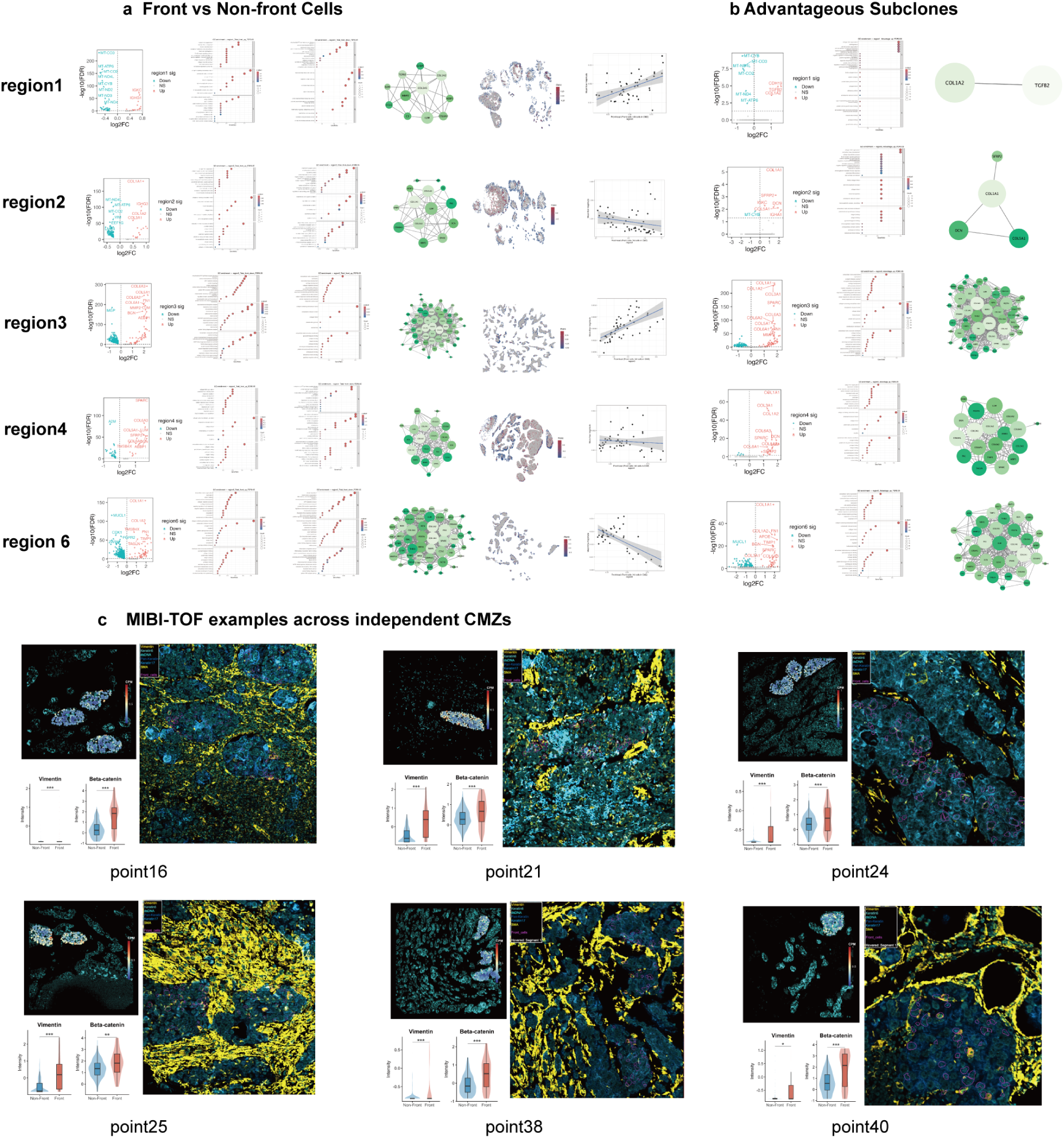
Molecular characterization of the tumour front and advantageous subclones. **a,** Biological characterization of front and non-front cells. Volcano plots of differential gene expression highlight significantly upregulated and downregulated genes. GO enrichment analysis of differentially expressed genes (DEGs) illustrates functional features of front cells, including extracellular matrix (ECM) remodelling and collagen metabolism. PPI network analysis reveals molecular interactions and ECM-related modules. The spatial distribution of the Front Molecular Heterogeneity Index (FMHI) indicates functional heterogeneity of tumour fronts across regions. Regression scatter plots show the relationship between CMZ front load and evolutionary flow magnitude. **b,** Molecular features and functional bias of advantageous subclones. Volcano plots show DEGs in advantageous subclones. GO enrichment of upregulated genes highlights extracellular matrix and collagen metabolism. PPI network analysis illustrates interaction networks associated with advantageous subclones and their central role in ECM remodelling. **c,** MIBI-TOF examples showing the “inner-low, outer-high” CPM-pp topology and peripherally localized front cells (magenta) across independent CMZs. Per-point violin/box plots confirm significantly elevated Vimentin and Beta-catenin in front versus non-front cells, consistent with EMT–Wnt activation at the invasive front (Wilcoxon rank-sum test; \*\*\**p <* 0.001).

**Extended Data Fig. 7.**
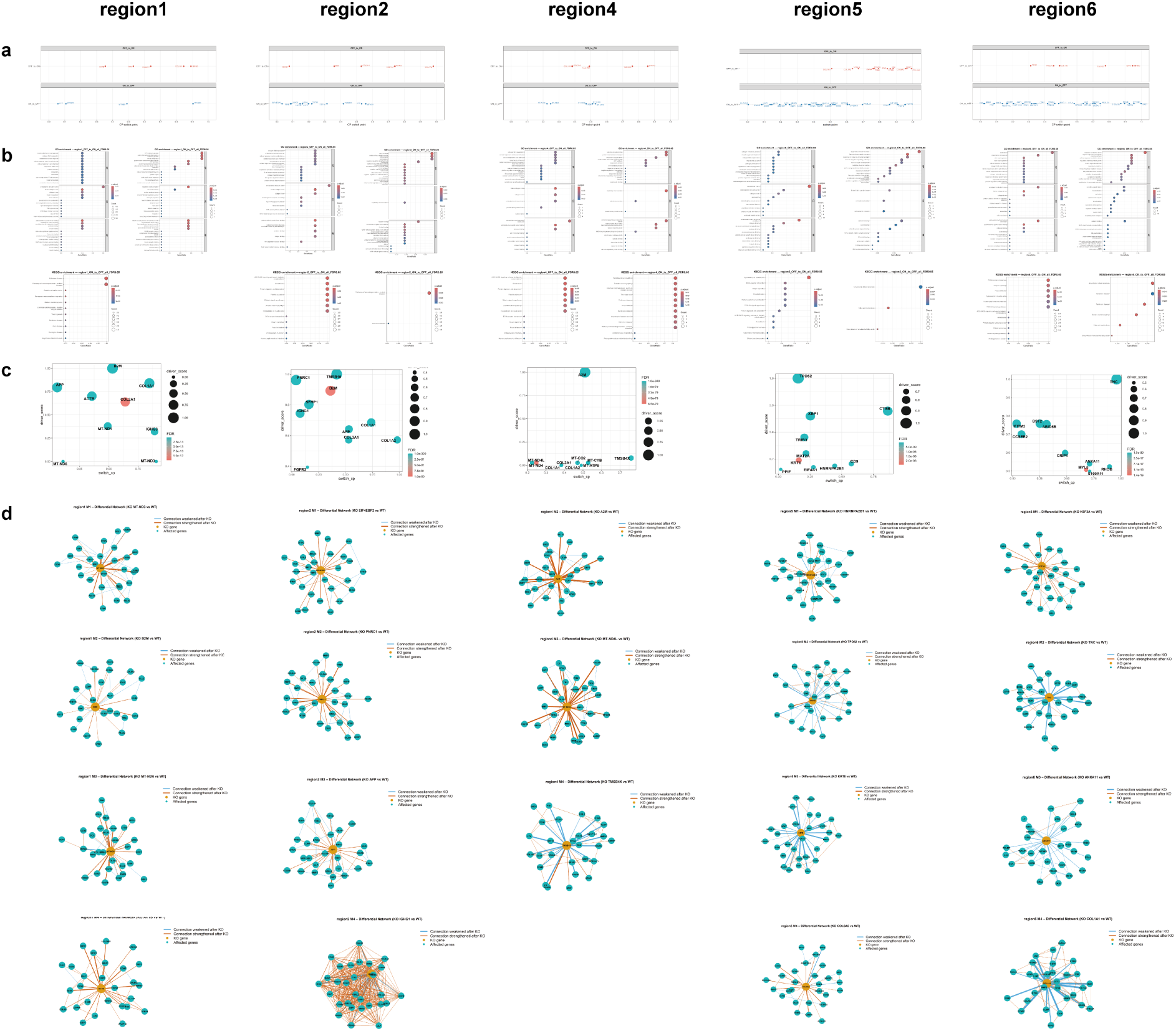
CPM-GeneSwitch dynamics, functional enrichment and driver-network analyses across spatial regions. **a,** CPM-GeneSwitch overview in each region showing the distribution of significant switch positions along the CPM axis stratified by switching direction (OFF→ON versus ON→OFF), enabling comparison of stage-specific transcriptional switching across spatial regions. **b,** Functional enrichment across regions: GO and KEGG dotplots for significant switch genes from OFF→ON and ON→OFF sets, visualizing GeneRatio/Count and statistical significance (FDR) to assess shared versus region-specific pathway activation or repression during evolutionary progression. **c,** Driver-gene bubble plots per region: switch cpm (x-axis) versus driver score (y-axis), with dot size indicating driver strength and colour indicating FDR, summarizing the stage-resolved distribution of network-informed candidate drivers along pseudotime. **d,** scTenifoldNet differential regulatory networks following in silico KO across regions, illustrating KO-induced strengthening/weakening of regulatory edges and affected gene neighbourhoods at the module/driver level, supporting reproducibility of driver-module perturbation patterns across spatial contexts.

**Extended Data Fig. 8.**
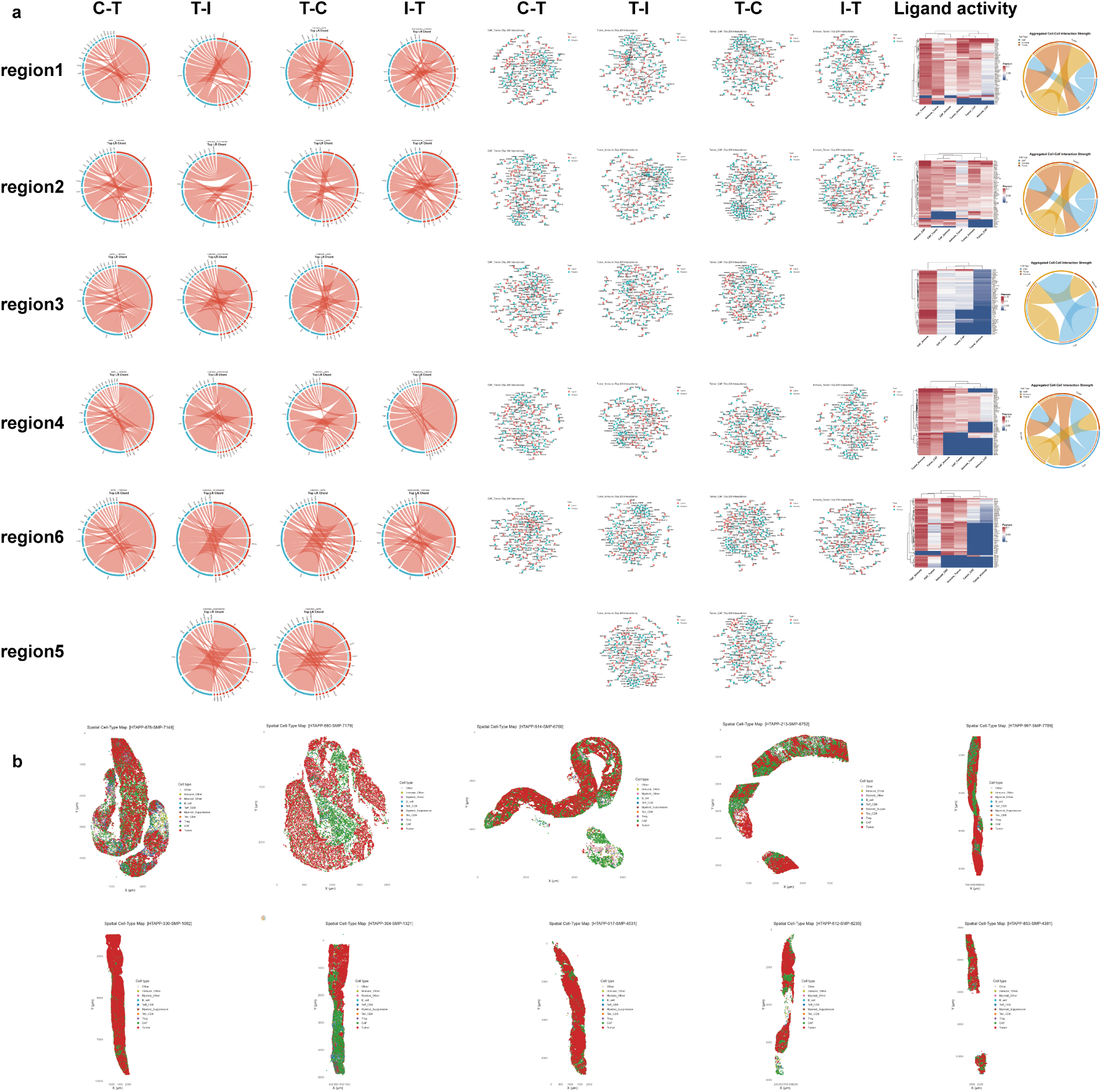
Spatial cell–cell communication networks and CODEX-based cell-type architecture across tumour regions. **a,** Comprehensive profiling of cell–cell communication and ligand–receptor network architecture across multiple regions (region1–region6). For each region, directional chord diagrams (C–T, T–I, T–C, I–T) depict inferred ligand–receptor interactions among CAF, tumour and immune populations, with ribbon width proportional to summed interaction weights, indicating relative signalling intensity. Corresponding network topology plots illustrate the structural organization of principal ligand and receptor nodes, highlighting region-specific arrangements of interaction hubs. Ligand activity heatmaps based on Pearson correlation evaluate the predicted regulatory potential of key ligands on target gene programs in receiver cells, revealing differences in activity across spatial contexts. Aggregated interaction-strength chord diagrams summarize overall signalling output and input among major cell types within each region, defining the communication architecture characteristic of individual spatial areas. **b,** Spatial cell-type maps of CODEX-profiled tumour samples. Single-cell spatial distributions of tumour cells (*KRT19* ^+^), CAFs (*FAP*/*VIM* ^+^) and immune subsets were reconstructed by marker-based gating from CODEX protein intensities, revealing recurrent juxtaposition of tumour nests and CAF-rich stroma that defines the CMZs used for downstream analysis.

**Extended Data Fig. 9.**
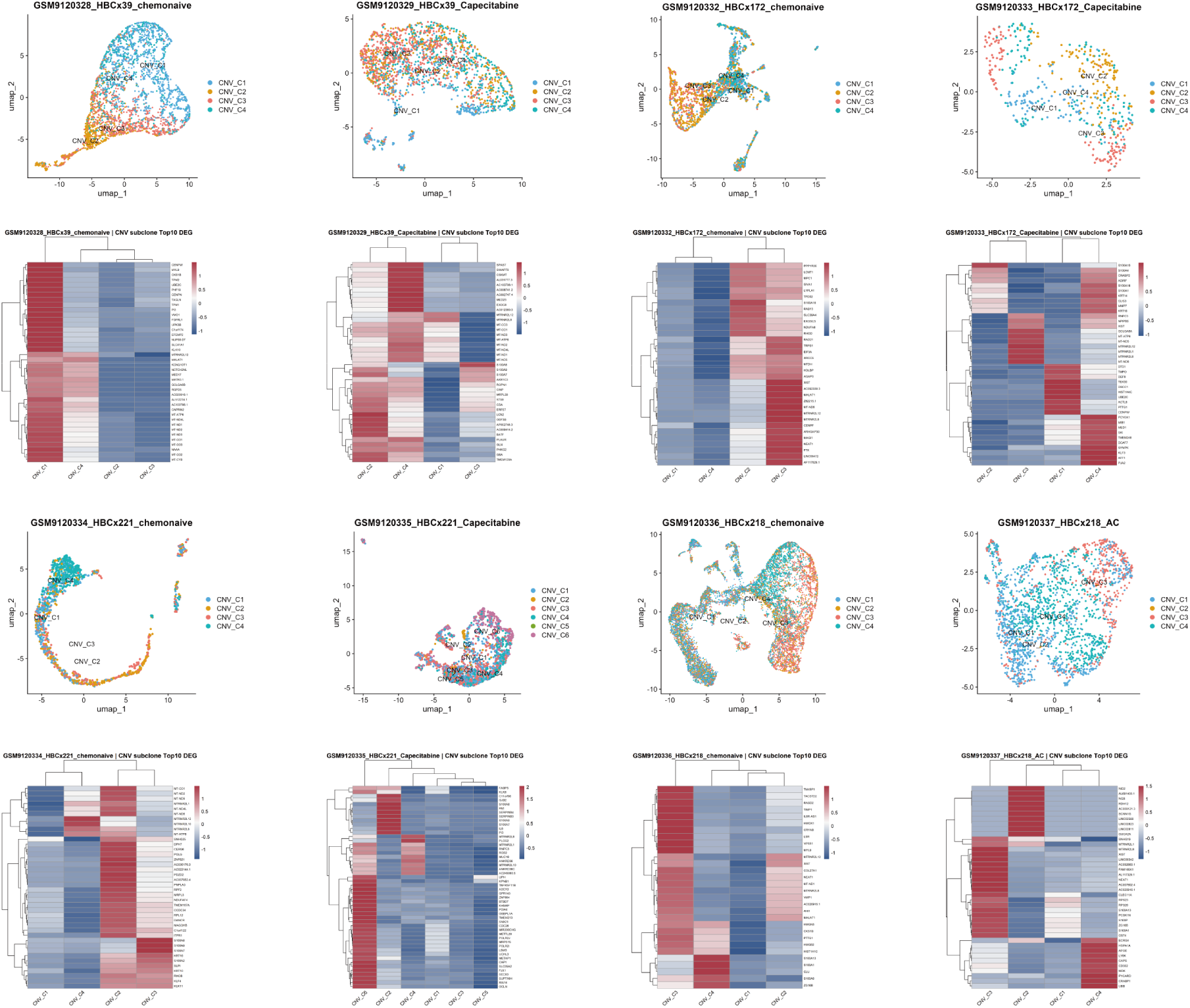
Therapeutic remodelling of tumour subclones. Transcriptional changes of CNV-defined subclones before and after therapy across multiple PDX models. UMAP embeddings illustrate the distribution of distinct CNV subclones in chemonaive and treated samples. Corresponding heatmaps of the Top10 differentially expressed genes highlight subclone-specific transcriptional programs and their remodelling following therapeutic intervention. Consistent patterns across models indicate reproducible therapy-associated subclonal reprogramming.

**Extended Data Fig. 10.**
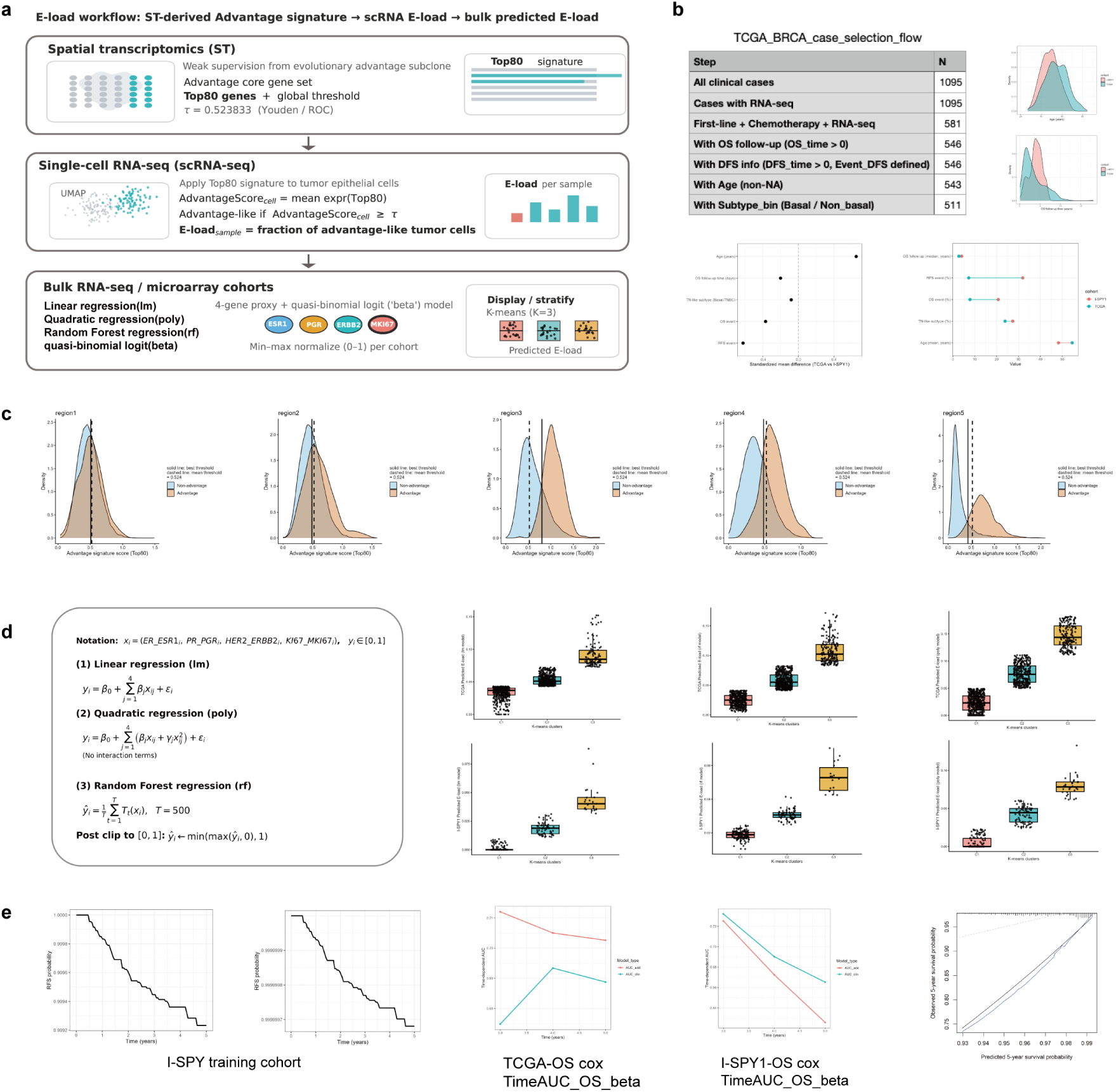
Workflow, cohort characteristics and model robustness analyses of E-load. **a,** Cross-scale construction of E-load. Evolutionary advantage subclones from spatial transcriptomics defined the Top80 Advantage core gene set with a unified cutoff. The signature was projected to scRNA-seq to compute sample-level E-load, and a four-gene regression model (ESR1, PGR, ERBB2, MKI67) predicted E-load in bulk cohorts. **b,** Clinical cohort selection and baseline comparison. TCGA-BRCA case selection, age distribution, OS follow-up and key clinical characteristics are shown for TCGA and I-SPY1. The cohorts are independent in population source and transcriptomic platform but comparable in age and molecular subtype composition. **c,** Regional distribution of Top80 AdvantageScore and threshold consistency. Density distributions for advantage and non-advantage cells are shown across region1–region5. Solid lines denote region-specific ROC–Youden optimal cutoffs; dashed lines denote the cross-region mean unified threshold (0.524). Despite regional variability, optimal thresholds cluster around the unified cutoff, supporting a global threshold. **d,** Comparison of regression models. Linear, polynomial and random forest models are shown with predicted E-load stratification. **e,** Prognostic performance and validation. In I-SPY1, higher E-load was associated with increased recurrence risk (RFS curves). In TCGA, adding E-load to Cox models (Clinical + E-load) improved 3–5 year time-dependent AUC versus clinical variables alone; similar trends were observed in I-SPY1, supporting cross-cohort transferability. The TCGA 5-year OS calibration plot shows strong agreement between predicted and observed survival, with the optimism-corrected curve close to the ideal reference line.

## Supplementary information

**Supplementary Figures 1–10 —** Provided as separate files; corresponding legends are included at the end of the manuscript.

**Supplementary Table 1 —** Multi-platform sample inventory: Visium HD clinical data, imaging panels (Xenium, MIBI-TOF, CODEX), and DepMap cell lines.

**Supplementary Table 2 —** Advantage-subclone signature and evolutionary driver-module (M1–M4) gene sets.

**Supplementary Table 3 —** Single-cell copy number variation subclonal assignments across paired PDX models.

**Supplementary Table 4 —** Sample-level quantification of E-load in the GSE161529 scRNA-seq cohort.

**Supplementary Table 5 —** Predicted E-load across bulk RNA-seq clinical cohorts.

**Supplementary Table 6 —** Clinical characteristics and survival outcomes of the TCGA-BRCA cohort.

**Supplementary Table 7 —** Clinical characteristics, treatment response, and survival outcomes of the I-SPY1 cohort.

## Notes

https://github.com/Euphonni/spatiotemporal-tumor-evolution

https://ngdc.cncb.ac.cn/gsa-human/browse/HRA017537

### Competing Interest Statement

The authors have declared no competing interest.

## References

[1] Pienta, K. J., Goodin, P. L. & Amend, S. R. Defeating lethal cancer: Interrupting the ecologic and evolutionary basis of death from malignancy. CA: A Cancer Journal for Clinicians 75, 183–202 (2025).

[2] Zuckerkandl, E. Molecular disease, evolution, and genic heterogeneity. Horizons in biochemistry 189–225 (1962).

[3] Nowell, P. C. The clonal evolution of tumor cell populations: Acquired genetic lability permits stepwise selection of variant sublines and underlies tumor progression. Science 194, 23–28 (1976).

[4] Tang, F. et al. mrna-seq whole-transcriptome analysis of a single cell. Nature methods 6, 377–382 (2009).

[5] Vickovic, S. et al. High-definition spatial transcriptomics for in situ tissue profiling. Nature methods 16, 987–990 (2019).

[6] Roth, A. et al. Pyclone: statistical inference of clonal population structure in cancer. Nature methods 11, 396–398 (2014).

[7] Deshwar, A. G. et al. Phylowgs: reconstructing subclonal composition and evolution from whole-genome sequencing of tumors. Genome biology 16, 35 (2015).

[8] Malikic, S., McPherson, A. W., Donmez, N. & Sahinalp, C. S. Clonality inference in multiple tumor samples using phylogeny. Bioinformatics 31, 1349–1356 (2015).

[9] Wang, K. et al. Single-cell phylodynamic inference of stem cell differentiation and tumor evolution. Cell Systems 16 (2025).

[10] Ma, C. et al. Inferring allele-specific copy number aberrations and tumor phylogeography from spatially resolved transcriptomics. Nature methods 21, 2239–2247 (2024).

[11] Gao, T. et al. Haplotype-aware analysis of somatic copy number variations from single-cell transcriptomes. Nature Biotechnology 41, 417–426 (2023).

[12] Elyanow, R., Zeira, R., Land, M. & Raphael, B. J. Starch: copy number and clone inference from spatial transcriptomics data. Physical biology 18, 035001 (2021).

[13] Gabbutt, C. et al. Fluctuating dna methylation tracks cancer evolution at clinical scale. Nature 645, 764–773 (2025).

[14] Pawlik, P. et al. Clone copy number diversity is linked to survival in lung cancer. Nature 646, 190–197 (2025).

[15] Salcedo, A. et al. Crowd-sourced benchmarking of single-sample tumor subclonal reconstruction. Nature biotechnology 43, 581–592 (2025).

[16] Nolan, E., Lindeman, G. J. & Visvader, J. E. Deciphering breast cancer: from biology to the clinic. Cell 186, 1708–1728 (2023).

[17] Laplane, L. & Maley, C. C. The evolutionary theory of cancer: challenges and potential solutions. Nature Reviews Cancer 24, 718–733 (2024).

[18] Alexandrov, L. B. et al. The repertoire of mutational signatures in human cancer. Nature 578, 94–101 (2020).

[19] George, J. et al. Evolutionary trajectories of small cell lung cancer under therapy. Nature 627, 880–889 (2024).

[20] Lomakin, A. et al. Spatial genomics maps the structure, nature and evolution of cancer clones. Nature 611, 594–602 (2022).

[21] Mo, C.-K. et al. Tumour evolution and microenvironment interactions in 2d and 3d space. Nature 634, 1178–1186 (2024).

[22] Wang, K. et al. Coalescing single-cell genomes and transcriptomes to decode breast cancer progression. Cell 188, 6355–6369 (2025).

[23] Househam, J. et al. Phenotypic plasticity and genetic control in colorectal cancer evolution. Nature 611, 744–753 (2022).

[24] Zhao, T. et al. Spatial genomics enables multi-modal study of clonal heterogeneity in tissues. Nature 601, 85–91 (2022).

[25] Ren, H., Walker, B. L., Cang, Z. & Nie, Q. Identifying multicellular spatiotemporal organization of cells with spaceflow. Nature communications 13, 4076 (2022).

[26] Long, W., Liu, T., Xue, L. & Zhao, H. spvelo: Rna velocity inference for multi-batch spatial transcriptomics data. Genome biology 26, 239 (2025).

[27] Pham, D. et al. stlearn: integrating spatial location, tissue morphology and gene expression to find cell types, cell-cell interactions and spatial trajectories within undissociated tissues. biorxiv 2020–05 (2020).

[28] Pastushenko, I. et al. Identification of the tumour transition states occurring during emt. Nature 556, 463–468 (2018).

[29] Cox, T. R. The matrix in cancer. Nature Reviews Cancer 21, 217–238 (2021).

[30] Greaves, M. & Maley, C. C. Clonal evolution in cancer. Nature 481, 306–313 (2012).

[31] Maley, C. C. et al. Classifying the evolutionary and ecological features of neoplasms. Nature Reviews Cancer 17, 605–619 (2017).

[32] Lewinsohn, M. A., Bedford, T., Müller, N. F. & Feder, A. F. State-dependent evolutionary models reveal modes of solid tumour growth. Nature Ecology & Evolution 7, 581–596 (2023).

[33] Gong, D., Arbesfeld-Qiu, J. M., Perrault, E., Bae, J. W. & Hwang, W. L. Spatial oncology: Translating contextual biology to the clinic. Cancer Cell 42, 1653–1675 (2024).

[34] Noble, R. et al. Spatial structure governs the mode of tumour evolution. Nature ecology & evolution 6, 207–217 (2022).

[35] Chitty, J. L. & Cox, T. R. The extracellular matrix in cancer: from understanding to targeting. Trends in Cancer (2025).

[36] Pavlova, N. N. & Thompson, C. B. The emerging hallmarks of cancer metabolism. Cell metabolism 23, 27–47 (2016).

[37] Binnewies, M. et al. Understanding the tumor immune microenvironment (time) for effective therapy. Nature medicine 24, 541–550 (2018).

[38] Masuda, H. Cancer-associated fibroblasts in cancer drug resistance and cancer progression: a review. Cell Death Discovery 11, 341 (2025).

[39] Kim, C. et al. Chemoresistance evolution in triple-negative breast cancer delineated by single-cell sequencing. Cell 173, 879–893 (2018).

[40] Network, C. G. A. Comprehensive molecular portraits of human breast tumours. Nature 490, 61–70 (2012).

[41] Magbanua, M. J. M. et al. Serial expression analysis of breast tumors during neoadjuvant chemotherapy reveals changes in cell cycle and immune pathways associated with recurrence and response. Breast Cancer Research 17, 73 (2015).

[42] McGranahan, N. & Swanton, C. Clonal heterogeneity and tumor evolution: past, present, and the future. Cell 168, 613–628 (2017).

[43] Frankell, A. M. et al. The evolution of lung cancer and impact of subclonal selection in tracerx. Nature 616, 525–533 (2023).

[44] Marx, V. Method of the year: spatially resolved transcriptomics. Nature methods 18, 9–14 (2021).

[45] Hanahan, D. & Weinberg, R. A. Hallmarks of cancer: the next generation. cell 144, 646–674 (2011).

[46] Marusyk, A., Janiszewska, M. & Polyak, K. Intratumor heterogeneity: the rosetta stone of therapy resistance. Cancer cell 37, 471–484 (2020).

[47] Zhang, J., Cunningham, J. J., Brown, J. S. & Gatenby, R. A. Integrating evolutionary dynamics into treatment of metastatic castrate-resistant prostate cancer. Nature communications 8, 1816 (2017).

[48] Acar, A. et al. Exploiting evolutionary steering to induce collateral drug sensitivity in cancer. Nature communications 11, 1923 (2020).

[49] Thermo Fisher Scientific. RecoverAll™ Total Nucleic Acid Isolation Kit for FFPE: User Guide. Thermo Fisher Scientific, Waltham, MA, USA (2019). Protocol handbook.

[50] Thermo Fisher Scientific. NanoDrop Spectrophotometers User Guide. Thermo Fisher Scientific, Waltham, MA, USA (2018). UV spectrophotometric quantification of nucleic acids.

[51] Agilent Technologies. Agilent RNA 6000 Nano Kit Guide. Agilent Technologies, Santa Clara, CA, USA (2016). RNA quality assessment using microfluidics-based electrophoresis.

[52] Agilent Technologies. Agilent 2100 Bioanalyzer System User Guide. Agilent Technologies, Santa Clara, CA, USA (2013).

[53] 10x Genomics. Visium HD FFPE Tissue Preparation Handbook. 10x Genomics, Pleasanton, CA, USA, rev. a edn. (2024). Protocol handbook.

[54] 10x Genomics. User Guide: Visium HD Spatial Gene Expression Reagent Kits. 10x Genomics, Pleasanton, CA, USA, rev. a edn. (2024). User guide.

[55] Stuart, T. et al. Comprehensive integration of single-cell data. cell 177, 1888–1902 (2019).

[56] Hafemeister, C. & Satija, R. Normalization and variance stabilization of single-cell rna-seq data using regularized negative binomial regression. Genome biology 20, 296 (2019).

[57] Wu, S. Z. et al. A single-cell and spatially resolved atlas of human breast cancers. Nature genetics 53, 1334–1347 (2021).

[58] Cable, D. M. et al. Robust decomposition of cell type mixtures in spatial transcriptomics. Nature biotechnology 40, 517–526 (2022).

[59] Qiu, X. et al. Reversed graph embedding resolves complex single-cell trajectories. Nature methods 14, 979–982 (2017).

[60] Satija, R., Farrell, J. A., Gennert, D., Schier, A. F. & Regev, A. Spatial reconstruction of single-cell gene expression data. Nature biotechnology 33, 495–502 (2015).

[61] Wang, H. & Song, M. Ckmeans. 1d. dp: optimal k-means clustering in one dimension by dynamic programming. The R journal 3, 29 (2011).

[62] Blondel, V. D., Guillaume, J.-L., Lambiotte, R. & Lefebvre, E. Fast unfolding of communities in large networks. Journal of statistical mechanics: theory and experiment 2008, P10008 (2008).

[63] Evangelidis, G. D. & Psarakis, E. Z. Parametric image alignment using enhanced correlation coefficient maximization. IEEE transactions on pattern analysis and machine intelligence 30, 1858–1865 (2008).

[64] Bankhead, P. et al. Qupath: Open source software for digital pathology image analysis. Scientific reports 7, 1–7 (2017).

[65] Lorensen, W. E. & Cline, H. E. Marching cubes: A high resolution 3d surface construction algorithm. In Seminal graphics: pioneering efforts that shaped the field, 347–353 (1998).

[66] Ashburner, M. et al. Gene ontology: tool for the unification of biology. Nature genetics 25, 25–29 (2000).

[67] Kanehisa, M. & Goto, S. Kegg: kyoto encyclopedia of genes and genomes. Nucleic acids research 28, 27–30 (2000).

[68] Szklarczyk, D. et al. String v11: protein–protein association networks with increased coverage, supporting functional discovery in genome-wide experimental datasets. Nucleic acids research 47, D607–D613 (2019).

[69] Scarselli, F., Gori, M., Tsoi, A. C., Hagenbuchner, M. & Monfardini, G. The graph neural network model. IEEE transactions on neural networks 20, 61–80 (2008).

[70] Kipf, T. Semi-supervised classification with graph convolutional networks. arXiv preprint arXiv:1609.02907 (2016).

[71] Osorio, D., Zhong, Y., Li, G., Huang, J. Z. & Cai, J. J. sctenifoldnet: a machine learning workflow for constructing and comparing transcriptome-wide gene regulatory networks from single-cell data. Patterns 1 (2020).

[72] Browaeys, R., Saelens, W. & Saeys, Y. Nichenet: modeling intercellular communication by linking ligands to target genes. Nature methods 17, 159–162 (2020).

[73] Iglesia, M. D. et al. Differential chromatin accessibility and transcriptional dynamics define breast cancer subtypes and their lineages. Nature Cancer 5, 1713–1736 (2024).

[74] Aran, D. et al. Reference-based analysis of lung single-cell sequencing reveals a transitional profibrotic macrophage. Nature immunology 20, 163–172 (2019).

[75] Kaplan, E. L. & Meier, P. Nonparametric estimation from incomplete observations. Journal of the American statistical association 53, 457–481 (1958).

[76] Cox, D. R. Analysis of survival data (Chapman and Hall/CRC, 2018).

[77] Harrell, F. E., Califf, R. M., Pryor, D. B., Lee, K. L. & Rosati, R. A. Evaluating the yield of medical tests. Jama 247, 2543–2546 (1982).

[78] Akaike, H. A new look at the statistical model identification. IEEE transactions on automatic control 19, 716–723 (2003).

[79] Belsley, D. A., Kuh, E. & Welsch, R. E. Regression diagnostics: Identifying influential data and sources of collinearity (John Wiley & Sons, 2005).

[80] Blanche, P., Dartigues, J.-F. & Jacqmin-Gadda, H. Estimating and comparing time-dependent areas under receiver operating characteristic curves for censored event times with competing risks. Statistics in medicine 32, 5381–5397 (2013).

[81] Vickers, A. J. & Elkin, E. B. Decision curve analysis: a novel method for evaluating prediction models. Medical Decision Making 26, 565–574 (2006).

[82] Efron, B. The bootstrap and modern statistics. Journal of the American Statistical Association 95, 1293–1296 (2000).

